# Clinical development of gene edited tacrolimus-resistant Treg (FKBP12^KO^-Treg) to enable simultaneous immunosuppression and support of immune regulation

**DOI:** 10.64898/2025.12.10.693159

**Authors:** Ghazaleh Zarrinrad, Lisa-Marie Burkhardt, Dimitrios Laurin Wagner, Stephan Schlickeiser, Maik Stein, Désirée Jacqueline Wendering, Lukas Ehlen, Gavin L. Kurgan, Pawel Durek, Frederik Heinrich, Anamika Giri, Kristy Ou, Henrike Hoffmann, Sandra Muench, Insa Stuewe, Sven Dolling, Oliver McCallion, Jaspal Kaeda, Jonas Kath, Sarah Schulenberg, Abdolreza Nazari, Lena Peter, Samira Picht, Rolf Turk, Garrett Rettig, Morgan Sturgeon, Thomas L. Osborne, Ashley Jacobi, Rebecca Friedrich, Masako Monika Kaufmann, Julia Klermund, Simon Fink, Markus F. Templin, Andy Roemhild, Daniel Kaiser, Oliver Klein, Toni Cathomen, Mir-Farzin Mashreghi, Fadi Issa, Julia K. Polánsky, Hans-Dieter Volk, Michael Schmueck-Henneresse, Petra Reinke, Leila Amini

## Abstract

**Background:** Unwanted immune responses play a central role in the pathogenesis of solid organ allograft rejection. These are managed by life-long immunosuppression with considerable burden for the patient and society. Adoptive therapy with regulatory T-cells (Treg) is a promising approach to restore sustainable immune balance and avoid long-term adverse effects of immunosuppression. While Treg effectively inhibit activation of unwanted immune responses, they are less effective in controlling pre-existing/activated memory effector T-cells (Teff). Thus, co-administration of Treg with immunosuppressants is required to achieve a sustainable organ acceptance. Calcineurin inhibitors (CNI) are powerful in controlling de novo generated and preformed Teff. However, CNI also dampen Treg immunoregulatory function. Thus, we hypothesize improved results of adoptive Treg therapy in immunosuppressed patients applying tacrolimus-resistant Treg. **Methods:** While retaining CNI modulation of Teff with tacrolimus, we knocked-out FKBP12 in Treg (FKBP12^KO^-Treg) by gene-editing using ribonucleoprotein-based CRISPR/Cas9 technology to generate tacrolimus-resistant Treg and characterised them using flow cytometry, functional assays and in-depth phenotyping. **Results:** This detailed in vitro analysis showed FKBP12^KO^-Treg were comparable to non-gene edited Treg and impervious to tacrolimus while maintaining immunoregulatory function and sensitivity to alternative CNIs raising no safety concerns. Furthermore, we aligned our methodology to achieve GMP compliance laying the basis for a manufacturing license in preparation of a clinical trial. **Conclusion:** Based on the presented preclinical dataset implying safety and efficacy of FKBP12^KO^-Treg, we are now seeking to undertake a proof-of-concept clinical trial to evaluate the co-administration of FKBP12^KO^-Treg and tacrolimus to enhance the management of living donor kidney transplant recipients.

## Introduction

Allo-transplantation represents a clearly defined “starting point” of an undesired immune reactivity. Currently, patients are managed with a cocktail of immunosuppressants, including calcineurin inhibitors (CNI), *i.e.* tacrolimus (FK506) or cyclosporine A (CsA), associated with significant adverse effects^1–5^. To overcome these challenges supplanting or weaning the standard immunosuppression with adoptive regulatory T-cell (Treg) therapy is envisioned. Stable Treg are characterized by constitutive high level expression of the interleukin-2 (IL-2) receptor α-chain, CD25, orchestrated by the transcription factor FOXP3 ^6^ and low CD127 ^7^. Immunomodulatory Treg are different to the activated conventional T-cells (Tcon) by the demethylated promoter region of the FOXP3 gene, the Treg-specific demethylation region (TSDR) ^8–12^. A clinical phase 1/2a study, conducted at our institute, reported first authoritative proof-of-concept of autologous Treg therapy facilitating the reduction of triple immunosuppression to low-dose tacrolimus monotherapy in the majority of living donor kidney transplant recipients ^13,14^. But because CNI therapy not only diminishes preformed alloreactive Teffs ^15^, but also impedes Treg ^16^, we report detailed characterization of a tacrolimus-resistant FKBP12^KO^-Treg product (FKBP12^KO^-Treg) ^17,18^ as a promising development for clinical application in transplantation and beyond.

The FKBP12^KO^-Treg products (FKBP12^KO^-Treg) were deeply characterized and compared to unedited Treg products, herein called Treg^WT^, produced under the same conditions. The assays demonstrated phenotypic and epigenetic stability and comparable properties of FKBP12^KO^-Treg and Treg^WT^. In depth profiling of transcriptomes and proteomes revealed a beneficial marker profile of FKBP12^KO^-Treg compared to Treg^WT^, especially in presence of tacrolimus. Safety assays reduced concerns about FKBP12^KO^-Treg. Furthermore, we developed a GMP compliant production process for FKBP12^KO^-Treg. Thus, we now aim for proof-of-concept of increased efficacy of FKBP12^KO^-Treg in a first-in-human/class clinical trial.

## Materials and Methods

All information on antibodies is attached in suppl. Table 1 and information on materials in suppl. Table. 2.

### Blood sampling and ethics

We recruited a total of 15 healthy volunteers who gave a written declaration of consent. This study was approved by the Ethics Committee of Charité-Universitaetsmedizin Berlin as EA4/091/19 and EA1/052/22.

### Isolation and culture of Treg

CD4^+^ T-cells were isolated from peripheral whole blood either by a research-grade or a GMP protocol; i.e. EasySep™ Human CD4^+^ T-cell Enrichment Kit (STEMCELL Technologies) and CliniMACS Plus device (Miltenyi Biotec), respectively. CD4^+^ cells subsequently underwent fluorescently activated cell sorting (FACS) using MACSQuant-Tyto (Suppl. Table 1, Miltenyi Biotec). Tregs were thereafter cultured with Treg medium supplemented with Rapamycin at a concentration of 100nM/mL (Suppl. Table 2). Tregs were subjected to expansion over the first 7 days with anti-CD3/CD28-MACSiBeads at 4:1 ratio and following 14 days at 1:1 ratio replenished every other day when cultivated in 24 well plates (research grade, pre-GMP). For GMP grade expansion, Tregs were cultured in G-Rex bioreactors (G-Rex 10 and 100, Wilson Wolf Manufacturing). An aliquot of the negative fraction of the FACSort, *i.e.* non-Tregs (Tcon) served as control, and was expanded using identical conditions.

### FKBP12 knock-out in Treg

On day 7, the Tregs were electroporated with ribonucleoprotein (RNP) complexes using the Amaxa-P3-primary-cell-4D-Nucleofector-X-Kit-L and the Amaxa-Nucleofector-4D (Lonza); after pre-complexing 10μg Alt-R™ S.p. (HiFi) Cas9 Nuclease V3^19^ (Integrated DNA Technologies) with 15μg 2O’-methyl-3’phosphothioate-modified sgRNA^17^ (Synthego).

### Phenotypic and functional cytokine profiling

Phenotypic and functional characterization was performed by flow cytometry. After resting overnight in 10% FCS containing RPMI, 1×10^6^ cells were stimulated with phorbol-12-myristate-13-acetate (PMA) at 10ng/mL and Ionomycin at 2,5μg/mL at 37°C for 6h. DMSO was added to the unstimulated control. Afterwards, cells were labelled with antibodies (Suppl. Table 1). For intracellular staining the cells were fixed and permeabilized with the FoxP3/Transcription Factor Staining Buffer kit and stained for activation markers (Suppl. Table 1). Cells were measured using flow cytometers (i.e. LSR II Fortessa/Cytoflex LX) and analyzed using FlowJo. Following 21 days of expansion the pSTAT5 levels were assessed, pre-stained cells were seeded at range of concentrations (0.5 x 1060.4 x 10 x 60.5 x 106 cells per well), stimulated at varying IL-2 concentrations (0, 1, 10, 100, 1000 U/mL) for 30 minutes at 37°C.

### T-cell proliferation suppression assay

After resting overnight in 10% FCS containing RPMI, Treg suppressive capacity was analyzed on freshly isolated, autologous CD3^+^ T-cells (Human T cell enrichment cocktail, STEMCELL Technologies), namely responder T-cells. Responder T-cells (Tresp) were stained with 10 μM carboxyfluorescein diacetate succinimidyl ester (CFSE; Sigma-Aldrich). Following that, autologous Treg^WT^ and FKBP12^KO^Treg were co-cultured at various Treg/Tresp ratios under stimulation with anti-CD3/CD28 microbeads (Treg Suppression Inspector, Miltenyi Biotec) and according to an already published co-culture protocol^20^. The percentage of Tresp/T-cell proliferation, detected by the level of CFSE dilution, was calculated in the presence and absence of Treg. Extracellular flow cytometric staining including viability assessment as well as CD3, CD4 and CD8 was implemented, measured using the LSR II Fortessa flow cytometer and analyzed using FlowJo.

### FKBP12 knockout efficiency analysis and validation using digital droplet PCR

Editing efficiency was determined, as previously described, using the Inference-of-CRISPR-Edits (ICE) algorithm (Synthego Corporation)^17,18,21^.

During GMP process development, the following digital droplet polymerase chain reaction (ddPCR) assay was established. On day 0.5×10^6^ cells of the negative fraction of the Tyto-sort were cryo-stored to elaborate on KO efficacy. In addition, 5×10^6^ cells of the FKBP12^KO^-Treg were taken for DNA isolation of both samples using the QIAmp DNA kit (QIAGEN) according to the manufacturer’s protocol on day 17. Following that, DNA concentration is measured using the Qubit Assay system (ThermoFisher) and samples undergo ddPCR for KO efficacy detection. KO efficacy data from biological samples is correlated to a standard curve generated by KO controls with 50, 75, and 100% KO rate (deviation of +/− 10 % allowed). The following sequences for ddPCR probes were used: RTP28 (5’- /5HEX/CAC CTT CCC CAA GC+G CGG /3IABkFQ/ −3’), RT-P22 (5’ /56-FAM/AGC CGC CGC /ZEN/GCG CCA CTA CT/ 3IABkFQ/ −3’) in combination with the following primers: Primer RT-25 (5’-ATG GGA GTG CAG GTG GAA ACC ATC −3’) and primer RT-26 (5’-CGC TGG GCC CCC GAC TCA −3’).

### Methylation profiling

The TSDR methylation status was assessed using the Infinium-Methylation-EPIC-Kit. 200–250ng of DNA was subjected to bisulfite conversion using EZ-DNA-Methylation-Gold-Kit. Bead Chips were imaged on Illumina’s Microarray Scanner iScan. Prior to sequencing by Genewiz/Azenta, amplicons were purified with QIAquick-PCR-Purification-Kit and normalized to 20ng/μl. Methods applied as per manufacturer’s instruction.

### T-cell receptor (TCR) sequencing

DNA was extracted using All-Prep-DNA/RNA-Kit (Qiagen) and subjected to TCR β sequencing (Adaptive Biotechnologies) and analyzed with the ImmunoSEQAnalyzer 3.0 software.

### Proteomic characterization by mass spectrometry

1×10^6^ Tregs were pelleted, snap-frozen and stored at −80°C before subjecting them to mass spectrometry-based proteome analysis as described previously^22^. Peak lists were searched and compared against the human Swiss-Prot database using PEAKS-studio-proteomics version 10.6 (Bioinformatics Solutions,). Peptide identification was performed using PEAKS-DB combined with PEAKS-de-novo sequencing. Label-free quantification with PEAKS-Q was used, with a false discovery rate set to 0.01.

### DigiWest

T-cell lysates were subject to SDS-PAGE. Eluted proteins were incubated with protein specific antibodies as described previously ^23^. Antibody specific signals were evaluated using DigiWest analysis tool and subjected to non-parametric statistical analysis and hierarchical clustering.

### Cellular indexing of transcriptomes and epitopes sequencing (CITE-seq)

Treg^WT^, FKBP12^KO^-Treg and Tcon were pooled post-labeling with TotalSeqC anti-human antibodies for surface characterization (Suppl. Table 3). Single cell suspensions were loaded onto Next-GEM-Chip-G. Tagged antibody, transcriptome and TCR libraries were prepared using the Chromium Single Cell 5′ Library & Gel Bead Kit as well as the Single Cell 5′ Feature Barcode Library Kit. Libraries for gene expression and CITE-seq were prepared using Single-Index-Kit-T-Set-A/Single-Index-Kit-N-Set A and quantified using Qubit-HS-DNA-assay kit. Further, fragment sizes were determined using 2100-Bioanalyzer with the High-Sensitivity-DNA-Kit. Sequencing was performed on a NextSeq500 device using High-Output-v2-Kits.

### Expansion of Treg products in the presence of IS

Following 21 days of expansion, the Tregs were cultured in 24-well plates for 3 or 10 days with or without tacrolimus (stock solution solved in ethanol 96%, reconstituted in PBS) and CsA (reconstituted in Ampuwa) at concentrations of 6ng/ml and 120 ng/ml, respectively.

### Off-target nomination (UNCOVERseq)

To nominate off-targets of the FKBP12 sgRNA, UNCOVERseq was performed as previously described^24^. Briefly, HEK293-Cas9 cells HEK293-Cas9 (ATCC) cells were cultured in Eagle’s Minimum Essential Medium (EMEM; ATCC) supplemented with 10% FBS at 37°C with 5% CO2. For each transfection, 8.0 x 10^5^ cells were washed with 1X phosphate-buffered saline, and resuspended in 20 µL of solution SF (Lonza). Following this, 5 µM gRNA and 0.5 µM dsODN were added to the SF solution. This mixture was transferred into 1 well of a 96-well Nucleocuvette plate (Lonza) and electroporated using program DS-150 with biological triplicate treatment/control samples. Following electroporation, cells were transferred to a 6-well plate preheated with EMEM and were incubated at 37°C with 5% CO2 for 72 hours. After incubation, genomic DNA (gDNA) was extracted using the MonarchTM Spin gDNA Extraction Kit (New England Biolabs) according to the manufacturer’s instructions, eluted in low-EDTA TE buffer (IDT, 11-05-01-05), and quantified using a NanoDrop 8000 UV-Vis Spectrophotometer (ND-8000-GL). 500 ng of purified gDNA was enzymatically fragmented and adapter-ligated using the xGen^TM^ DNA Library Prep EZ UNI kit along with the xGen Deceleration Module (IDT, xGen DNA Library Prep EZ UNI 96 rxn, 10009822; xGen Deceleration Module 96 rxn, 10009823) according to the manufacturer’s instructions and cleaned with AMPure XP beads (Beckman).

### Off-target nomination (CAST-seq)

CAST-Seq analyses were performed as previously described, with some adjustments to the workflow^25,26^. In brief, the average fragmentation size of the genomic DNA was aimed at a length of 500 bp, and the libraries were sequenced on a NovaSeq 6000 using 2×150 bp paired-end sequencing (GENEWIZ, Azenta Life Sciences). Sites under investigation were categorized as off-target mediated translocation (OMT) if the p-value met the cut-off of 0.005.

### Off-target Nomination (in silico)

To nominate sites *in silico,* CALITAS^27^ was used to nominate off-target sites both in the reference genome (GRCh38) and with consideration of known mutations from haplotype resolved variants from the Human Genome Diversity Project (HGDP) and the Genome Aggregation Database (gnomAD) occurring at >1% in any of the annotated super-populations. Super populations evaluated included African (AFR), Admixed American (AMR), East Asian (EAS), Non-Finnish Eurpoean (NFE), and South Asian (SAS) populations.

### Computational Analysis (UNCOVERseq)

Following NGS (2×150), analysis of UNCOVERseq data was performed as previously described^24^. Briefly, Illumina adapters and UMIs were identified and annotated using Picard MarkIlluminaAdapters. Tag sequences were identified and trimmed using Cutadapt v4.248. Sequencing reads were aligned to hg38 (GRCh38) reference genome using BWA mem v0.7.15 and UMI consensus reads were generated based on consensus from a single-strand (minimum UMI consensus size = 1) using fgbio v0.7.0 (https://github.com/fulcrumgenomics/fgbio). Nomination of candidate off-target sites began by using mapped UMI consensus reads to create a flanked search space (+/− 40 bp) to perform alignment between the guide and empirical target region using a glocal implementation of the Needleman-Wunsch alignment. After a candidate match to the gRNA spacer region was identified in the sequencing data, nominated off-target sites were identified using a hypergeometric test with multiple testing correction (Benjamini & Hochberg; FDR<0.05) by comparing individual treatment samples and pooled control samples for significant differences in representation between the two. We used the following criteria to nominate off-target sites from this analysis for verification: 1) at least one sample nominated a given site with NGS evidence on both sides of the cut site, 2) Levenshtein distance < 7 as determined post-alignment, and 3) significant adjusted p-value when comparing the frequency of the event to the pooled control(s). Nominated on/off-target sites had additional meta-data added based on alignment/genomic context and were placed into described Tiers based on this meta-data (Suppl. Tables 7 and 8).

### Off-target Prioritization

Candidate off-target sites nominated by different methods are associated with method-specific unique enrichment scores, supporting information, and annotations. To prioritize off-targets for confirmation a ruleset was used to determine the appropriate off-target sites for interrogation. Off-targets were prioritized if they met one or more of following criteria: 1) They were Tier 1 – Tier 3 UNCOVERseq nominations 2) they were derived Abnoba-seq 3) they were reproducibly derived from CAST-seq 4) they were an *in silico* derived site with a Levenshtein distance < 3 from the intended gRNA spacer or 5) they were an *in silico* derived site positioned within an annotated exonic region. All sites were designed for multiplexed amplicon sequencing using the IDT rhAmpSeq Design Tool (https://www.idtdna.com/pages/tools/rhampseq-design-tool).

### Statistics

All data points represent biological replicates. Analysis was performed using GraphPad PRISM and R. Data are shown as mean±SEM unless otherwise stated.

## Results

### FKBP12^KO^ is feasible in Treg products

To avoid gene-editing of contaminating, potentially allo-reactive Teff, Treg isolation was optimized for increased purity prior to genetic modification by a combination of CD4 enrichment during density gradient centrifugation succeeded by flow cytometry sorting (Fig. 1A/B, suppl. Fig. 1A)^14^. Research grade Treg enrichment yielded approximately 3.75 - 7.5 x 10^7^ CD4^+^ T-cells per 50 ml blood (Fig. 1H, suppl. Fig 1C). Sorted Treg were subsequently stained for FOXP3, the master transcription factor of Treg (Fig. 1B). Sorting yielded cell fractions with a minimized contamination of <2% of CD3^-^ (Fig. 1C) or <0.05% CD3^+^CD8^+^ cells (Fig. 1D) and achieved >90% purity of the CD25^+^FOXP3^+^/CD127^-^CD25^+^ Tregs within CD4^+^ cells (Fig. 1E/Suppl. Fig. 1B). High-fidelity Cas9 editing achieved approximately 60% and standard Cas9 enzyme 80% efficiency in the final FKBP12^KO^-Treg product expanded for 21 days (Fig. 1F, suppl. Fig. 1D)^46^. Gene editing enhanced cell proliferation rate in the presence of IL-2, anti-CD3/CD28 microbeads and rapamycin (Fig. 1G), yielding a greater number of FKBP12^KO^-Tregs (Fig. 1H, suppl. Fig. 1C) >400×10^8^ cells, sufficient for clinical application if applying 3×10^6^ cells/kg bodyweight and including 1×10^8^ cells required for release testing^13,14^ (Suppl. Fig. 1C).

**Figure 1:**
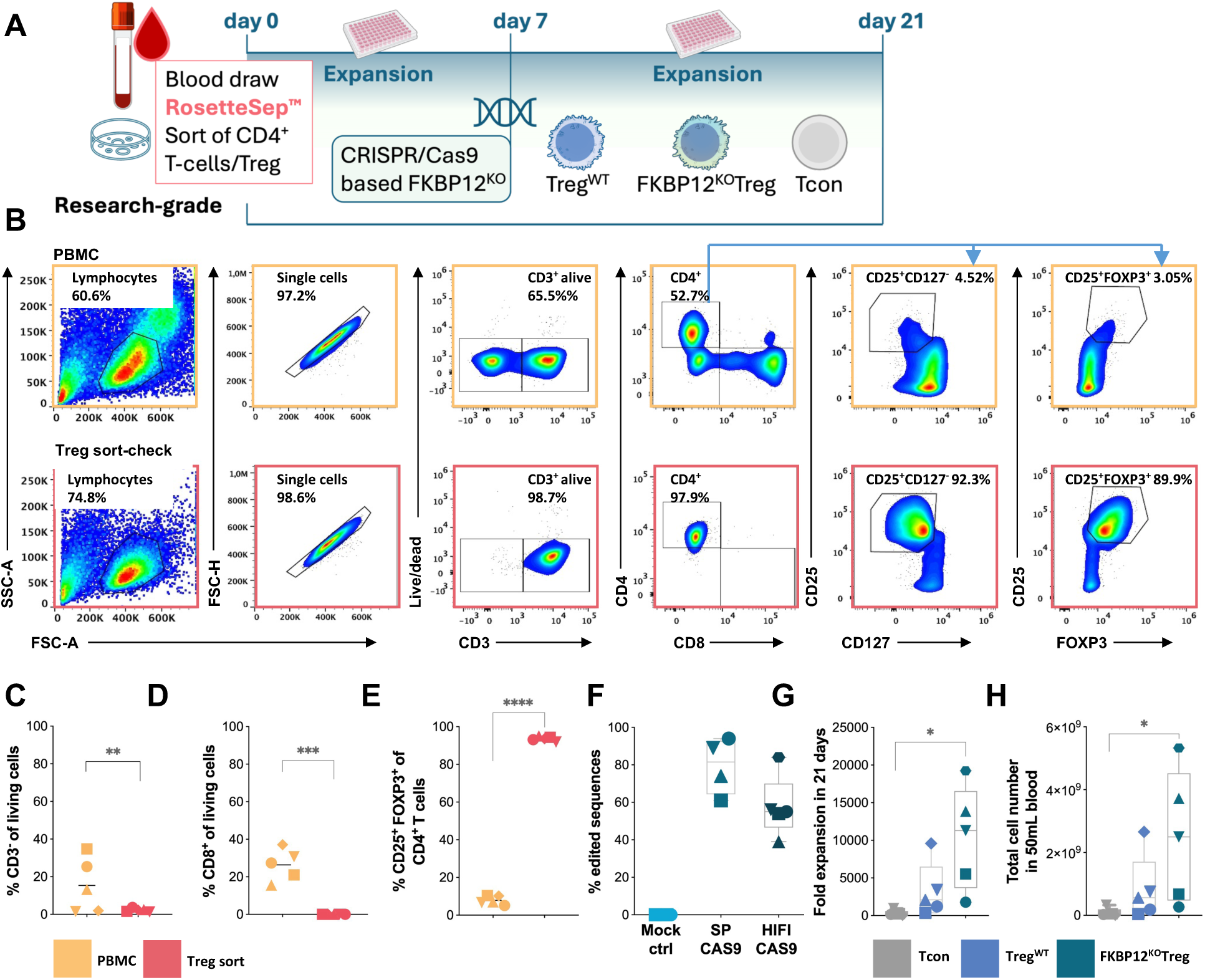
FKBP12KO-Treg are pure and show improved expansion. (A) A description of the Treg product production process. NIH Bioart source was used. (B) Representative dot plots show the gating strategy and phenotype of PBMCs before and after Treg sorting. For the following subfigures, each symbol represents data from an individual donor. Pre-sort (yellow) post-sort (red). Conventional T-cells (Tcon) (grey), Treg^WT^ (blue), FKBP12^KO^-Treg (teal) unless stated otherwise. ANOVA, post-test t-test, p<0.05(*), p<0.01(**), p<0.001(***), p<0.0001(****) n=5. (C) Percentage of CD3^-^ T-cell subset before and after sorting, p<0.01(**). (D) Percentage of CD8^+^ T-cells before and after sorting, p<0.001(***). (E) Percentage of CD4^+^CD25^+^FOXP3^+^ T-cells before and after sorting, p<0.0001(****). (F) Analysis of the KO efficiency of Treg after a 3-week expansion period, performed using Sanger sequencing of FKBP12 PCR products. Sequencing data was analysed using the Inference of CRISPR edits (ICE) tool (Synthego). Electroporation with Cas9 without sgRNA/mock electroporated (turquoise), with classical Staphylococcus pyogenes Cas9 (teal) and with a high-fidelity Staphylococcus pyogenes Cas9 (dark teal). (G) Fold expansion from day 0-21. Boxplots with whiskers indicate extreme values, n=5, p<0.05(*), p<0.01(**). (H) The total number of cells obtained from 50ml of blood is shown after 21 days of expansion, p<0.05(*).

### FKBP12^KO^-Treg quality is comparable with Treg^WT^

The FKBP12^KO^-Treg, Treg^WT^ and non-Treg CD4^+^ T-cells (negative sort fraction), were subjected to detailed analysis following expansion in identical conditions. The Treg markers; FOXP3 and CD25 were highly expressed by both Treg^WT^ and FKBP12^KO^-Treg, but also Tcon (Fig. 2A/B). However, the mean fluorescence intensity (MFI) of FOXP3 was brighter in Treg *vs*. Tcon and comparable between FKBP12^KO^-Treg and Treg^WT^ (Fig. 2C). Interestingly, CD25 MFI was lower among the FKBP12^KO^-Treg compared to Treg^WT^/Tcon (Fig. 2D). After PMA/Ionomycin stimulation, the effector cytokines IL-2 (Fig. 2E/F) and tumor necrosis factor (TNFα) (Fig. 2G) were produced at very low levels, medians of 1% and 2%, respectively, among Treg^WT^ and FKBP12^KO^-Treg compared with 15% and 40% among the parallelly expanded CD4^+^ Tcon. The median frequencies of producers of IFN-γ were determined to be 10% for Tcon; while 3% vs. 2% for FKBP12^KO^-Treg and Treg^WT^, respectively (Fig. 2E/H). Clonality determined by TCRβ sequencing was found to be similar for FKBP12^KO^-Treg and Treg^WT^ (Fig. 2I). While frequencies of CD137 expressing cells were similar among FKBP12^KO^-Treg and Treg^WT^, they were less frequent among Tcon (Suppl. Fig. 2A/B). Frequencies of cells expressing CD40L (CD154) were higher among Tcon than both Treg populations (Suppl. Fig. 2A/C). The TSDR methylation of the Tcon was found to be 85% and <20% among the FKBP12^KO^-Treg and Treg^WT^ (Fig. 2J). Treg suppressed effector T-cells (Teff) in a dose dependent manner (Fig. 2K, L).

**Figure 2:**
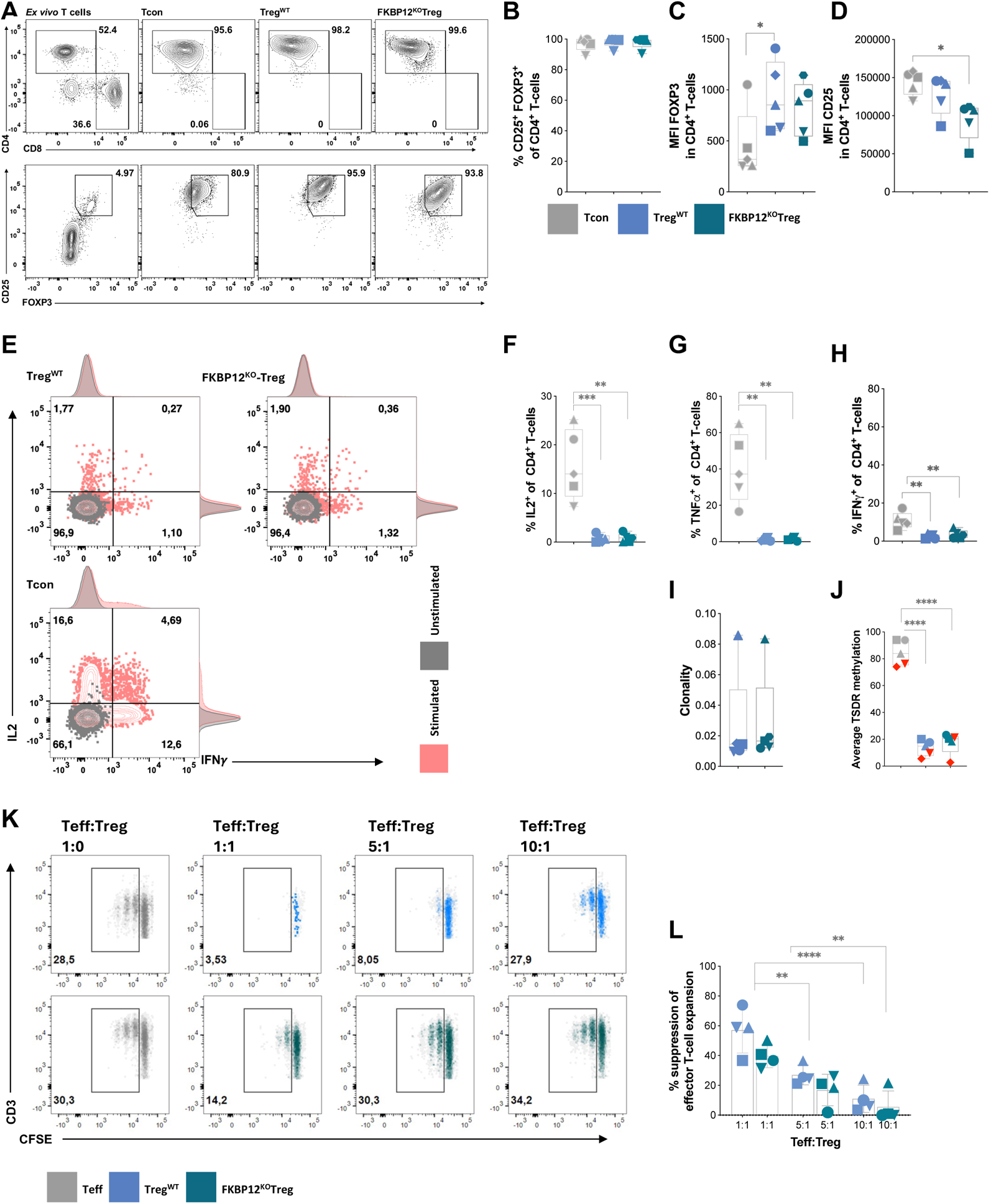
FKBP12KO-Treg phenotype and function is comparable to Treg^WT^. Each symbol represents data from an individual donor. Conventional T-cells (Tcon) (grey), Treg^WT^ (blue), FKBP12^KO^-Treg (teal). One-way ANOVA (and non-parametric), p<0.05(*), p<0.01(**), p<0.001(***), p<0.0001(****) mean +/− SD n=5. (A) Representative dot plot showing the gating strategy for living T-cells, CD4^+^, CD8^+^, and CD25^+^FOXP3. (B) The percentage of CD4^+^CD25^+^FOXP3^+^ T-cells on day 21. (C) MFI of FOXP3 at d21. (D) MFI of CD25 at d21. (E) Schematic outline of the experimental setup following 6h stimulation by PMA/Ionomycin and intracellular capture of cytokines by addition of brefeldin A after 45 min. NIH Bioart source was used. (F) Representative dot plots for the producers of IL-2 and IFNLJ, unstimulated (grey) and stimulated (pink). (G) Summary of the percentage of IL-2 producers (H) TNFα producers (I) IFNLJ producers after stimulation. (J) Comparison of FKBP12^KO^-Treg clonality with Treg^WT^ using TCRb sequencing. (K) TSDR demethylation at the promoter of FOXP3 of DNA collected on day 21: Female donors highlighted in red. (L) Representative dot plots and gating of proliferated responder T-cells (Teff) co-cultured with Treg at different Teff:Treg ratios and stimulated 1:1 with anti-CD3/CD28-coated microbeads. (N) The percentage of Treg suppressive capacity on CFSE labelled autologous fresh CD3^+^ T-cells stimulated with CD3/CD28 beads, p<0.01(**).

### FKBP12^KO^-Treg exhibit enhanced characteristics when exposed to tacrolimus

FKBP12^KO^ conferred proliferative advantage over Treg^WT^ when exposed to tacrolimus (Fig. 3B). Furthermore, the expression of CTLA-4 was increased in FKBP12^KO^-Treg *vs.* Treg^WT^ (Fig. 3C/E). Tacrolimus-treated FKBP12^KO^-Treg exhibited significantly brighter PD1 MFI in comparison to Treg^WT^ (Fig. 3C/D). Additionally, Treg^WT^’s CTLA-4 MFI was diminished relative to medium controls in presence of both CNIs, but not in FKBP12^KO^-Treg treated with tacrolimus (Fig. 3E). Interestingly, knocking out FKBP12 resulted in downregulation of the FKBP1A gene, underlining knock-out efficiency and further affected regulation of a number of other proteins (Fig. 3G-M); (i) increased metabolic arginase-2 (FOXP3 stability) ii) decreased NIT2 levels (histidine metabolism) in comparison to Treg^WT^, (iii) significantly, upregulated Guanylate binding protein-2 (Treg suppressive capacity) and (iiii) downregulated zyxin (stress fiber protein) when exposed to tacrolimus ^49–52,28^. Pathway analysis identified differentially regulated proteins in FKBP12^KO-^Treg and Treg^WT^ highlighting increased IL-2 and IL-10 signaling as well as regulation of TNFR signals. Furthermore, IFNγ signaling was decreased in FKBP12^KO^-Treg compared to Treg^WT^ (Fig. 3N).

**Figure 3:**
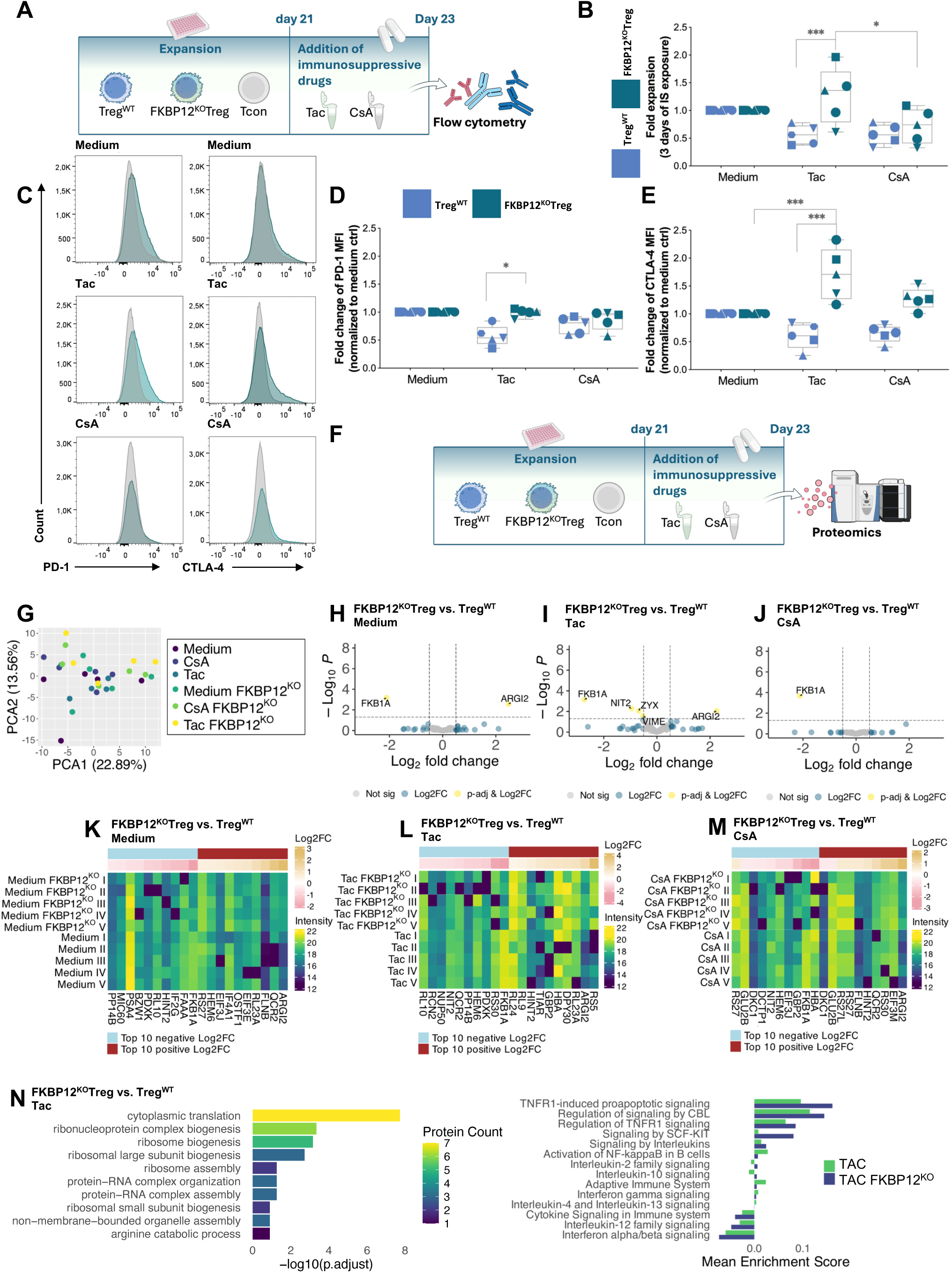
FKBP12KO-Treg show preferable characteristics under tacrolimus. Each symbol represents data from an individual donor. Tacrolimus was used at 6 ng/ml and cyclosporine (CsA) at 120ng/ml with Treg^WT^ (blue), FKBP12KOTreg (teal). One-way ANOVA (and non-parametric), p<0.05(*), p<0.01(**), p<0.001(***), p<0.0001(****), mean +/− SD n=5. (A) Schematic overview of Treg expansion in the presence of indicated immunosuppression. NIH Bioart source was used. (B) Treg products were exposed to the indicated immunosuppression for a total of 3 days post 3-weeks of manufacturing. Data normalization using Remove baseline and column math’ function of GraphPad Prism, p<0.05(*), p<0.001(***), p<0.0001(****). (C) Histograms of CTLA-4 and PD-1 staining of Treg^WT^ and FKBP12^KO^-Treg products exposed to indicated immunosuppression. (D) Flow cytometry analysis showing the fold change of MFI of PD-1 in Treg^WT^ and FKBP12^KO^-Treg and (E) CTLA-4 after exposure to the indicated immunosuppression, data normalization using Remove baseline and column math’ function of GraphPad Prism, p<0.05(*), p<0.001(***). (F) Schematic overview of proteomics approach. NIH Bioart source was used. (G) Principal component analysis (PCA) of Treg^WT^ and FKBP12^KO^ samples under medium control conditions or following treatment with Cyclosporin A (CsA) or Tacrolimus (Tac). (H-J) Differential expression analysis comparing FKBP12^KO^-Treg versus Treg^WT^ under medium control conditions and following Tac or CsA treatment. Volcano plots display −log10 adjusted p-values (y-axis) versus log2 fold change (Log2FC, x-axis). Proteins with Log2FC > 0.5 are highlighted in blue; proteins with Log2FC > 0.5 and adjusted p-value < 0.05 are highlighted in yellow. (K-M) Heatmaps displaying intensity levels of the top 10 proteins with highest positive Log2FC and top 10 proteins with highest negative Log2FC comparing FKBP12^KO^-Treg versus Treg^WT^ under medium control conditions and following Tac or CsA treatment. (N) Left: Gene Ontology (GO) pathway enrichment analysis showing the top 10 enriched pathways comparing FKBP12^KO^-Treg versus Treg^WT^ following Tacrolimus treatment. The −log10 adjusted p-value is displayed, with the number of enriched proteins in each pathway indicated by color coding. Right: Single-sample Gene Set Enrichment Analysis (ssGSEA) of FKBP12^KO^-Treg and Treg^WT^ following Tac treatment using the Reactome database. Mean enrichment score of selected pathways is shown.

### FKBP12^KO^Treg can be produced under pre-GMP conditions

We adapted the production process for FKBP12^KO^-Treg to pre-GMP conditions using CliniMACS Plus to isolate CD4^+^ cells from whole blood (Fig. 4A), which resulted in a product with < 0.002% CD8^+^ cell contamination Fig. 4B/C). Afterwards, it was possible to isolate CD25^+^FOXP3^+^ cells with > 85% purity among CD4^+^ cells (Fig. 4D/E). Following expansion, pre-GMP FKBP12^KO^-Treg and Treg^WT^ had comparable FOXP3 MFI, but both significantly lower CD25 MFI as Tcon (Fig. 4F/G) with negligible changes in effector cytokine expression between FKBP12^KO^-Treg and Treg^WT^ (Fig. 4H-K). TSDR methylation levels were lower in both Treg products compared to Tcon (Fig. 4L).

**Figure 4:**
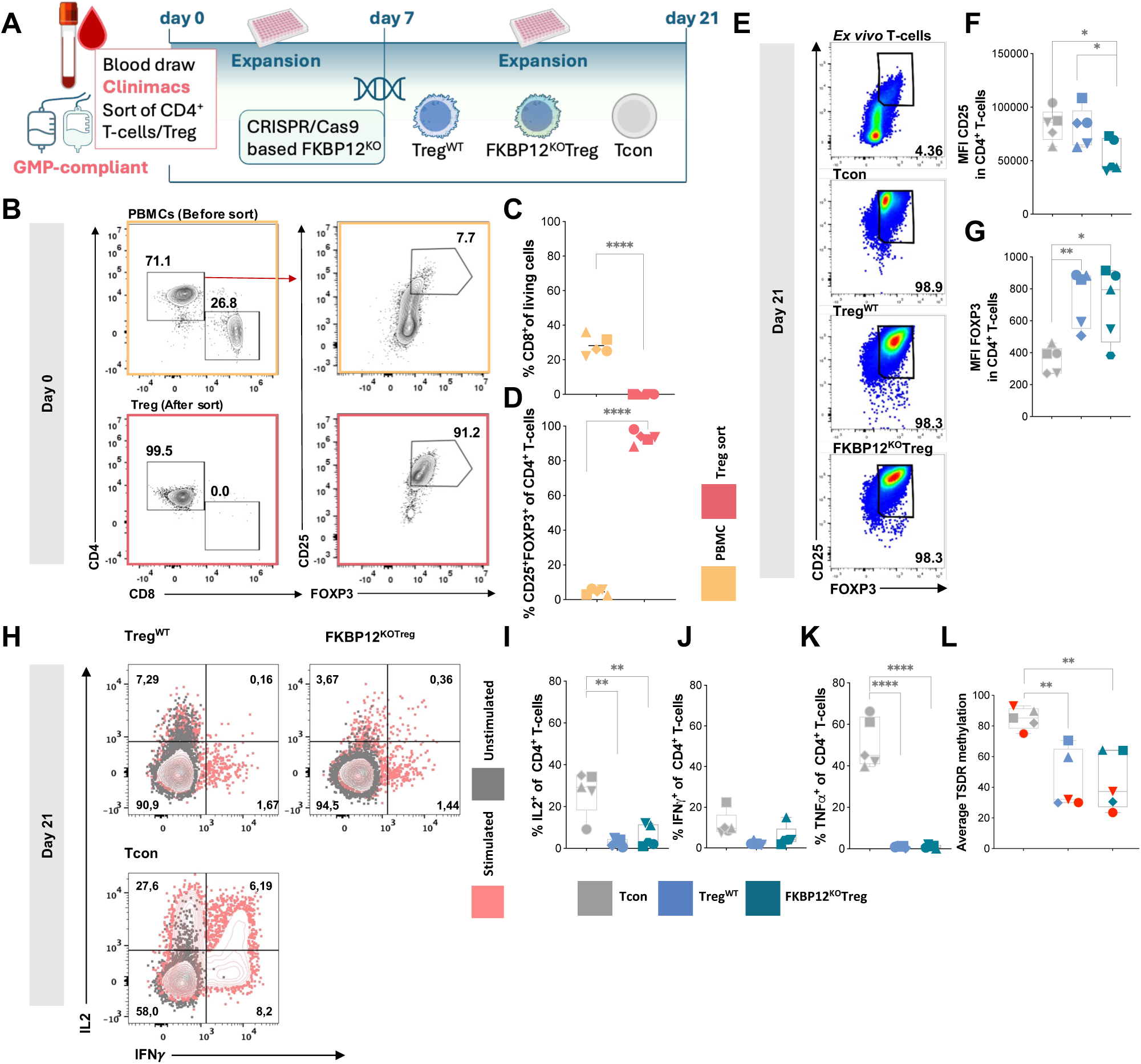
FKBP12KO-Treg can be manufactured in a GMP compatible manner (“pre-GMP”) Each symbol represents data from an individual donor. Pre-sort (yellow) post-sort (red). ANOVA, post-test t-test, p<0.05(*), p<0.01(**), p<0.001(***), p<0.0001(****). Conventional T-cells Tcon) (grey), Treg^WT^ (blue), FKBP12^KO^-Treg (teal). One-way ANOVA (and non-parametric), mean +/− SD n=5. (A) A description of the production process. NIH Bioart source was used. (B) Percentage of CD8^+^ of living T-cells before and after sorting. (C) Percentage of CD25^+^FOXP3^+^ CD4^+^ T-cells before and after sorting. (D) Representative dot plots and gating for CD4^+^, CD8^+^, and CD25^+^FOXP3^+^ of living T-cells before and after sorting. (E) MFI of FOXP3 on day 21. (F) MFI of CD25 on day 21, p<0.05(*). (G) A representative dot plot showing percentages of CD25^+^FOXP3^+^ T-cells in *ex-vivo* T-cells, Tcon, Treg^WT^ and FKBP12^KO^-Treg on day 21. (H) Representative dot plots of producers of IL-2 and IFNLJ following 6 h stimulation by PMA/Ionomycin and intracellular capture of cytokines by addition of brefeldin A after 45 min. Unstimulated (orange), stimulated (black) (I)(J)(K) Summary of the percentage of IL-2, TNFα and IFNLJ producers. (L) Methylation of the TSDR at the promoter of FOXP3 of DNA collected on day 21, p<0.01. Female donors are highlighted in red.

### Pre-GMP FKBP12^KO^-Treg mirror enhanced characteristics of research grade FKBP12^KO^-Treg

To test the FKBP12^KO^-Treg expansion in comparison to Treg^WT^, we expanded both in culture media containing CNIs+IL2 for 10 days. FKBP12^KO^-Treg proliferation was superior in comparison to Treg^WT^ in the presence of tacrolimus, but both showed sensitivity to CsA (Fig. 5A). Both products maintained FOXP3 expression in contrast to Tcon (Fig. 5B). Significantly, the CD25 MFI variance between FKBP12^KO^-Treg and Treg^WT^ was not observed after 10-days exposure to tacrolimus (Fig. 5C). DigiWest based proteome analysis confirmed upregulation of already described proteins in FKBP12^KO^-Treg *vs*. Treg^WT^ (Fig. 5D, Fig. 3 I/K/M)^23^. Reactome pathway analysis showed enrichment of ribosomal processes correlating with single upregulated proteins in FKBP12^KO^-Treg vs. Treg^WT^ (Fig. 3 N). Furthermore, FKBP12^KO^-Treg show upregulation of IL-2 and IL-10 family signaling as well as increased downregulation of cytokine, especially IFNα/β signaling under tacrolimus treatment (Fig. 3O). Interestingly, CD4 was upregulated and CD8a was downregulated in tacrolimus-treated FKBP12^KO^-Treg *vs.* Treg^WT^. CTLA-4 gene expression in FKBP12^KO^-Treg increased under tacrolimus (Suppl. Fig. 4E). Moreover, we observed increased CD25 (IL2Ra) and phospho-ERK (Erk1/2/Thr202/Tyr204) expression in tacrolimus-treated FKBP12^KO^-Treg *vs.* Treg^WT^ (Fig. 5D) ^29^. These findings align with increased levels of activation markers e.g. GITR and LAG3^30,31^. STAT5 and pSTAT5 protein was shown to be upregulated in FKBP12^KO^-Treg *vs*. Treg^WT^ upon exposure to tacrolimus (Fig. 5D, suppl. Fig. 4B).

**Figure 5:**
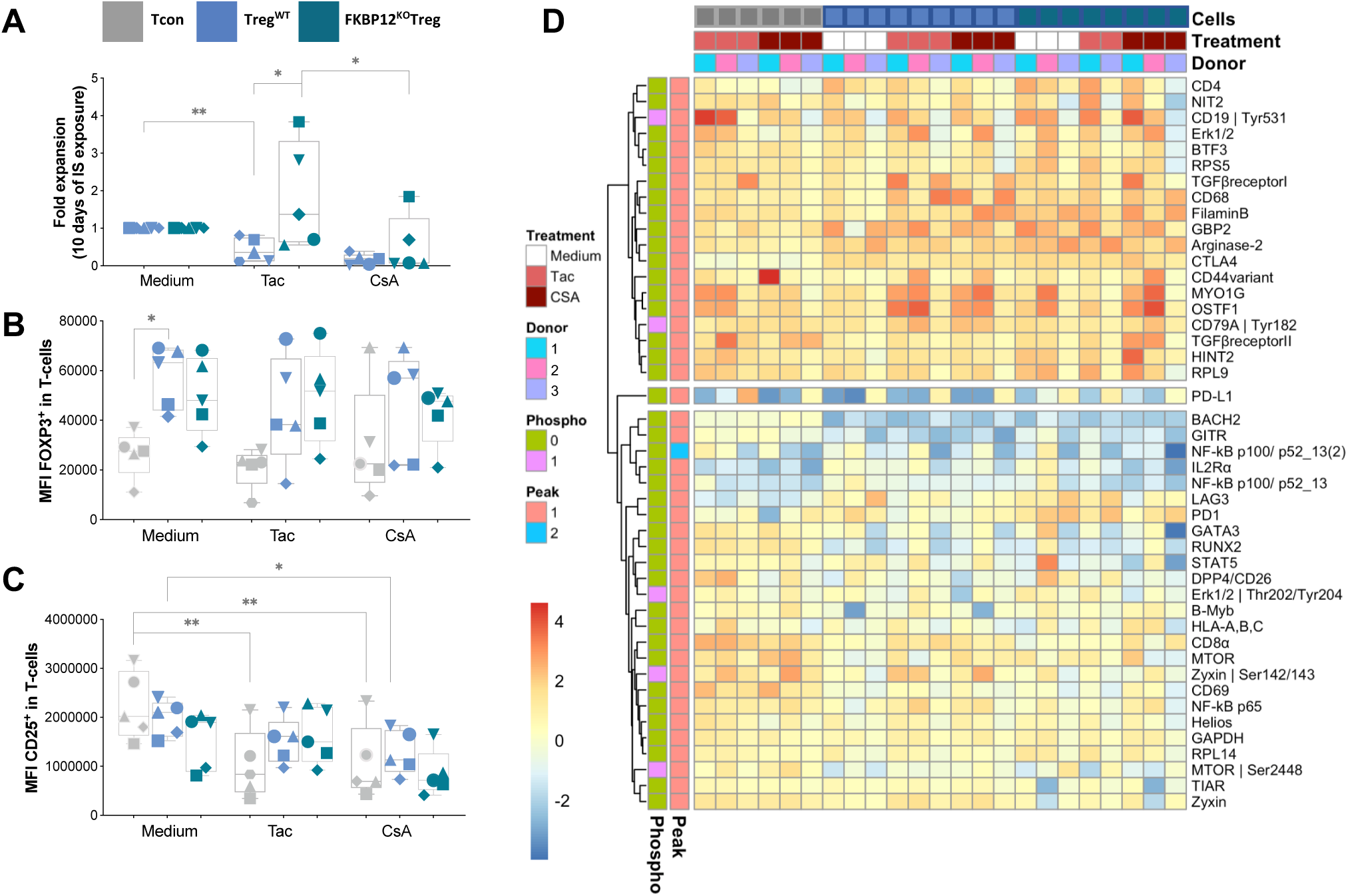
Characterization of Treg Response to Immunosuppression: Impact of Pre-GMP FKBP12^KO^-Treg. Each symbol represents data from an individual donor. Treg products were exposed to 6 ng/ml tacrolimus or 120 ng/ml cyclosporine (CsA) for 10 days, immunosuppressants were added and re-stimulation was performed every 2nd day. Conventional T-cells (Tcon, grey), Treg^WT^ (blue), FKBP12^KO^-Treg (teal). One-way ANOVA (and non-parametric), p<0.05(*), p<0.01(**), p<0.001(***), mean +/− SD n=5. (A) Expansion of Treg products in presence of immunosuppressants, data normalization was performed using remove baseline and column math’ function of GraphPad Prism p<0.05(*), p<0.01(**), p<0.001(***). (B) MFI of FOXP3 of living T-cells and (C) MFI of CD25^+^ is shown for all conditions. (D) The heat map depicts the differentially expressed proteins of expanded Tcon, Treg^WT^, and FKBP12^KO^-Treg exposed to tacrolimus or CsA medium control and measured by DigiWest technology.

### Cellular Indexing of Transcriptomes and Epitope sequencing (CITE-seq) confirms pre-GMP FKBP12^KO^-Treg’s functionality

Using single-cell CITE-seq data, 14 cell clusters were annotated, which were shared between for FKBP12^KO^ and Treg^WT^, but distinct from the Tcon following (re)stimulation (Fig. 6A-C, Suppl. Fig. 3). The cluster composition of both FKBP12^KO^-Treg and Treg^WT^ changed in the presence of CNIs mainly regarding cluster 11 and 14 (Suppl. Fig. 3). However, FKBP12^KO^-Treg were not affected by the CNI tacrolimus (Fig. 6C). We compared FKBP12^KO^ and Treg^WT^ with respect to expression of key genes (Fig. 6D). Compared with Treg^WT^, FKBP12^KO^ showed an increase in T helper 2 (TH2) signature e.g. IL-5 upregulation in control medium and tacrolimus conditions, while under CsA treatment TH2 signature was reduced (Fig. 6D, E). KEGG and GOTERM highlighted TH2 signaling, regulation of IL-10 and B-cell activation pathways (p>0.005, Suppl. Table 4). Furthermore, LAG3, STAT5, IL-10 and TNFR gene and protein expression levels were upregulated in FKBP12^KO^*-*Treg vs. Treg^WT^ especially when exposed to tacrolimus (Fig. 5D, Fig. 6E/F). Tacrolimus treated FKBP12^KO^*-*Treg showed higher PD-1 levels and increased TIGIT levels compared to Treg^WT^ under tacrolimus (Fig. 6D, Suppl. Fig. 4A). Additionally, CTLA-4 and CD73 gene and surface protein expression levels were upregulated and preserved under tacrolimus (Suppl. Fig. 4C/D/E).

**Figure 6:**
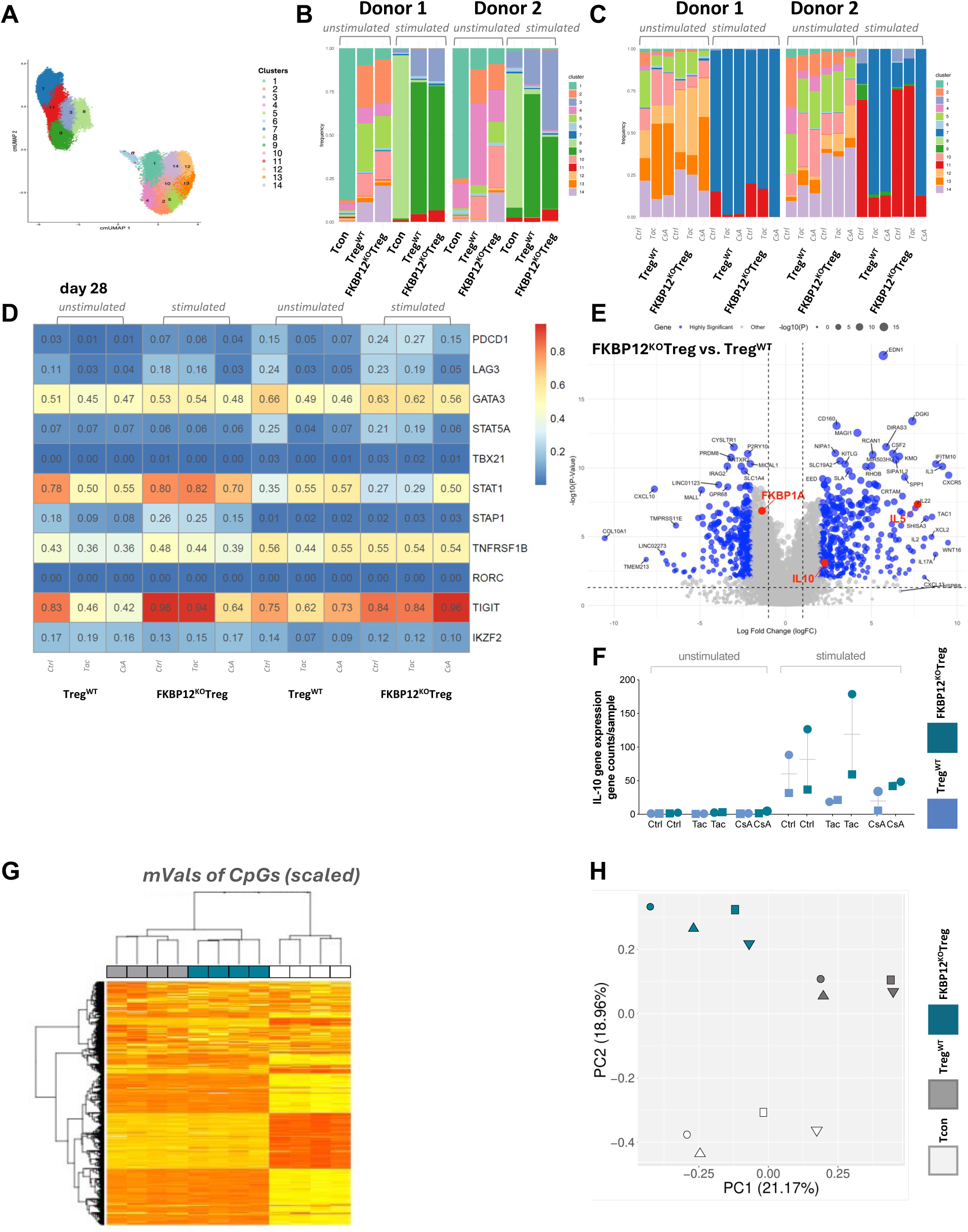
Tacrolimus resistance and CsA safety switch are visualized by CITEseq of FKBP12^KO^-Treg and methylation profile of FKBP12^KO^-Treg proves identity. Where indicated, Treg products were exposed to 6 ng/ml tacrolimus or 120ng/ml cyclosporine A (CsA). CITEseq was performed at day 21 and day 28, the latter after a 7-day exposure to tacrolimus or cyclosporine A (CsA) or medium as a control from n=2 donors. (A) UMAP clustering of all analysed cells showing 14 different clusters. The assigned clusters are marked based on the colour code indicated. (B) Cluster distribution is shown for an unstimulated state and after 6h stimulation with PMA/Ionomycin on day 21. (C) Cluster distribution is shown after exposure to the indicated immunosuppressant for 7 additional day. (D) A comparison of gene expression data between FKBP12^KO^-Tregs to Treg^WT^ in unstimulated and stimulated conditions. Log2 fold change (Log2FC) is displayed after 28 days of exposure to ctrl medium, Tac and CsA. (E) Volcano plot displaying differential gene expression within FKBP12^KO^-Treg and Treg^WT^ on day 21 (F) Differential gene expression of IL-10 is plotted for unstimulated and stimulated FKBP12^KO^-treg in comparison to Treg^WT^ and under supplementation of ctrl medium, Tac and CsA for 7 additional days. (G) PCA analysis of data determined in EPIC arrays showing clustering for Tcon (white), Treg^WT^ (grey) and FKBP12^KO^-Treg (teal).

### Pre-GMP FKBP12^KO^-Treg and Treg^WT^ have different epigenetic profiles

FKBP12^KO^-Treg, Treg^WT^ and Tcon were subjected to an EPIC array and revealed distinct Treg and Tcon clusters in PC2. Furthermore, PC1 separated FKBP12^KO^-Treg and Treg^WT^ irrespective of donor origin (Fig. 6G/H).

### Off-target nomination

We performed CAST-Seq, in silico population based nominations and UNCOVERseq to nominate potential off-targets hit by the sgRNA (Suppl. Table 5). Nomination of off-targets using UNCOVERseq in a promiscuous nomination system (HEK293-Cas9) discovered 258 off-targets significantly enriched (p_adj_ < 0.05) in comparison to unedited samples. Of these off-targets, 182 met Tier1 to Tier 3 criteria for prioritization, and 133 of these were reproduced between 2 or more biological replicates (Suppl. Table 5). Reproducibly nominated off-targets occurred in the following genomic regions: 9.7% exonic (13 / 133), 45.8% intronic (61 / 133) and the remaining sites in intergenic or atypical genomic context (Suppl. Table 5). CAST-seq identified 10 off-target sites reproducibly, and *in silico* nomination identified an additional 103 unique sites. This led to a total of 255 putative off-target sites being prioritized for off-target confirmation (Suppl. Table 5). Confirmation of prioritized off-targets nominated is currently ongoing.

### GMP process validation

Based on the reseach grade and pre-GMP data, we decided to develop GMP compatible manufacturing using semi-closed bioreactors for expansion and the newly established isolation and gene-editing processes described above. Indeed, we succeeded in producing stable “TregTacRes” products with >94% viable cells, being >95% CD4^+^CD25^+^FoxP3^+^, with contaminations of less than 0.3% CD8+ T cells, <6% IL-2 and IFNγ producers, including >79% of FKBP12^KO^-Treg (Table 1). Validation of KO efficacy in FKBP12^KO^-Treg was implemented using a ddPCR as qualified method for GMP manufacturing showing validated protocol results in comparison to non-validated results and Sanger sequencing (Fig. 7AB, Table 1, Suppl. Table 6). Based on these and other dara, e.g. sterility testing according to the European Pharmacopoeia, the local authority granted us manufacturing authorization for the “TregTacRes” product on May 16^th^, 2024.

**Figure 7:**
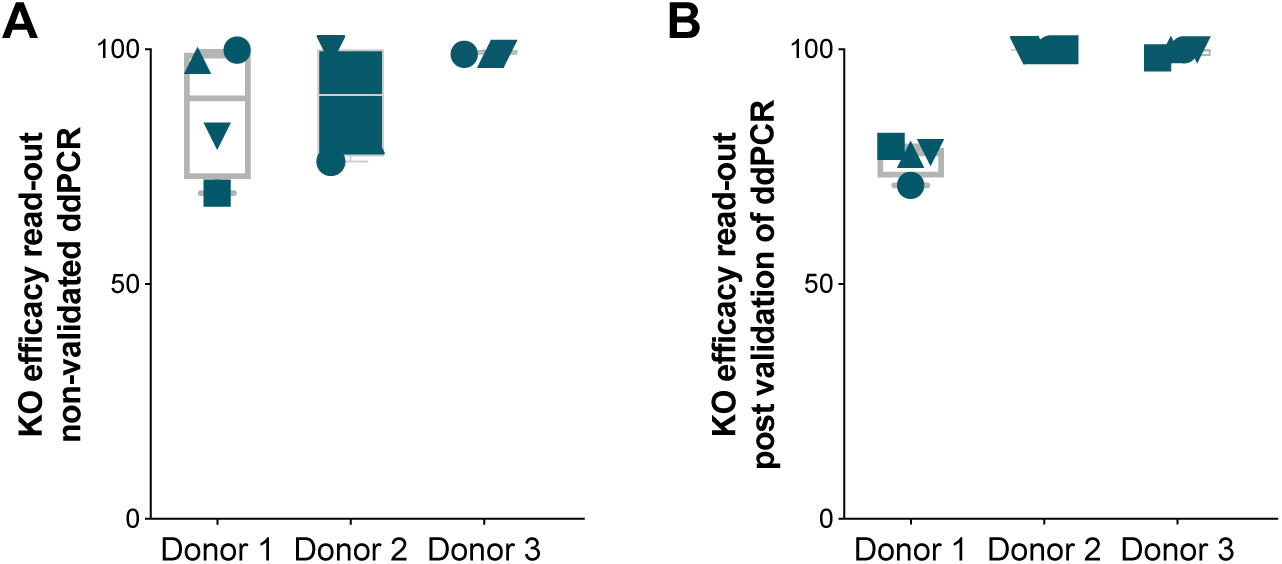
Validation results of KO-efficacy in FKBP12^KO^-Treg. KO-efficacy results are shown in (A) for non-validated method and in (B) for the validated approach. Three different donors were analysed for their FKBP12 KO efficacy while using 4 independent replicate samples each.

**Table 1:**
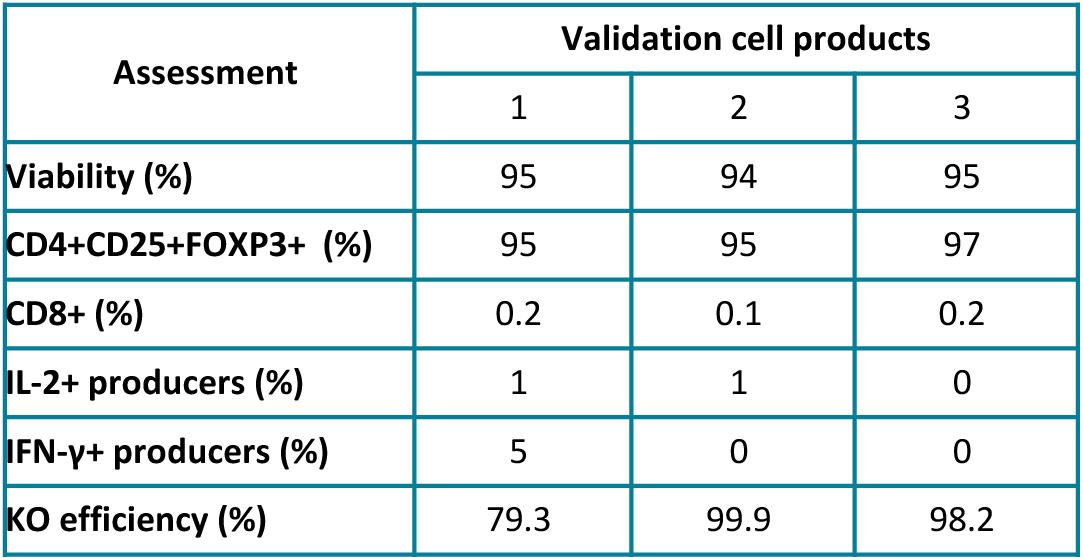
Assessment and validation of KO-efficacy in FKBP12^KO^-Treg. Data from GMP compliant validation runs is shown containing parameters such as % of viability, Treg, CD8 contamination, IL2/IFNγ/TNFα producers and final KO efficacy.

## Discussion

Drugs like tacrolimus revolutionized transplantation of organs between individuals regulating immune responses artificially^32^. In contrast, Treg are “living drugs”, that enable endogenous, regenerative immunosuppression without toxicities^33^. We envision to overcome tacrolimus adversely affecting adoptively transferred Treg by a next-generation FKBP12^KO^-Treg product resistant to tacrolimus. In our hands, research grade, pre-GMP and GMP manufacturing of Treg demonstrated FKBP12^KO^-Treg to be safe and efficacious in presence of tacrolimus. Analysis of potential off-target editing is ongoing as a pre-requisite to enter clinical phases. We suggest FKBP12^KO^-Tregs will significantly enhance *in-vivo* functionality, efficacy and longevity in patients prescribed with tacrolimus.

### Manufacturing and performance

We were able to generate a pure FKBP12^KO^-Treg product from 50 ml of whole blood with sufficient cell numbers for clinical trials applying e.g. 3×10^6^ Treg/kg^14,34,35^. FKBP12^KO^-Treg exhibited resistance to tacrolimus, but maintained sensitivity to CsA, thus providing a fail-safe clinical management^36,37^. FKBP12 is known to bind rapamycin which impedes T cell proliferation^38^. Consistent with this, we observed FKBP12^KO^ conferred proliferative advantage under IL-2 and rapamycin, when compared with Treg^WT^, probably due to dependence of rapamycin-mediated anti-proliferative effects on FKBP12 ^39,40,41^. Excluding any potential gene-edited Tcon impurities^42–44^, we observed reduced CD8a protein levels in FKBP12^KO^-Treg vs. Treg^WT^. Consistent with published data on general FOXP3 upregulation in Treg as well as activated CD4^+^ T-cells, we show that FOXP3 was upregulated in activated Tcon, however the MFI was diminished in comparison to Treg ^45–47^. The TSDR methylation levels were similar in Treg^WT^ and FKBP12^KO^-Treg while Tcon showed increased methylation, thus confirming Treg identity and stability according to published scientific guidelines ^48,49^. Tacrolimus-treated FKBP12^KO^-Treg and untreated Treg^WT^ cluster distributions indentified in CITEseq experiments were similar, confirming FKBP12^KO^ did not alter the Treg product composition ^18^. Furthermore, common phenotypic markers were similarly expressed by FKBP12^KO^-Treg and Treg^WT^. Tacrolimus is known to stabilize CD25 and induce loss of FOXP3 upon adoptive Treg transfer *in-vivo*^50^. Our *in-vitro* data shows stable CD25 protein expression in presence of tacrolimus in both FKBP12^KO^-Treg and Treg^WT^, as well as stable FOXP3 expression and demethylation. Despite the reduced CD25 expression levels in conditions without tacrolimus treatment, the FKBP12^KO^-Treg showed proliferative advantage and phosphor-STAT5 levels, a signaling protein phosphorylated during IL-2 signal transduction, were comparable to Treg^WT^. However, untreated FKBP12^KO^-Treg showed decreased suppressive capacity in the classical suppression assay ^51^, which may be due to diminished competition for IL-2 conferred by CD25 ^52,53^. Unfortunately, suppression assays in presence of tacrolimus were not feasible as the tacrolimus impact on Tcon proliferation was dominant. Of note, Treg suppress by a myriad of additional mechanisms ^53^, but there is currently no standardized assay to evaluate all modes of action. We show increased IL-10 mRNA levels and enhanced IL-10 signaling as well as increased CD73 mRNA in FKBP12^KO^-Tregs implying greater suppressive capacity. Furthermore, IL-2 protein signaling was increased in FKBP12^KO^-Treg in part compensating for CD25 reduction in presence of tacrolimus. Hereby highlighting alternative signaling pathways such as heterodimeric CD25-chemokine receptor pathway^54^.

Furthermore, FKBP12^KO^-Treg show favorable CTLA-4 expression upon exposure to tacrolimus ^55–58,59^ and upregulation of genes associated with Treg function, stability and immune suppression. We saw scarce levels of IFNγ^+^ FKBP12^KO^-Treg and Treg^WT^, which were not associated with TH1-Treg phenotype shown by the low expression Tbet. GATA3 expression pointing towards a TH2-Treg phenotype will be further investigated.

We validated production of the “TregTacRes” in our GMP unit and were granted manufacturing authorization.

### Safety

To circumvent safety issues of viral vectors, we elected vector-free gene-editing using CRIPSR-Cas9 RNPs and a high-fidelity Cas9 enzyme to prevent off-target editing for pre-GMP and GMP runs. We nominated putative off-targets using different in silico and cell-based prediction approached and validation of these off-targets via amplicon sequencing (rhAMPseq) is ongoing ^60,61^. TCRβ sequencing revealed no skewing of FKBP12^KO^-Treg *vs*. Treg^WT^ TCR repertoires implying transformative events to be unlikely. We are now preparing to test co-administration of FKBP12^KO^-Treg and tacrolimus within a clinical trial in solid organ transplant recipients.

## Authorship

Conceptualization, L.A., M.S.H, P.R. and H.D.V.; Methodology: F.I., A.R., D.K., J.P.B.,, T.C. and D.L.W., single cell sequencing: M.F.M., proteomics: O.K., epic arrays: K.O., DigiWest: M.T., S.F.; Software and formal analysis, single cell/proteomics and epic array: S.S., A.G., P.D., F.H., O.K., K.O., L.E.; Investigations: G.Z., D.J.W., M.St., H.H., I.S., S.D., Jo.K., L.P., S.P., R.F., L.A., D.L.W., S.M., M.S., A.J., R.T., M.G., G.L.K., G.R., J.K., O.M.C., M.M.K., P.D., M.F.M.; Data Curation, S.S., A.G., P.D., F.H.; Writing, L.A., G.Z., L.M.B., Ja.K.; Review: all co-authors; Visualization, L.A., M.S.H, L.M.B, M.M.K., G.Z.; Supervision, L.A., M.S.H., P.R.; Project Administration, L.A., M.S.H., P.R.; Funding Acquisition, P.R., M.S.H., L.A.

## Abbreviations

(APCs): Antigen-presenting cells,
(CNI): calcineurin inhibitors,
(CITE-seq): cellular indexing of transcriptomes and epitopes sequencing,
(Tcon): conventional T-cells,
(CsA): cyclosporine A,
(TSDR): Treg-specific demethylation region,
(Teff): effector T-cells,
(fig.): figure,
(FACS): fluorescently activated cell sorting,
(FKBP12): FK506 binding protein,
(GBP2): gyanylate binding protein-2,
(ICE): inference-of-CRISPR-Edits,
(IL): interleukin,
(KO): knock-out,
(MFI): mean fluorescence intensity,
(nLC): nano-liquid chromatography,
(PBMC): peripheral blood mononuclear cells,
(PMA): phorbol-12-myristate-13-acetate,
(PAM): Protospacer Adjacent Motif,
(RNP): ribonucleoprotein,
(SOT): solid organ transplant,
(suppl.): supplementary,
(TH2): T helper 2.
(tac): tacrolimus,
(TNFα): tumor necrosis factor-alpha
(WT): unedited,
(Treg): regulatory T-cell

## Acknowledgements

We thank Anne Schulze for technical assistance in DNA methylation analysis and Geoffroy Andrieux for bioinformatic CASTSeq analyses. Furthermore, we thank Grit Nebrich for sample preparation (BIH-Core Unit Imaging Mass Spectrometry). In addition, we thank Dr. Harald Stachelscheid and Dr. Valeria Vallone Fernandez from the Core Unit for Stem Cells and Organoids (CUSCO) at the Berlin Institute of Health for access to their devices and technical support.

