## Supplemental files for "Clinical development of gene edited tacrolimus-resistant Treg (FKBP12^KO^-Treg) to enable simultaneous immunosuppression and support of immune regulation"

**Suppl. Table 1: List of general antibodies**

| Isolation and culture of Treg | Antibody | Fluorescence label | clone |
|---|---|---|---|
| | anti-CD4 | Vioblue | MT466 |
| | anti-CD25 | PE | 3G10 |
| | anti-CD127 | APC | MB1518C9 |
| **Phenotypic and functional assessment** | **Antibody** | **Fluorescence label** | **clone** |
| | anti-CD25 | PE | MA251 |
| | anti-CD127 | APC | A019D5 |
| | LIVE/DEAD™ | UV | - |
| | anti-CD3 | BV650 | OKT3 |
| | anti-CD8 | BV510 | RPAT8 |
| | anti-CD4 | PerCP Cy5.5 | SK3 |
| | anti-FOXP3 | AF488 | 259D |
| | anti-IL2 | PE-Cy7 | MQ117H12 |
| | anti-TNFa | Alexa 700 | MAb11 |
| | anti-CD137 | PE-Cy5 | 4B41 |
| | anti-CD154 | BV711 | 2431 |
| | anti-IFNg | eFluor 450 | 4S.B3 |
| | anti-CTLA4 | PE-Cy5 | BNI3 |
| | anti-PD1 | PE-Cy7 | EH12.2H7 |

### Suppl. Table 2: List of materials and instruments

| | | | | |
|---|---|---|---|---|
| **Isolation and culture of Treg** | **Material/Instrument** | **Company** | **Concentration** | **Additional information** |
| | EasySep™ Human CD4+ T-cell Enrichment Kit | Stemcell Technologies | - | |
| | CliniMACS Plus device | Miltenyi Biotec | - | |
| | anti-CD3/CD28-MACSiBeads | Miltenyi Biotec | - | |
| | X-Vivo-15 | Lonza | - | |
| | fetal calf serum (FCS) | Biochrom | 0,1ml/ml | |
| | penicillin/streptomycin | Biochrom | 0,01ml/ml | |
| | rapamycin | Sigma Aldrich | 1ul/ml | |
| | recombinant human IL-2 | Miltenyi Biotec | 1,250 IU/ml | |
| **FKBP12 knock-out in Treg** | **Material/Insturument** | **Company** | **Concentration** | |
| | HiFi Cas9 Nuclease V3 | IDT | 10ug | |
| | 2O'-methyl-3'phosphothioate-modified sgRNA | Synthego Corporation | 15ug | Sequence: 5'-G*G*G*CGCACCUUCCCCAAGCG + mod-scaffold |
| | Amaxa-P3-primary-cell-4D-Nucleofector-X-Kit-l | ThermoFisher | - | |
| | Amaxa-Nucleofector-4D | Lonza | - | program CO-115 |
| **Phenotypic and functional assessment** | **Material/Instrument** | **Company** | **Concentration** | **Additional information** |
| | PMA | | - | |
| | Ionomycin | | - | |
| | FoxP3/Transcription Factor Staining Buffer kit | eBioscience | - | |
| | Fortessa | BD | - | |
| | Cytoflex LC | Beckman Coulter | - | |
| **Methylation profiling** | **Material/Insturument** | **Company** | **Concentration** | |
| | Infinium-Methylation-EPIC-Kit | Illumina-EPIC-8-BeadCl | - | |
| | EZ-DNA-Methylation-Gold-Kit | Zymo Research | - | |
| | QIAquick-PCR-Purification-Kit | Qiagen | - | |
| **Cellular indexing of transcriptomes and epitopes sequencing (CITE-seq)** | **Material/Insturument** | **Company** | **Concentration** | |
| | Single-Index-Kit-T-Set-A/Single-Index-Kit-N-Se | 10X Genomics | - | |
| | High-Sensitivity-DNA-Kit | Agilent | - | |
| | NextSeq500 device | Illumina | - | |

**Suppl. Table 3: List of CITEseq antibodies**

| REAGENT or RESOURCE | SOURCE | IDENTIFIER |
|---|---|---|
| Antibodies | | |
| TotalSeq™-C0034 anti-human CD3 Antibody (UCHT1) | BioLegend | Cat# 300479 |
| TotalSeq™-C0045 anti-human CD4 Antibody (SK3) | BioLegend | Cat# 344651 |
| TotalSeq™-C0080 anti-human CD8a Antibody (RPA-T8) | BioLegend | Cat# 301071 |
| TotalSeq™-C0185 anti-human CD11a (TS2/4) | BioLegend | Cat# 350617 |
| TotalSeq™-C0085 anti-human CD25 (BC96) | BioLegend | Cat# 302649 |
| TotalSeq™-C0386 anti-human CD28 (CD28.2) | BioLegend | Cat# 302963 |
| TotalSeq™-C0176 anti-human CD39 (A1) | BioLegend | Cat# 328237 |
| TotalSeq™-C0576 anti-human CD49d (9F10) | BioLegend | Cat# 304345 |
| TotalSeq™-C0174 anti-human CD58 (TS2/9) | BioLegend | Cat# 330921 |
| TotalSeq™-C0146 anti-human CD69 (FN50) | BioLegend | Cat# 310951 |
| TotalSeq™-C0577 anti-human CD73 (AD2) | BioLegend | Cat# 344031 |
| TotalSeq™-C0156 anti-human CD95 (DX2) | BioLegend | Cat# 305651 |
| TotalSeq™-C0155 anti-human CD107a (H4A3) | BioLegend | Cat# 328649 |
| TotalSeq™-C0246 anti-human CD122 (TU27) | BioLegend | Cat# 339021 |
| TotalSeq™-C0390 anti-human CD127 (A019D5) | BioLegend | Cat# 351352 |
| TotalSeq™-C0355 anti-human CD137 (4B4-1) | BioLegend | Cat# 309839 |
| TotalSeq™-C0151 anti-human CD152 (BNI3) | BioLegend | Cat# 369621 |
| TotalSeq™-C0032 anti-human CD154 (24-31) | BioLegend | Cat# 310849 |
| TotalSeq™-C0140 anti-human CD183 (G025H7) | BioLegend | Cat# 353747 |
| TotalSeq™-C0144 anti-human CD185 (J252D4) | BioLegend | Cat# 356939 |
| TotalSeq™-C0071 anti-human CD194 (L291H4) | BioLegend | Cat# 359425 |
| TotalSeq™-C0141 anti-human CD195 (J418F1) | BioLegend | Cat# 359137 |
| TotalSeq™-C0143 anti-human CD196 (G034E3) | BioLegend | Cat# 353440 |
| TotalSeq™-C0148 anti-human CD197 (G043H7) | BioLegend | Cat# 353251 |
| TotalSeq™-C0152 anti-human CD223 (11C3C65) | BioLegend | Cat# 369335 |
| TotalSeq™-C0171 anti-human CD278 (C398.4A) | BioLegend | Cat# 313553 |
| TotalSeq™-C0088 anti-human CD279 (EH12.2H7) | BioLegend | Cat# 329963 |
| TotalSeq™-C0169 anti-human CD366 (F38-2E2) | BioLegend | Cat# 345049 |
| TotalSeq™-C0063 anti-human CD45RA (HI100) | BioLegend | Cat# 304163 |
| TotalSeq™-C0087 anti-human CD45RO (UCHL1) | BioLegend | Cat# 304259 |
| TotalSeq™-C0159 anti-human HLA-DR (L243) | BioLegend | Cat# 307663 |
| TotalSeq™-C0089 anti-human TIGIT (A15153G) | BioLegend | Cat# 372729 |

### Suppl. Table 4: Database comparison of differentially expressed CITEseq genes in FKBP12$^{KO}$-Treg and Treg$^{WT}$

| Category | Term | Count | % | PValue | Genes | List Total | Pop Hits | Pop Total | Fold Enrichment | Bonferroni | Benjamini | FDR |
|---|---|---|---|---|---|---|---|---|---|---|---|---|
| KEGG_PATHWAY | hsa04060:Cytokine-cytokine receptor interaction | 10 | 7.692.307.692.307.690 | 4,85E+02 | IL10, IL4, IL22, IL3, CXCL10, IL5, IL23R, IL13, CXCR5, CCL1 | 11 | 298 | 8840 | 26.967.663.209.273.900 | 2,23E+05 | 2,23E+05 | 1,55E+04 |
| INTERPRO | IPR009079:4_helix_cytokine-like_core | 6 | 4.615.384.615.384.610 | 1,06E+06 | IL10, IL4, IL22, IL3, IL5, IL13 | 13 | 57 | 20607 | 1.668.582.995.951.410 | 3,80E+06 | 3,80E+06 | 3,80E+06 |
| UP_KW_MOLECULAR_FUNCTION | KW-0202~Cytokine | 8 | 6.153.846.153.846.150 | 2,20E+05 | IL10, IL4, IL22, IL3, CXCL10, IL5, IL13, CCL1 | 13 | 196 | 11889 | 37.328.100.470.957.600 | 1,98E+07 | 1,98E+05 | 1,98E+05 |
| KEGG_PATHWAY | hsa05321:Inflammatory bowel disease | 6 | 4.615.384.615.384.610 | 4,50E+05 | IL10, IL4, IL22, IL5, IL23R, IL13 | 11 | 65 | 8840 | 7.418.181.818.181.810 | 2,07E+09 | 1,04E+09 | 7,20E+07 |
| KEGG_PATHWAY | hsa04630:JAK-STAT signaling pathway | 7 | 5.384.615.384.615.380 | 8,50E+06 | IL10, IL4, IL22, IL3, IL5, IL23R, IL13 | 11 | 168 | 8840 | 33.484.848.484.848.400 | 3,91E+07 | 1,30E+09 | 9,07E+06 |
| KEGG_PATHWAY | hsa05310:Asthma | 5 | 3.846.153.846.153.840 | 2,56E+08 | IL10, IL4, IL3, IL5, IL13 | 11 | 31 | 8840 | 12.961.876.832.844.500 | 1,18E+10 | 2,94E+08 | 2,05E+09 |
| GOTERM_MF_DIRECT | GO:0005125~cytokine activity | 6 | 4.615.384.615.384.610 | 7,47E+07 | IL10, IL4, IL22, IL3, IL5, IL13 | 13 | 195 | 19304 | 4.568.994.082.840.230 | 3,89E+09 | 3,89E+09 | 3,89E+09 |
| REACTOME_PATHWAY | R-HSA-449147~Signaling by Interleukins | 8 | 6.153.846.153.846.150 | 1,61E+09 | IL10, IL4, IL22, IL3, CXCL10, IL5, IL23R, IL13 | 13 | 473 | 11091 | 14.429.663.359.895.900 | 1,18E+11 | 1,18E+10 | 1,03E+10 |
| GOTERM_CC_DIRECT | GO:0005576~extracellular region | 10 | 7.692.307.692.307.690 | 1,72E+09 | IL10, IL4, IL22, IL3, CXCL10, IL5, CD160, IL13, IGF2, CCL1 | 13 | 2101 | 20887 | 7.647.274.191.776.810 | 3,10E+09 | 3,10E+10 | 3,10E+10 |
| BIOCARTA | h_cytokinePathway:Cytokine Network | 5 | 3.846.153.846.153.840 | 3,74E+09 | IL10, IL4, IL3, IL5, IL13 | 7 | 22 | 1623 | 5.269.480.519.480.510 | 7,86E+09 | 3,93E+10 | 3,93E+10 |
| BIOCARTA | h_dcPathway:Dendritic cells in regulating TH1 and TH2 Development | 5 | 3.846.153.846.153.840 | 3,74E+09 | IL10, IL4, IL3, IL5, IL13 | 7 | 22 | 1623 | 5.269.480.519.480.510 | 7,86E+09 | 3,93E+10 | 3,93E+10 |
| UP_KW_CELLULAR_COMPONENT | KW-0964~Secreted | 10 | 7.692.307.692.307.690 | 8,86E+07 | IL10, IL4, IL22, IL3, CXCL10, IL5, CD160, IL13, IGF2, CCL1 | 13 | 2180 | 17942 | 6.330.980.945.659.840 | 4,43E+09 | 4,43E+09 | 4,43E+09 |
| BIOCARTA | h_inflamPathway:Cytokines and Inflammatory Response | 5 | 3.846.153.846.153.840 | 1,21E+10 | IL10, IL4, IL3, IL5, IL13 | 7 | 29 | 1623 | 39.975.369.458.128.000 | 2,53E+10 | 8,45E+09 | 8,45E+09 |
| GOTERM_MF_DIRECT | GO:0008083~growth factor activity | 5 | 3.846.153.846.153.840 | 2,42E+10 | IL10, IL4, IL3, IL5, IGF2 | 13 | 165 | 19304 | 44.997.668.997.669 | 1,26E+10 | 6,28E+09 | 6,28E+09 |
| GOTERM_CC_DIRECT | GO:0005615~extracellular space | 9 | 6.923.076.923.076.920 | 2,75E+09 | IL10, IL4, IL22, IL3, CXCL10, IL5, IL13, IGF2, CCL1 | 13 | 2033 | 20887 | 7.112.754.928.298.450 | 4,95E+11 | 2,47E+11 | 2,47E+11 |
| GOTERM_BP_DIRECT | GO:0006954~inflammatory response | 6 | 4.615.384.615.384.610 | 3,06E+09 | IL22, CXCL10, IL5, IL23R, IL13, CCL1 | 13 | 422 | 19734 | 21.582.938.388.625.500 | 8,40E+10 | 7,38E+11 | 6,84E+10 |
| REACTOME_PATHWAY | R-HSA-1280215~Cytokine Signaling in Immune system | 8 | 6.153.846.153.846.150 | 5,05E+09 | IL10, IL4, IL22, IL3, CXCL10, IL5, IL23R, IL13 | 13 | 786 | 11091 | 8.683.499.706.400.470 | 3,69E+12 | 1,84E+10 | 1,62E+12 |
| GOTERM_BP_DIRECT | GO:0006955~immune response | 6 | 4.615.384.615.384.610 | 7,34E+09 | IL10, IL4, IL3, IL5, IL13, CXCR5 | 13 | 505 | 19734 | 18.035.643.564.356.400 | 0.002017521835727676 | 7,38E+11 | 6,84E+10 |
| GOTERM_BP_DIRECT | GO:0042531~positive regulation of tyrosine phosphorylation of STAT protein | 4 | 3.076.923.076.923.070 | 8,05E+09 | IL4, IL3, IL23R, IL13 | 13 | 67 | 19734 | 906.268.656.716.418 | 0.002211635767839004 | 7,38E+11 | 6,84E+10 |
| REACTOME_PATHWAY | R-HSA-168256~Immune System | 10 | 7.692.307.692.307.690 | 3,23E+10 | IL10, FKBP1A, IL4, IL22, IL3, CXCL10, IL5, CD160, IL23R, IL13 | 13 | 2054 | 11091 | 4.153.621.451.576.660 | 0.0023527486272645337 | 7,85E+11 | 6,88E+11 |
| UP_SEQ_FEATURE | CARBOHYD:N-linked (GlcNAc...) asparagine | 10 | 7.692.307.692.307.690 | 3,49E+10 | IL10, IL4, IL22, IL3, IL5, CD160, IL23R, IL13, CXCR5, CCL1 | 12 | 4422 | 20569 | 38.762.626.262.626.200 | 0.0015700679246376525 | 0.001571274340042267 | 0.001571274340042267 |
| KEGG_PATHWAY | hsa04664:Fc epsilon RI signaling pathway | 4 | 3.076.923.076.923.070 | 5,25E+10 | IL4, IL3, IL5, IL13 | 11 | 69 | 8840 | 4.658.761.528.326.740 | 0.0024131673310502366 | 4,83E+12 | 3,36E+12 |
| GOTERM_BP_DIRECT | GO:0050728~negative regulation of inflammatory response | 4 | 3.076.923.076.923.070 | 8,48E+10 | IL10, IL4, IL22, IL13 | 13 | 147 | 19734 | 4.130.612.244.897.950 | 0.0230543374958041 | 0.005830814016344317 | 0.00540675481515564 |
| UP_KW_PTM | KW-1015~Disulfide bond | 11 | 8.461.538.461.538.460 | 8,84E+10 | IL10, IL4, IL22, IL3, CXCL10, IL5, CD160, IL13, CXCR5, IGF2, CCL1 | 13 | 3906 | 14252 | 30.874.000.551.419.900 | 7,07E+10 | 7,07E+10 | 7,07E+10 |
| GOTERM_BP_DIRECT | GO:0002639~positive regulation of immunoglobulin production | 3 | 23.076.923.076.923.000 | 1,18E+12 | IL10, IL5, IL13 | 13 | 27 | 19734 | 16.866.666.666.666.600 | 0.031925366083911055 | 0.006488835902186441 | 0.006016920563845609 |
| KEGG_PATHWAY | hsa04657:IL-17 signaling pathway | 4 | 3.076.923.076.923.070 | 1,37E+11 | IL4, CXCL10, IL5, IL13 | 11 | 95 | 8840 | 3.383.732.057.416.260 | 0.006265140941786318 | 9,15E+11 | 6,36E+11 |
| KEGG_PATHWAY | hsa05200:Pathways in cancer | 6 | 4.615.384.615.384.610 | 1,53E+11 | IL4, IL3, IL5, IL23R, IL13, IGF2 | 11 | 533 | 8840 | 9.046.563.192.904.650 | 0.007010567151745528 | 9,15E+11 | 6,36E+11 |
| KEGG_PATHWAY | hsa04061:Viral protein interaction with cytokine and cytokine receptor | 4 | 3.076.923.076.923.070 | 1,59E+12 | IL10, CXCL10, CXCR5, CCL1 | 11 | 100 | 8840 | 3.214.545.454.545.450 | 0.00729359127725604 | 9,15E+11 | 6,36E+11 |
| REACTOME_PATHWAY | R-HSA-6785807~Interleukin-4 and Interleukin-13 signaling | 4 | 3.076.923.076.923.070 | 1,85E+11 | IL10, IL4, IL23R, IL13 | 13 | 108 | 11091 | 31.598.290.598.290.500 | 0.01343866343017519 | 0.0033821314459422107 | 0.0029651563361685136 |
| BBID | 35.Chemokines_EAE_Autoreactive_T_Cells | 4 | 3.076.923.076.923.070 | 1,95E+12 | IL10, IL4, IL5, IL13 | 8 | 8 | 388 | 24.25 | 0.005642215555549668 | 0.005462549961342284 | 0.004292003541054652 |
| UP_KW_DOMAIN | KW-0732~Signal | 11 | 8.461.538.461.538.460 | 2,19E+12 | IL10, IL4, IL22, IL3, CXCL10, IL5, CD160, IL23R, IL13, IGF2, CCL1 | 13 | 4410 | 14597 | 28.007.500.436.071.800 | 0.0010942038551782218 | 0.0010946830826670478 | 0.0010946830826670478 |
| GOTERM_BP_DIRECT | GO:2000352~negative regulation of endothelial cell apoptotic process | 3 | 23.076.923.076.923.000 | 2,35E+12 | IL10, IL4, IL13 | 13 | 38 | 19734 | 1.198.421.052.631.570 | 0.06269576054053494 | 0.010789955427523316 | 0.010005231396430711 |
| UP_KW_MOLECULAR_FUNCTION | KW-0339~Growth factor | 4 | 3.076.923.076.923.070 | 2,74E+12 | IL4, IL3, IL5, IGF2 | 13 | 132 | 11889 | 27.713.286.713.286.700 | 0.0024590550887176876 | 0.0012308733957028485 | 0.0012308733957028485 |
| GOTERM_BP_DIRECT | GO:0030890~positive regulation of B cell proliferation | 3 | 23.076.923.076.923.000 | 3,46E+12 | IL4, IL5, IL13 | 13 | 46 | 19734 | 99.0 | 0.09069343829004739 | 0.013579508233352586 | 0.012591907634563308 |
| BBID | 88.Alternatively_Activated_APC | 4 | 3.076.923.076.923.070 | 5,61E+11 | IL10, IL4, IL3, IL13 | 8 | 11 | 388 | 17.636.363.636.363.600 | 0.016152287614736194 | 0.007859110790195738 | 0.006175015620868079 |
| UP_KW_PTM | KW-0325~Glycoprotein | 11 | 8.461.538.461.538.460 | 6,09E+11 | IL10, IL4, IL22, IL3, IL5, CD160, IL23R, IL13, CXCR5, IGF2, CCL1 | 13 | 4813 | 14252 | 2.505.585.833.240.100 | 0.0048651907378476356 | 0.002437788998845631 | 0.002437788998845631 |
| KEGG_PATHWAY | hsa05330:Allograft rejection | 3 | 23.076.923.076.923.000 | 7,92E+10 | IL10, IL4, IL5 | 11 | 38 | 8840 | 63.444.976.076.555.000 | 0.035804681662974014 | 0.004049660663206581 | 0.0028171552439697953 |
| GOTERM_BP_DIRECT | GO:0070098~chemokine-mediated signaling pathway | 3 | 23.076.923.076.923.000 | 8,00E+11 | CXCL10, CXCR5, CCL1 | 13 | 70 | 19734 | 6.505.714.285.714.280 | 0.1975579271550194 | 0.027500944617755253 | 0.025500875918282144 |
| SMART | SM00190:IL4_13 | 2 | 15.384.615.384.615.300 | 9,35E+09 | IL4, IL13 | 6 | 2 | 10692 | 1782.0 | 0.003735171745645305 | 0.00374041498983108 | 0.00374041498983108 |
| UP_KW_BIOLOGICAL_PROCESS | KW-0145~Chemotaxis | 2 | 15.384.615.384.615.300 | 0.052284623 | CXCL10, CCL1 | 7 | 102 | 11450 | 3.207.282.913.165.260 | 0.4155070575523566 | 0.261423116074253 | 0.261423116074253 |
| GOTERM_BP_DIRECT | GO:0030183~B cell differentiation | 2 | 15.384.615.384.615.300 | 0.053390275 | IL10, IL4 | 13 | 90 | 19734 | 33.733.333.333.333.300 | 0.9999997200925137 | 0.39681961168345536 | 0.36796000356102226 |
| GOTERM_BP_DIRECT | GO:0050829~defense response to Gram-negative bacterium | 2 | 15.384.615.384.615.300 | 0.056278332 | CD160, IL23R | 13 | 95 | 19734 | 31.957.894.736.842.100 | 0.9999998791964378 | 0.40727740740919105 | 0.37765723232488624 |
| GOTERM_BP_DIRECT | GO:0120162~positive regulation of cold-induced thermogenesis | 2 | 15.384.615.384.615.300 | 0.059733339 | IL4, IL13 | 13 | 101 | 19734 | 3.005.940.594.059.400 | 0.9999999559409309 | 0.42119662273979325 | 0.39056414108599014 |
| REACTOME_PATHWAY | R-HSA-447115~Interleukin-12 family signaling | 2 | 15.384.615.384.615.300 | 0.059986843 | IL10, IL23R | 13 | 57 | 11091 | 2.993.522.267.206.470 | 0.9890660509584789 | 0.29193597099144775 | 0.2559438649788035 |
| KEGG_PATHWAY | hsa04151:PI3K-Akt signaling pathway | 3 | 23.076.923.076.923.000 | 0.060546582 | IL4, IL3, IGF2 | 11 | 362 | 8840 | 6.659.969.864.389.750 | 0.9434722732923543 | 0.16383193012460376 | 0.11397003834755043 |
| UCSC_TFBS | CREL | 5 | 3.846.153.846.153.840 | 0.060733031 | IL10, IL4, CXCL10, IL13, IGF2 | 13 | 2703 | 20672 | 2.941.461.054.668.600 | 0.999977770614214 | 1.0 | 1.0 |
| GOTERM_BP_DIRECT | GO:0061844~antimicrobial humoral immune response mediated by antimicrobial peptide | 2 | 15.384.615.384.615.300 | 0.064321978 | CXCL10, CCL1 | 13 | 109 | 19734 | 27.853.211.009.174.300 | 0.9999999885246025 | 0.44221360440969154 | 0.4100526149980776 |
| GOTERM_BP_DIRECT | GO:0045893~positive regulation of DNA-templated transcription | 3 | 23.076.923.076.923.000 | 0.067699692 | IL10, IL4, IL5 | 13 | 713 | 19734 | 6.387.096.774.193.540 | 0.9999999957552459 | 0.4540833039495026 | 0.42105906366226603 |
| UP_SEQ_FEATURE | CARBOHYD:O-linked (GalNAc...) threonine | 2 | 15.384.615.384.615.300 | 0.070389986 | IL5, IGF2 | 12 | 136 | 20569 | 25.207.107.843.137.200 | 0.9625427671855729 | 1.0 | 1.0 |
| GOTERM_BP_DIRECT | GO:0006935~chemotaxis | 2 | 15.384.615.384.615.300 | 0.071166460 | CXCL10, CCL1 | 13 | 121 | 19734 | 25.090.909.090.909.000 | 0.9999999984762297 | 0.46597087111755625 | 0.4320820804908249 |
| GOTERM_BP_DIRECT | GO:0016525~negative regulation of angiogenesis | 2 | 15.384.615.384.615.300 | 0.079093666 | CXCL10, CD160 | 13 | 135 | 19734 | 2.248.888.888.888.880 | 0.999999999855705 | 0.5058315896041464 | 0.46904383763293583 |
| GOTERM_BP_DIRECT | GO:0032496~response to lipopolysaccharide | 2 | 15.384.615.384.615.300 | 0.081347164 | IL23R, IL13 | 13 | 139 | 19734 | 2.184.172.661.870.500 | 0.9999999999264402 | 0.5084197775147989 | 0.4714437936955409 |
| KEGG_PATHWAY | hsa05140:Leishmaniasis | 2 | 15.384.615.384.615.300 | 0.083808988 | IL10, IL4 | 11 | 77 | 8840 | 20.873.671.782.762.600 | 0.9821615450796923 | 0.21417852543354188 | 0.1489937568233335 |
| BIOGRID_INTERACTION | 2:alpha-2-macroglobulin(A2M) | 2 | 15.384.615.384.615.300 | 0.084808717 | IL10, IL4 | 13 | 145 | 19712 | 20.914.588.859.416.400 | 0.9999999995559974 | 1.0 | 1.0 |
| REACTOME_PATHWAY | R-HSA-5663202~Diseases of signal transduction by growth factor receptors and second messengers | 3 | 23.076.923.076.923.000 | 0.085757231 | IL10, FKBP1A, IL22 | 13 | 459 | 11091 | 5.576.168.929.110.100 | 0.9985628835000173 | 0.3618458020778368 | 0.31723467579426784 |
| BBID | 33.GProt-coupled_Rec_T_Cell_med_Inflamm | 2 | 15.384.615.384.615.300 | 0.087445138 | CXCL10, CCL1 | 8 | 5 | 388 | 19.400.000.000.000.000 | 0.9296100735946986 | 0.24295526909003207 | 0.1908934257135966 |
| BBID | 34.Chemokines_in_EAE | 2 | 15.384.615.384.615.300 | 0.087445138 | CXCL10, CCL1 | 8 | 5 | 388 | 19.400.000.000.000.000 | 0.9296100735946986 | 0.24295526909003207 | 0.1908934257135966 |
| REACTOME_PATHWAY | R-HSA-500792~GPCR ligand binding | 3 | 23.076.923.076.923.000 | 0.088678669 | CXCL10, CXCR5, CCL1 | 13 | 468 | 11091 | 5.468.934.911.242.600 | 0.9988623148702614 | 0.3618458020778368 | 0.31723467579426784 |
| GOTERM_BP_DIRECT | GO:0007204~positive regulation of cytosolic calcium ion concentration | 2 | 15.384.615.384.615.300 | 0.090868249 | CXCR5, CCL1 | 13 | 156 | 19734 | 19.461.538.461.538.400 | 0.9999999999958085 | 0.5553059666084488 | 0.5149200781278344 |
| REACTOME_PATHWAY | R-HSA-9700206~Signaling by ALK in cancer | 2 | 15.384.615.384.615.300 | 0.094179044 | IL10, IL22 | 13 | 91 | 11091 | 1.875.063.398.140.320 | 0.9992687051025686 | 0.3618458020778368 | 0.31723467579426784 |
| REACTOME_PATHWAY | R-HSA-9725370~Signaling by ALK fusions and activated point mutants | 2 | 15.384.615.384.615.300 | 0.094179044 | IL10, IL22 | 13 | 91 | 11091 | 1.875.063.398.140.320 | 0.9992687051025686 | 0.3618458020778368 | 0.31723467579426784 |
| UP_KW_BIOLOGICAL_PROCESS | KW-0395~Inflammatory response | 2 | 15.384.615.384.615.300 | 0.097950821 | CXCL10, IL23R | 7 | 195 | 11450 | 16.776.556.776.556.700 | 0.6433007842250972 | 0.32650273752293685 | 0.32650273752293685 |
| BIOCARTA | h_nktPathway:Selective expression of chemokine receptors during T-cell polarization | 2 | 15.384.615.384.615.300 | 0.099295396 | IL4, IL5 | 7 | 28 | 1623 | 1.656.122.448.979.590 | 0.8887679266409064 | 0.23168925814668317 | 0.23168925814668317 |

| | | | | | | | | | | | | |
|---|---|---|---|---|---|---|---|---|---|---|---|---|
| GOTERM_BP_DIRECT | GO:0050731~positive regulation of peptidyl-tyrosine phosphorylation | 3 | 23.076.923.076.923.000 | 0.0011493737195443503 | IL3, IL5, IGF2 | 13 | 84 | 19734 | 5.421.428.571.428.570 | 0.27112975843296017 | 0.03511975254163293 | 0.032565588720423255 |
| INTERPRO | IPR018096:IL-4/IL-13_CS | 2 | 15.384.615.384.615.300 | 0.0011643419274750427 | IL4, IL13 | 13 | 2 | 20607 | 15.851.538.461.538.400 | 0.04107338630403168 | 0.013972103129700512 | 0.013972103129700512 |
| INTERPRO | IPR001325:IL-4/IL-13 | 2 | 15.384.615.384.615.300 | 0.0011643419274750427 | IL4, IL13 | 13 | 2 | 20607 | 15.851.538.461.538.400 | 0.04107338630403168 | 0.013972103129700512 | 0.013972103129700512 |
| BIOCARTA | h_stemPathway:Regulation of hematopoiesis by cytokines | 3 | 23.076.923.076.923.000 | 0.0011712009494656186 | IL4, IL3, IL5 | 7 | 15 | 1623 | 4.637.142.857.142.850 | 0.02430928598420745 | 0.005612485533047828 | 0.005612485533047828 |
| KEGG_PATHWAY | hsa04672:Intestinal immune network for IgA production | 3 | 23.076.923.076.923.000 | 0.001316650598107268 | IL10, IL4, IL5 | 11 | 49 | 8840 | 4.920.222.634.508.340 | 0.05880584718622017 | 0.006056592751293432 | 0.004213281913943258 |
| BIOCARTA | h_gata3Pathway:GATA3 participate in activating the Th2 cytokine genes expression | 3 | 23.076.923.076.923.000 | 0.0013363060792971019 | IL4, IL5, IL13 | 7 | 16 | 1623 | 43.473.214.285.714.200 | 0.027690582466258995 | 0.0056124855330478285 | 0.0056124855330478285 |
| KEGG_PATHWAY | hsa05320:Autoimmune thyroid disease | 3 | 23.076.923.076.923.000 | 0.0015390876918740237 | IL10, IL4, IL5 | 11 | 53 | 8840 | 45.488.850.771.869.600 | 0.06840077417046875 | 0.006436184893291372 | 0.004477346012724433 |
| GOTERM_CC_DIRECT | GO:0009897~external side of plasma membrane | 4 | 3.076.923.076.923.070 | 0.0016488094787702222 | CXCL10, IL23R, IL13, CXCR5 | 13 | 429 | 20887 | 14.980.814.057.737.100 | 0.029266264129145014 | 0.009892856872621333 | 0.009892856872621333 |
| REACTOME_PATHWAY | R-HSA-380108~Chemokine receptors bind chemokines | 3 | 23.076.923.076.923.000 | 0.0017743449986740763 | CXCL10, CXCR5, CCL1 | 13 | 59 | 11091 | 4.338.070.404.172.090 | 0.12159036016063762 | 0.025905436980641513 | 0.022711615983028177 |
| GOTERM_BP_DIRECT | GO:1903660~negative regulation of complement-dependent cytotoxicity | 2 | 15.384.615.384.615.300 | 0.001823245945208361 | IL4, IL13 | 13 | 3 | 19734 | 1012.0 | 0.3945905495860319 | 0.050139263493229926 | 0.046492771602813204 |
| BBID | 97.Immune_injury_MS-lesions_MS_antigen | 4 | 3.076.923.076.923.070 | 0.0018310303061562134 | IL10, IL4, IL5, IL13 | 8 | 16 | 388 | 12.125 | 0.05176086084520992 | 0.017089616190791326 | 0.013427555578478897 |
| GOTERM_BP_DIRECT | GO:0045191~regulation of isotype switching | 2 | 15.384.615.384.615.300 | 0.0024303171048020863 | IL10, IL4 | 13 | 4 | 19734 | 7.590.000.000.000.000 | 0.48785662186446155 | 0.06075792762005216 | 0.05633916924768473 |
| GOTERM_BP_DIRECT | GO:0008284~positive regulation of cell population proliferation | 4 | 3.076.923.076.923.070 | 0.003102043656139053 | IL4, IL3, CXCL10, IGF2 | 13 | 506 | 19734 | 12.000.000.000.000.000 | 0.57445639886133 | 0.07108850045318663 | 0.06591842769295488 |
| INTERPRO | IPR020423:IL-10_CS | 2 | 15.384.615.384.615.300 | 0.003489298473354292 | IL10, IL22 | 13 | 6 | 20607 | 5.283.846.153.846.150 | 0.11823915438777954 | 0.03140368626018863 | 0.03140368626018863 |
| DRUGBANK | DB16318~Romilkimab | 2 | 15.384.615.384.615.300 | 0.0040329316472629845 | IL4, IL13 | 7 | 2 | 2973 | 42.471.428.571.428.500 | 0.13189100735925285 | 0.10577540923890473 | 0.10577540923890473 |
| KEGG_PATHWAY | hsa04658:Th1 and Th2 cell differentiation | 3 | 23.076.923.076.923.000 | 0.004566497271140984 | IL4, IL5, IL13 | 11 | 92 | 8840 | 26.205.533.596.837.900 | 0.189853315910406 | 0.01750490620604044 | 0.012177326056375956 |
| GOTERM_BP_DIRECT | GO:0042092~type 2 immune response | 2 | 15.384.615.384.615.300 | 0.004855218874646386 | IL10, IL4 | 13 | 8 | 19734 | 37.950.000.000.000.000 | 0.7377447541217337 | 0.10270655311751971 | 0.09523698561806374 |
| KEGG_PATHWAY | hsa04640:Hematopoietic cell lineage | 3 | 23.076.923.076.923.000 | 0.005269613414394301 | IL4, IL3, IL5 | 11 | 99 | 8840 | 24.352.617.079.889.800 | 0.215762328928767 | 0.01864632438939522 | 0.012971356096970588 |
| BBID | 113.Th1andTh2cells | 3 | 23.076.923.076.923.000 | 0.005624196814673069 | IL10, IL4, IL13 | 8 | 7 | 388 | 2.078.571.428.571.420 | 0.15088622639085147 | 0.03149550216216918 | 0.024746465984561507 |
| BBID | 89.CD4_T_Cell_Differentiation | 3 | 23.076.923.076.923.000 | 0.005624196814673069 | IL4, IL5, IL13 | 8 | 7 | 388 | 2.078.571.428.571.420 | 0.15088622639085147 | 0.03149550216216918 | 0.024746465984561507 |
| DRUGBANK | DB12159~Dupilumab | 2 | 15.384.615.384.615.300 | 0.0060443090993659845 | IL4, IL13 | 7 | 3 | 2973 | 2.831.428.571.428.570 | 0.19119121189438082 | 0.10577540923890473 | 0.10577540923890473 |
| KEGG_PATHWAY | hsa04659:Th17 cell differentiation | 3 | 23.076.923.076.923.000 | 0.006242573683948696 | IL4, IL22, IL23R | 11 | 108 | 8840 | 2.232.323.232.323.230 | 0.25028222320602633 | 0.020511313532974286 | 0.01426873984902559 |
| UP_KW_BIOLOGICAL_PROCESS | KW-0075~B-cell activation | 2 | 15.384.615.384.615.300 | 0.006273123181982339 | IL4, CXCR5 | 7 | 12 | 11450 | 2.726.190.476.190.470 | 0.060989688950836896 | 0.06273123181982339 | 0.06273123181982339 |
| GOTERM_BP_DIRECT | GO:0043306~positive regulation of mast cell degranulation | 2 | 15.384.615.384.615.300 | 0.0066703479896124155 | IL4, IL13 | 13 | 11 | 19734 | 276.0 | 0.8412605184814952 | 0.13102469265310102 | 0.12149562409651186 |
| BBID | 80.T_cell_Activation | 4 | 3.076.923.076.923.070 | 0.006995795064845473 | IL10, IL4, IL5, IL13 | 8 | 25 | 388 | 7.760.000.000.000.000 | 0.18420406152740276 | 0.03264704363594554 | 0.025651248571100065 |
| KEGG_PATHWAY | hsa04660:T cell receptor signaling pathway | 3 | 23.076.923.076.923.000 | 0.007907293601836214 | IL10, IL4, IL5 | 11 | 122 | 8840 | 1.976.154.992.548.430 | 0.3059291355550534 | 0.024249033712297724 | 0.016868893071725059 |
| GOTERM_BP_DIRECT | GO:0045348~positive regulation of MHC class II biosynthetic process | 2 | 15.384.615.384.615.300 | 0.008482441772482712 | IL10, IL4 | 13 | 14 | 19734 | 21.685.714.285.714.200 | 0.9039245220201289 | 0.1457919679645466 | 0.1351889157489432 |
| GOTERM_BP_DIRECT | GO:0030595~leukocyte chemotaxis | 2 | 15.384.615.384.615.300 | 0.008482441772482712 | IL10, CXCR5 | 13 | 14 | 19734 | 21.685.714.285.714.200 | 0.9039245220201289 | 0.1457919679645466 | 0.1351889157489432 |
| REACTOME_PATHWAY | R-HSA-9012546~Interleukin-18 signaling | 2 | 15.384.615.384.615.300 | 0.008625671919673871 | IL4, IL13 | 13 | 8 | 11091 | 21.328.846.153.846.100 | 0.4686877818761561 | 0.10494567502269876 | 0.09200716714318796 |
| GOTERM_BP_DIRECT | GO:0090026~positive regulation of monocyte chemotaxis | 2 | 15.384.615.384.615.300 | 0.012097541795365097 | CXCL10, CCL1 | 13 | 20 | 19734 | 151.8 | 0.9648141396237214 | 0.18482355520696675 | 0.17138184210100552 |
| GOTERM_BP_DIRECT | GO:0043032~positive regulation of macrophage activation | 2 | 15.384.615.384.615.300 | 0.012097541795365097 | IL10, IL13 | 13 | 20 | 19734 | 151.8 | 0.9648141396237214 | 0.18482355520696675 | 0.17138184210100552 |
| GOTERM_BP_DIRECT | GO:0042116~macrophage activation | 2 | 15.384.615.384.615.300 | 0.013299887534963603 | IL4, IL13 | 13 | 22 | 19734 | 138.0 | 0.9748276438047009 | 0.19249837221657845 | 0.1784984906008273 |
| BIOGRID_INTERACTION | 6373:C-X-C motif chemokine ligand 11(CXCL11) | 2 | 15.384.615.384.615.300 | 0.013915976112981482 | CXCL10, CCL1 | 13 | 23 | 19712 | 13.185.284.280.936.400 | 0.9668041959131123 | 1.0 | 1.0 |
| GOTERM_BP_DIRECT | GO:0032740~positive regulation of interleukin-17 production | 2 | 15.384.615.384.615.300 | 0.01629988567027995 | IL23R, CCL1 | 13 | 27 | 19734 | 11.244.444.444.444.400 | 0.9891044661886887 | 0.21871927511290795 | 0.20281241874106012 |
| BBID | 22.Cytokine-chemokine_CNS | 3 | 23.076.923.076.923.000 | 0.0169215796248871 | IL10, IL4, IL3 | 8 | 12 | 388 | 12.125 | 0.39038348153718627 | 0.0676863184995484 | 0.053182107392502315 |
| GOTERM_BP_DIRECT | GO:0001774~microglial cell activation | 2 | 15.384.615.384.615.300 | 0.017497542009032637 | IL4, IL13 | 13 | 29 | 19734 | 10.468.965.517.241.300 | 0.9922061413206921 | 0.21871927511290795 | 0.20281241874106012 |
| GOTERM_BP_DIRECT | GO:0042104~positive regulation of activated T cell proliferation | 2 | 15.384.615.384.615.300 | 0.017497542009032637 | IL23R, IGF2 | 13 | 29 | 19734 | 10.468.965.517.241.300 | 0.9922061413206921 | 0.21871927511290795 | 0.20281241874106012 |
| GOTERM_BP_DIRECT | GO:0051281~positive regulation of release of sequestered calcium ion into cytosol | 2 | 15.384.615.384.615.300 | 0.01869386157489154 | CXCL10, IL13 | 13 | 31 | 19734 | 9.793.548.387.096.770 | 0.9944250398338118 | 0.22351356230848582 | 0.2072580305042323 |
| KEGG_PATHWAY | hsa04062:Chemokine signaling pathway | 3 | 23.076.923.076.923.000 | 0.019015456796674147 | CXCL10, CXCR5, CCL1 | 11 | 193 | 8840 | 12.491.756.947.715.400 | 0.5865151825718107 | 0.054669438290438174 | 0.038030913593348294 |
| REACTOME_PATHWAY | R-HSA-375276~Peptide ligand-binding receptors | 3 | 23.076.923.076.923.000 | 0.01913996259209226 | CXCL10, CXCR5, CCL1 | 13 | 201 | 11091 | 12.733.639.494.833.500 | 0.7560426142940163 | 0.1996024670318193 | 0.17499394369912924 |
| BBID | 50.Hypothesis_of_bronchial_asthma | 3 | 23.076.923.076.923.000 | 0.01982409815913659 | IL4, IL5, IL13 | 8 | 13 | 388 | 11.192.307.692.307.600 | 0.44047872153966683 | 0.06938434355697806 | 0.05451626993762562 |
| SMART | SM00199:SCY | 2 | 15.384.615.384.615.300 | 0.020411263268639054 | CXCL10, CCL1 | 6 | 44 | 10692 | 8.099.999.999.999.990 | 0.07917917642820205 | 0.04082252653727811 | 0.04082252653727811 |
| GOTERM_BP_DIRECT | GO:0050776~regulation of immune response | 2 | 15.384.615.384.615.300 | 0.021082495818646057 | FKBP1A, IL4 | 13 | 35 | 19734 | 8.674.285.714.285.710 | 0.9971478266128886 | 0.22929522227497717 | 0.2126192061095243 |
| GOTERM_BP_DIRECT | GO:0002230~positive regulation of defense response to virus by host | 2 | 15.384.615.384.615.300 | 0.021082495818646057 | IL4, IL23R | 13 | 35 | 19734 | 8.674.285.714.285.710 | 0.9971478266128886 | 0.22929522227497717 | 0.2126192061095243 |
| GOTERM_BP_DIRECT | GO:0042113~B cell activation | 2 | 15.384.615.384.615.300 | 0.02167882101508875 | IL4, CXCR5 | 13 | 36 | 19734 | 8.433.333.333.333.330 | 0.997587872696435 | 0.22929522227497717 | 0.2126192061095243 |
| INTERPRO | IPR039809:Chemokine_b/g/d | 2 | 15.384.615.384.615.300 | 0.024761181693129814 | CXCL10, CCL1 | 13 | 43 | 20607 | 7.372.808.586.762.070 | 0.5944958862033065 | 0.13904092264134646 | 0.13904092264134646 |
| GOTERM_BP_DIRECT | GO:0032733~positive regulation of interleukin-10 production | 2 | 15.384.615.384.615.300 | 0.025249784957361826 | IL4, IL13 | 13 | 42 | 19734 | 7.228.571.428.571.420 | 0.9991175940872095 | 0.2538228653262671 | 0.23536302057526587 |
| GOTERM_BP_DIRECT | GO:0046427~positive regulation of receptor signaling pathway via JAK-STAT | 2 | 15.384.615.384.615.300 | 0.02584378265140174 | IL10, IL5 | 13 | 43 | 19734 | 706.046.511.627.907 | 0.9992537800888601 | 0.2538228653262671 | 0.23536302057526587 |
| BIOGRID_INTERACTION | 3566:interleukin 4 receptor(IL4R) | 2 | 15.384.615.384.615.300 | 0.025872288846690183 | IL4, IL13 | 13 | 43 | 19712 | 7.052.593.917.710.190 | 0.9982873713525411 | 1.0 | 1.0 |
| INTERPRO | IPR036048:Interleukin_8-like_sf | 2 | 15.384.615.384.615.300 | 0.027035734958039587 | CXCL10, CCL1 | 13 | 47 | 20607 | 6.745.335.515.548.280 | 0.6271895873607103 | 0.13904092264134646 | 0.13904092264134646 |
| INTERPRO | IPR001811:Chemokine_IL8-like_dom | 2 | 15.384.615.384.615.300 | 0.027035734958039587 | CXCL10, CCL1 | 13 | 47 | 20607 | 6.745.335.515.548.280 | 0.6271895873607103 | 0.13904092264134646 | 0.13904092264134646 |
| REACTOME_PATHWAY | R-HSA-912526~Interleukin receptor SHC signaling | 2 | 15.384.615.384.615.300 | 0.02883900693023833 | IL3, IL5 | 13 | 27 | 11091 | 6.319.658.119.658.110 | 0.8818972993133898 | 0.26315593823842476 | 0.23071205544190665 |
| BIOCARTA | h_eosinophilsPathway:The Role of Eosinophils in the Chemokine Network of Allergy | 2 | 15.384.615.384.615.300 | 0.029257344903427008 | IL3, IL5 | 7 | 8 | 1623 | 5.796.428.571.428.570 | 0.4639733553067985 | 0.10240070716199452 | 0.10240070716199452 |
| GOTERM_BP_DIRECT | GO:0051384~response to glucocorticoid | 2 | 15.384.615.384.615.300 | 0.029992484826157798 | IL10, IL22 | 13 | 50 | 19734 | 60.72 | 0.9997692444138082 | 0.2844114940411515 | 0.2637270217472496 |
| GOTERM_MF_DIRECT | GO:0008009~chemokine activity | 2 | 15.384.615.384.615.300 | 0.03125541419426438 | CXCL10, CCL1 | 13 | 51 | 19304 | 5.823.227.752.639.510 | 0.8081848120251579 | 0.5417605127005825 | 0.5417605127005825 |
| GOTERM_BP_DIRECT | GO:0007165~signal transduction | 4 | 3.076.923.076.923.070 | 0.035512480915770694 | IL22, CXCL10, IL13, CCL1 | 13 | 1240 | 19734 | 4.896.774.193.548.380 | 0.9999519638367415 | 0.3058393064034729 | 0.2835964477559476 |
| GOTERM_BP_DIRECT | GO:0042110~T cell activation | 2 | 15.384.615.384.615.300 | 0.03589111142904025 | FKBP1A, IL4 | 13 | 60 | 19734 | 5.060.000.000.000.000 | 0.9999568804835458 | 0.3058393064034729 | 0.2835964477559476 |
| BIOCARTA | h_il5Pathway:IL 5 Signaling Pathway | 2 | 15.384.615.384.615.300 | 0.0364591209512846 | IL4, IL5 | 7 | 10 | 1623 | 46.371.428.571.428.500 | 0.5415699913185681 | 0.1093773628538538 | 0.1093773628538538 |
| GOTERM_BP_DIRECT | GO:0042102~positive regulation of T cell proliferation | 2 | 15.384.615.384.615.300 | 0.036479161987166504 | IL4, IL23R | 13 | 61 | 19734 | 49.770.491.803.278.600 | 0.9999635409223181 | 0.3058393064034729 | 0.2835964477559476 |
| GOTERM_BP_DIRECT | GO:0045944~positive regulation of transcription by RNA polymerase II | 4 | 3.076.923.076.923.070 | 0.03670071676841675 | IL10, IL4, CXCL10, IGF2 | 13 | 1256 | 19734 | 4.834.394.904.458.590 | 0.9999657752556356 | 0.3058393064034729 | 0.2835964477559476 |
| REACTOME_PATHWAY | R-HSA-162582~Signal Transduction | 7 | 5.384.615.384.615.380 | 0.0406044321697458 | FKBP1A, IL3, CXCL10, IL5, CXCR5, IGF2, CCL1 | 13 | 2599 | 11091 | 22.978.364.459.703.400 | 0.9514898441923377 | 0.26457308096133686 | 0.2319544819387063 |
| REACTOME_PATHWAY | R-HSA-418594~G alpha (i) signalling events | 3 | 23.076.923.076.923.000 | 0.04474334338816362 | CXCL10, CXCR5, CCL1 | 13 | 318 | 11091 | 8.048.621.190.130.620 | 0.9646193906980208 | 0.26457308096133686 | 0.2319544819387063 |
| GOTERM_BP_DIRECT | GO:0032729~positive regulation of type II interferon production | 2 | 15.384.615.384.615.300 | 0.04642585064105849 | CD160, IL23R | 13 | 78 | 19734 | 3.892.307.692.307.690 | 0.9999978987437252 | 0.36934863617414576 | 0.3424869171796624 |
| REACTOME_PATHWAY | R-HSA-451927~Interleukin-2 family signaling | 2 | 15.384.615.384.615.300 | 0.046603652296059735 | IL3, IL5 | 13 | 44 | 11091 | 3.877.972.027.972.020 | 0.9693123011385119 | 0.26457308096133686 | 0.2319544819387063 |
| GOTERM_BP_DIRECT | GO:0032720~negative regulation of tumor necrosis factor production | 2 | 15.384.615.384.615.300 | 0.047008008240345824 | IL10, IL4 | 13 | 79 | 19734 | 3.843.037.974.683.540 | 0.9999982235864793 | 0.36934863617414576 | 0.3424869171796624 |
| BIOCARTA | h_asbcellPathway:Antigen Dependent B Cell Activation | 2 | 15.384.615.384.615.300 | 0.047178268279158615 | IL10, IL4 | 7 | 13 | 1623 | 3.567.032.967.032.960 | 0.6375528148818954 | 0.12384295423279136 | 0.12384295423279136 |
| REACTOME_PATHWAY | R-HSA-373076~Class A/1 (Rhodopsin-like receptors) | 3 | 23.076.923.076.923.000 | 0.0488955745063329 | CXCL10, CXCR5, CCL1 | 13 | 334 | 11091 | 766.305.849.838.784 | 0.9742570223809903 | 0.26457308096133686 | 0.2319544819387063 |
| GOTERM_BP_DIRECT | GO:0030593~neutrophil chemotaxis | 2 | 15.384.615.384.615.300 | 0.04933338223244672 | CXCL10, CCL1 | 13 | 83 | 19734 | 3.657.831.325.301.200 | 0.9999990926661083 | 0.3768522253867458 | 0.3494447908131642 |
| GOTERM_MF_DIRECT | GO:0005515~protein binding | 12 | 923.076.923.076.923 | 0.04950281420682114 | IL10, FKBP1A, IL4, IL22, IL3, CXCL10, IL5, CD160, IL23R, IL13, CXCR5, IGF2 | 13 | 12754 | 19304 | 13.971.363.433.492.900 | 0.9286418736085917 | 0.6435365846886748 | 0.6435365846886748 |
| REACTOME_PATHWAY | R-HSA-6783783~Interleukin-10 signaling | 2 | 15.384.615.384.615.300 | 0.049707490397801445 | IL10, CXCL10 | 13 | 47 | 11091 | 3.630.441.898.527.000 | 0.9758129254094707 | 0.26457308096133686 | 0.2319544819387063 |
| REACTOME_PATHWAY | R-HSA-512988~Interleukin-3, Interleukin-5 and GM-CSF signaling | 2 | 15.384.615.384.615.300 | 0.050740042924092 | IL3, IL5 | 13 | 48 | 11091 | 3.554.807.692.307.690 | 0.9776582713087077 | 0.26457308096133686 | 0.2319544819387063 |

### Suppl. Table 5: Off-target nomination and CASTseq sites

| Chr | Start | Stop | Name | Score | Strand | Reason | Pool | UNCOVERseqNom? | ChariteNom? | GUIDE-seqNom? | InSilicoNom? | CAST-seqNom? | variants_ref | score | variant_infl | effect_of_va | AF_AFR | AF_AMR | AF_EAS | AF_NFE | AF_SAS | **CASTseq re** | **CASTseq hi** |
|---|---|---|---|---|---|---|---|---|---|---|---|---|---|---|---|---|---|---|---|---|---|---|---|
| chr20 | 1392862 | 1392882 | OnT | 100 | - | EmpiricalOverlap | 1 | true | true | true | true | false | N/A | N/A | N/A | N/A | N/A | N/A | N/A | N/A | N/A | N/A | N/A |
| chr16 | 4114504 | 4114524 | Tier1_OT2_3_Exon | 12,054 | - | EmpiricalOverlap | 1 | true | true | true | false | false | N/A | N/A | N/A | N/A | N/A | N/A | N/A | N/A | N/A | N/A | N/A |
| chr17 | 19743182 | 19743202 | Tier1_OT3_3_Exon | 2,785 | - | EmpiricalOverlap | 1 | true | true | true | true | false | N/A | N/A | N/A | N/A | N/A | N/A | N/A | N/A | N/A | N/A | N/A |
| chr1 | 17232953 | 17232973 | Tier2_OT1_3_Exon | 410,421 | + | EmpiricalOverlap | 1 | true | true | true | true | false | N/A | N/A | N/A | N/A | N/A | N/A | N/A | N/A | N/A | N/A | N/A |
| chr16 | 4925244 | 4925264 | Tier2_OT2_3_Exon | 33,539 | - | EmpiricalOverlap | 1 | true | true | true | true | false | N/A | N/A | N/A | N/A | N/A | N/A | N/A | N/A | N/A | N/A | N/A |
| chr11 | 114399682 | 1,14E+08 | Tier2_OT3_3_Exon | 13,444 | - | EmpiricalOverlap | 1 | true | true | true | true | false | N/A | N/A | N/A | N/A | N/A | N/A | N/A | N/A | N/A | N/A | N/A |
| chr17 | 64780889 | 64780909 | Tier2_OT4_3_Exon | 7,386 | + | EmpiricalOverlap | 1 | true | true | true | false | false | N/A | N/A | N/A | N/A | N/A | N/A | N/A | N/A | N/A | N/A | N/A |
| chr7 | 5428495 | 5428515 | Tier2_OT5_3_Exon | 3,971 | - | EmpiricalOverlap | 1 | true | true | true | true | false | N/A | N/A | N/A | N/A | N/A | N/A | N/A | N/A | N/A | N/A | N/A |
| chr17 | 75262316 | 75262336 | Tier2_OT6_3_Exon | 1,672 | - | EmpiricalOverlap | 1 | true | true | true | false | false | N/A | N/A | N/A | N/A | N/A | N/A | N/A | N/A | N/A | N/A | N/A |
| chr15 | 26111371 | 26111391 | Tier2_OT7_3_Exon | 1,627 | + | EmpiricalOverlap | 1 | true | false | false | true | false | N/A | N/A | N/A | N/A | N/A | N/A | N/A | N/A | N/A | N/A | N/A |
| chr19 | 48393561 | 48393581 | Tier2_OT9_3_Exon | 0,906 | + | EmpiricalOverlap | 1 | true | false | false | true | false | N/A | N/A | N/A | N/A | N/A | N/A | N/A | N/A | N/A | N/A | N/A |
| chr12 | 29784506 | 29784526 | Tier2_OT10_3_Exon | 0,878 | + | EmpiricalOverlap | 1 | true | true | true | false | false | N/A | N/A | N/A | N/A | N/A | N/A | N/A | N/A | N/A | N/A | N/A |
| chr10 | 88574789 | 88574809 | Tier3_OT1_3_Intron | 215,583 | - | EmpiricalOverlap | 1 | true | true | true | true | false | N/A | N/A | N/A | N/A | N/A | N/A | N/A | N/A | N/A | N/A | N/A |
| chr18 | 76787668 | 76787688 | Tier3_OT2_3_Intergenic | 158,461 | - | EmpiricalOverlap | 1 | true | true | true | true | false | N/A | N/A | N/A | N/A | N/A | N/A | N/A | N/A | N/A | N/A | N/A |
| chrX | 21733091 | 21733111 | Tier3_OT3_3_Intron | 62,154 | + | EmpiricalOverlap | 1 | true | true | true | false | false | N/A | N/A | N/A | N/A | N/A | N/A | N/A | N/A | N/A | N/A | N/A |
| chr12 | 22800505 | 22800525 | Tier3_OT4_3_Intergenic | 53,25 | + | EmpiricalOverlap | 1 | true | true | true | false | false | N/A | N/A | N/A | N/A | N/A | N/A | N/A | N/A | N/A | N/A | N/A |
| chr8 | 1823549 | 1823569 | Tier3_OT5_3_Intergenic | 50,32 | + | EmpiricalOverlap | 1 | true | true | true | true | false | N/A | N/A | N/A | N/A | N/A | N/A | N/A | N/A | N/A | N/A | N/A |
| chr1 | 53517375 | 53517395 | Tier3_OT6_3_Intron | 30,554 | + | EmpiricalOverlap | 1 | true | true | true | false | false | N/A | N/A | N/A | N/A | N/A | N/A | N/A | N/A | N/A | N/A | N/A |
| chr6 | 99319061 | 99319081 | Tier3_OT7_3_Intron | 24,3 | + | EmpiricalOverlap | 1 | true | true | true | true | false | N/A | N/A | N/A | N/A | N/A | N/A | N/A | N/A | N/A | N/A | N/A |
| chr14 | 104972675 | 1,05E+08 | Tier3_OT8_3_Intron | 19,59 | + | EmpiricalOverlap | 1 | true | true | true | true | false | N/A | N/A | N/A | N/A | N/A | N/A | N/A | N/A | N/A | N/A | N/A |
| chr12 | 58804887 | 58804907 | Tier3_OT9_3_Other | 17,868 | - | EmpiricalOverlap | 1 | true | true | true | true | false | N/A | N/A | N/A | N/A | N/A | N/A | N/A | N/A | N/A | N/A | N/A |
| chr12 | 30200891 | 30200911 | Tier3_OT10_3_Intergenic | 17,867 | - | EmpiricalOverlap | 1 | true | true | true | false | false | N/A | N/A | N/A | N/A | N/A | N/A | N/A | N/A | N/A | N/A | N/A |
| chr11 | 2269204 | 2269224 | Tier3_OT11_3_Intron | 17,035 | + | EmpiricalOverlap | 1 | true | true | true | true | false | N/A | N/A | N/A | N/A | N/A | N/A | N/A | N/A | N/A | N/A | N/A |
| chrX | 140837128 | 1,41E+08 | Tier3_OT12_3_Intergenic | 16,984 | + | EmpiricalOverlap | 1 | true | true | true | true | false | N/A | N/A | N/A | N/A | N/A | N/A | N/A | N/A | N/A | N/A | N/A |
| chr13 | 29551855 | 29551875 | Tier3_OT13_3_Intron | 15,118 | - | EmpiricalOverlap | 1 | true | true | true | false | false | N/A | N/A | N/A | N/A | N/A | N/A | N/A | N/A | N/A | N/A | N/A |
| chr1 | 234922607 | 2,35E+08 | Tier3_OT14_3_Intergenic | 12,167 | + | EmpiricalOverlap | 1 | true | true | true | false | false | N/A | N/A | N/A | N/A | N/A | N/A | N/A | N/A | N/A | N/A | N/A |
| chr10 | 71551850 | 71551870 | Tier3_OT15_3_Intron | 11,215 | + | EmpiricalOverlap | 1 | true | true | true | false | false | N/A | N/A | N/A | N/A | N/A | N/A | N/A | N/A | N/A | N/A | N/A |
| chr7 | 68843471 | 68843491 | Tier3_OT16_3_Intergenic | 10,628 | + | EmpiricalOverlap | 1 | true | true | true | false | false | N/A | N/A | N/A | N/A | N/A | N/A | N/A | N/A | N/A | N/A | N/A |
| chr11 | 2378955 | 2378975 | Tier3_OT17_3_Intron | 8,365 | + | EmpiricalOverlap | 1 | true | true | true | true | false | N/A | N/A | N/A | N/A | N/A | N/A | N/A | N/A | N/A | N/A | N/A |
| chr21 | 31387820 | 31387840 | Tier3_OT18_3_Intron | 8 | + | EmpiricalOverlap | 1 | true | true | true | true | false | N/A | N/A | N/A | N/A | N/A | N/A | N/A | N/A | N/A | N/A | N/A |
| chr15 | 72169222 | 72169242 | Tier3_OT19_3_Intron | 7,073 | + | EmpiricalOverlap | 1 | true | true | true | false | false | N/A | N/A | N/A | N/A | N/A | N/A | N/A | N/A | N/A | N/A | N/A |
| chr20 | 44774034 | 44774054 | Tier3_OT20_3_Intron | 6,693 | + | EmpiricalOverlap | 1 | true | false | false | true | false | N/A | N/A | N/A | N/A | N/A | N/A | N/A | N/A | N/A | N/A | N/A |
| chr5 | 140561928 | 1,41E+08 | Tier3_OT21_3_Intron | 6,439 | + | EmpiricalOverlap | 1 | true | true | true | true | false | N/A | N/A | N/A | N/A | N/A | N/A | N/A | N/A | N/A | N/A | N/A |
| chr4 | 7423135 | 7423155 | Tier3_OT22_3_Intron | 6,4 | - | EmpiricalOverlap | 1 | true | true | false | true | false | N/A | N/A | N/A | N/A | N/A | N/A | N/A | N/A | N/A | N/A | N/A |
| chr15 | 31983496 | 31983516 | Tier3_OT23_3_Intergenic | 6,169 | - | EmpiricalOverlap | 1 | true | true | true | false | false | N/A | N/A | N/A | N/A | N/A | N/A | N/A | N/A | N/A | N/A | N/A |
| chr12 | 30170651 | 30170671 | Tier3_OT24_3_Intergenic | 5,333 | + | EmpiricalOverlap | 1 | true | true | true | false | false | N/A | N/A | N/A | N/A | N/A | N/A | N/A | N/A | N/A | N/A | N/A |
| chr14 | 99320885 | 99320905 | Tier3_OT25_3_Intergenic | 5,053 | + | EmpiricalOverlap | 1 | true | true | true | false | false | N/A | N/A | N/A | N/A | N/A | N/A | N/A | N/A | N/A | N/A | N/A |
| chr6 | 81017763 | 81017782 | Tier3_OT27_3_Intergenic | 4,682 | + | EmpiricalOverlap | 1 | true | true | false | true | false | A>T | 0,924311 | 0,89071 | Reduced | 0,078 | 0,00642 | 0 | 0,00057 | 0,00104 | N/A | N/A |
| chr3 | 27630389 | 27630408 | Tier3_OT28_3_Intergenic | 4,081 | + | EmpiricalOverlap | 3 | true | false | false | true | false | N/A | N/A | N/A | N/A | N/A | N/A | N/A | N/A | N/A | N/A | N/A |
| chr1 | 19645120 | 19645140 | Tier3_OT29_3_Intron | 3,874 | + | EmpiricalOverlap | 1 | true | false | false | true | false | N/A | N/A | N/A | N/A | N/A | N/A | N/A | N/A | N/A | N/A | N/A |
| chr10 | 49768942 | 49768962 | Tier3_OT31_3_Intergenic | 3,299 | - | EmpiricalOverlap | 1 | true | true | false | true | false | N/A | N/A | N/A | N/A | N/A | N/A | N/A | N/A | N/A | N/A | N/A |
| chr19 | 6048056 | 6048076 | Tier3_OT34_3_Intron | 3,099 | + | EmpiricalOverlap | 1 | true | true | true | false | false | N/A | N/A | N/A | N/A | N/A | N/A | N/A | N/A | N/A | N/A | N/A |
| chr1 | 41476583 | 41476603 | Tier3_OT36_3_Intergenic | 2,912 | + | EmpiricalOverlap | 1 | true | true | true | false | false | N/A | N/A | N/A | N/A | N/A | N/A | N/A | N/A | N/A | N/A | N/A |
| chr3 | 184604572 | 1,85E+08 | Tier3_OT37_3_Intergenic | 2,889 | - | EmpiricalOverlap | 1 | true | true | true | true | false | N/A | N/A | N/A | N/A | N/A | N/A | N/A | N/A | N/A | N/A | N/A |
| chr1 | 13805459 | 13805479 | Tier3_OT38_3_Intron | 2,847 | - | EmpiricalOverlap | 1 | true | true | true | true | false | N/A | N/A | N/A | N/A | N/A | N/A | N/A | N/A | N/A | N/A | N/A |
| chr13 | 30444488 | 30444508 | Tier3_OT39_3_Intergenic | 2,649 | - | EmpiricalOverlap | 1 | true | true | true | false | false | N/A | N/A | N/A | N/A | N/A | N/A | N/A | N/A | N/A | N/A | N/A |
| chr13 | 50409011 | 50409031 | Tier3_OT40_3_Other | 2,566 | - | EmpiricalOverlap | 1 | true | true | true | false | false | N/A | N/A | N/A | N/A | N/A | N/A | N/A | N/A | N/A | N/A | N/A |
| chr17 | 9700314 | 9700334 | Tier3_OT45_3_Intron | 1,947 | + | EmpiricalOverlap | 1 | true | false | false | true | false | N/A | N/A | N/A | N/A | N/A | N/A | N/A | N/A | N/A | N/A | N/A |
| chr15 | 72288652 | 72288671 | Tier3_OT46_3_Intron | 1,749 | + | EmpiricalOverlap | 1 | true | false | false | true | false | N/A | N/A | N/A | N/A | N/A | N/A | N/A | N/A | N/A | N/A | N/A |
| chr16 | 66866410 | 66866430 | Tier3_OT47_3_Intergenic | 1,74 | - | EmpiricalOverlap | 1 | true | true | true | true | false | N/A | N/A | N/A | N/A | N/A | N/A | N/A | N/A | N/A | N/A | N/A |
| chr19 | 14394829 | 14394849 | Tier3_OT49_3_Intron | 1,655 | - | EmpiricalOverlap | 1 | true | false | false | true | false | N/A | N/A | N/A | N/A | N/A | N/A | N/A | N/A | N/A | N/A | N/A |
| chr16 | 54896144 | 54896164 | Tier3_OT51_2_Intergenic | 1,61 | - | EmpiricalOverlap | 2 | true | false | false | true | false | N/A | N/A | N/A | N/A | N/A | N/A | N/A | N/A | N/A | N/A | N/A |
| chr20 | 57029869 | 57029889 | Tier3_OT53_3_Intergenic | 1,513 | + | EmpiricalOverlap | 1 | true | false | false | true | false | N/A | N/A | N/A | N/A | N/A | N/A | N/A | N/A | N/A | N/A | N/A |
| chr1 | 65955715 | 65955735 | Tier3_OT54_3_Intron | 1,482 | - | EmpiricalOverlap | 1 | true | true | true | false | false | N/A | N/A | N/A | N/A | N/A | N/A | N/A | N/A | N/A | N/A | N/A |
| chrX | 105797334 | 1,06E+08 | Tier3_OT60_3_Intergenic | 1,193 | - | EmpiricalOverlap | 3 | true | true | false | true | false | N/A | N/A | N/A | N/A | N/A | N/A | N/A | N/A | N/A | N/A | N/A |
| chr18 | 1521794 | 1521813 | Tier3_OT61_3_Other | 1,185 | + | EmpiricalOverlap | 3 | true | true | false | true | false | N/A | N/A | N/A | N/A | N/A | N/A | N/A | N/A | N/A | N/A | N/A |
| chr19 | 49645652 | 49645672 | Tier3_OT63_3_Intron | 1,125 | + | EmpiricalOverlap | 1 | true | false | false | true | false | N/A | N/A | N/A | N/A | N/A | N/A | N/A | N/A | N/A | N/A | N/A |
| chr3 | 127224479 | 1,27E+08 | Tier3_OT68_3_Intergenic | 0,992 | + | EmpiricalOverlap | 1 | true | true | true | false | false | N/A | N/A | N/A | N/A | N/A | N/A | N/A | N/A | N/A | N/A | N/A |
| chr3 | 48639290 | 48639309 | Tier3_OT69_3_Intron | 0,984 | + | EmpiricalOverlap | 1 | true | true | true | true | false | N/A | N/A | N/A | N/A | N/A | N/A | N/A | N/A | N/A | N/A | N/A |
| chr15 | 40916634 | 40916654 | Tier3_OT70_3_Intergenic | 0,964 | + | EmpiricalOverlap | 1 | true | false | false | true | false | N/A | N/A | N/A | N/A | N/A | N/A | N/A | N/A | N/A | N/A | N/A |
| chr6 | 2948030 | 2948050 | Tier3_OT71_3_Intergenic | 0,954 | + | EmpiricalOverlap | 1 | true | true | false | true | false | N/A | N/A | N/A | N/A | N/A | N/A | N/A | N/A | N/A | N/A | N/A |
| chr10 | 77537419 | 77537439 | Tier3_OT75_3_Intron | 0,757 | + | EmpiricalOverlap | 1 | true | true | true | true | false | N/A | N/A | N/A | N/A | N/A | N/A | N/A | N/A | N/A | N/A | N/A |
| chr14 | 73262830 | 73262850 | Tier3_OT77_3_Intron | 0,709 | + | EmpiricalOverlap | 1 | true | false | false | true | false | N/A | N/A | N/A | N/A | N/A | N/A | N/A | N/A | N/A | N/A | N/A |
| chr12 | 44248246 | 44248266 | Tier3_OT78_2_Intron | 0,707 | - | EmpiricalOverlap | 1 | true | false | false | true | false | N/A | N/A | N/A | N/A | N/A | N/A | N/A | N/A | N/A | N/A | N/A |
| chr9 | 126228187 | 1,26E+08 | Tier3_OT79_2_Other | 0,676 | + | EmpiricalOverlap | 1 | true | true | false | true | false | N/A | N/A | N/A | N/A | N/A | N/A | N/A | N/A | N/A | N/A | N/A |
| chr3 | 178018347 | 1,78E+08 | Tier3_OT80_3_Other | 0,641 | - | EmpiricalOverlap | 1 | true | false | false | true | false | N/A | N/A | N/A | N/A | N/A | N/A | N/A | N/A | N/A | N/A | N/A |
| chr19 | 38702809 | 38702829 | Tier3_OT82_2_Intron | 0,588 | - | EmpiricalOverlap | 1 | true | false | false | true | false | N/A | N/A | N/A | N/A | N/A | N/A | N/A | N/A | N/A | N/A | N/A |
| chr16 | 81403917 | 81403937 | Tier3_OT84_2_Intergenic | 0,557 | - | EmpiricalOverlap | 1 | true | true | true | true | false | N/A | N/A | N/A | N/A | N/A | N/A | N/A | N/A | N/A | N/A | N/A |
| chr10 | 54288800 | 54288820 | Tier3_OT87_2_Intron | 0,528 | - | EmpiricalOverlap | 1 | true | false | false | true | false | N/A | N/A | N/A | N/A | N/A | N/A | N/A | N/A | N/A | N/A | N/A |
| chr9 | 136288654 | 1,36E+08 | Tier3_OT89_2_Intron | 0,519 | + | EmpiricalOverlap | 1 | true | false | false | true | false | N/A | N/A | N/A | N/A | N/A | N/A | N/A | N/A | N/A | N/A | N/A |
| chr1 | 109748498 | 1,1E+08 | Tier3_OT90_3_Intergenic | 0,518 | + | EmpiricalOverlap | 1 | true | false | false | true | false | N/A | N/A | N/A | N/A | N/A | N/A | N/A | N/A | N/A | N/A | N/A |
| chr13 | 60832898 | 60832917 | Tier3_OT96_3_Other | 0,417 | + | EmpiricalOverlap | 1 | true | true | false | true | false | N/A | N/A | N/A | N/A | N/A | N/A | N/A | N/A | N/A | N/A | N/A |
| chr14 | 102509166 | 1,03E+08 | Tier3_OT100_1_Exon | 0,383 | + | EmpiricalOverlap | 1 | true | false | false | true | false | N/A | N/A | N/A | N/A | N/A | N/A | N/A | N/A | N/A | N/A | N/A |
| chr2 | 126831888 | 1,27E+08 | Tier3_OT104_2_Intergenic | 0,355 | + | EmpiricalOverlap | 1 | true | false | false | true | false | N/A | N/A | N/A | N/A | N/A | N/A | N/A | N/A | N/A | N/A | N/A |
| chr12 | 117676233 | 1,18E+08 | Tier3_OT105_2_Intron | 0,351 | + | EmpiricalOverlap | 1 | true | false | false | true | false | N/A | N/A | N/A | N/A | N/A | N/A | N/A | N/A | N/A | N/A | N/A |
| chr15 | 52253098 | 52253117 | Tier3_OT106_2_Intron | 0,338 | + | EmpiricalOverlap | 1 | true | false | false | true | false | N/A | N/A | N/A | N/A | N/A | N/A | N/A | N/A | N/A | N/A | N/A |
| chr22 | 43489428 | 43489448 | Tier3_OT108_1_Intron | 0,33 | - | EmpiricalOverlap | 1 | true | false | false | true | false | N/A | N/A | N/A | N/A | N/A | N/A | N/A | N/A | N/A | N/A | N/A |
| chr11 | 128690983 | 1,29E+08 | Tier3_OT109_2_Intron | 0,315 | + | EmpiricalOverlap | 1 | true | false | false | true | false | N/A | N/A | N/A | N/A | N/A | N/A | N/A | N/A | N/A | N/A | N/A |
| chr17 | 13943481 | 13943501 | Tier3_OT110_2_Intergenic | 0,305 | - | EmpiricalOverlap | 1 | true | false | false | true | false | N/A | N/A | N/A | N/A | N/A | N/A | N/A | N/A | N/A | N/A | N/A |
| chr5 | 176597699 | 1,77E+08 | Tier3_OT120_1_Exon | 0,247 | + | EmpiricalOverlap | 1 | true | false | false | true | false | N/A | N/A | N/A | N/A | N/A | N/A | N/A | N/A | N/A | N/A | N/A |
| chrX | 17667850 | 17667870 | Tier3_OT130_1_Intron | 0,19 | + | EmpiricalOverlap | 1 | true | false | false | true | false | N/A | N/A | N/A | N/A | N/A | N/A | N/A | N/A | N/A | N/A | N/A |
| chr16 | 68263917 | 68263937 | Tier3_OT137_2_Intergenic | 0,153 | + | EmpiricalOverlap | 1 | true | false | false | true | false | N/A | N/A | N/A | N/A | N/A | N/A | N/A | N/A | N/A | N/A | N/A |
| chr1 | 10752073 | 10752093 | Tier3_OT146_1_Intron | 0,095 | + | EmpiricalOverlap | 1 | true | true | true | true | false | N/A | N/A | N/A | N/A | N/A | N/A | N/A | N/A | N/A | N/A | N/A |
| chr21 | 38423357 | 38423377 | Tier3_OT150_1_Intron | 0,079 | - | EmpiricalOverlap | 1 | true | false | false | true | false | N/A | N/A | N/A | N/A | N/A | N/A | N/A | N/A | N/A | N/A | N/A |
| chrX | 154996509 | 1,55E+08 | Tier3_OT152_1_Intron | 0,077 | - | EmpiricalOverlap | 1 | true | false | false | true | false | N/A | N/A | N/A | N/A | N/A | N/A | N/A | N/A | N/A | N/A | N/A |
| chr1 | 92486864 | 92486884 | Tier3_OT160_1_Exon | 0,045 | + | EmpiricalOverlap | 1 | true | false | false | true | false | N/A | N/A | N/A | N/A | N/A | N/A | N/A | N/A | N/A | N/A | N/A |
| chr2 | 70767629 | 70767649 | Tier3_OT169_1_Exon | 0,014 | + | EmpiricalOverlap | 1 | true | false | false | true | false | N/A | N/A | N/A | N/A | N/A | N/A | N/A | N/A | N/A | N/A | N/A |
| chr2 | 51743495 | 51743515 | Tier5_OT3_1_Other | 0,377 | - | EmpiricalOverlap | 1 | true | false | false | true | false | N/A | N/A | N/A | N/A | N/A | N/A | N/A | N/A | N/A | N/A | N/A |
| chr7 | 76149915 | 76149935 | Tier5_OT14_1_Intergenic | 0,16 | + | EmpiricalOverlap | 1 | true | false | false | true | false | N/A | N/A | N/A | N/A | N/A | N/A | N/A | N/A | N/A | N/A | N/A |
| chr17 | 16836693 | 16836713 | Tier5_OT19_1_Intergenic | 0,131 | + | EmpiricalOverlap | 1 | true | false | false | true | false | N/A | N/A | N/A | N/A | N/A | N/A | N/A | N/A | N/A | N/A | N/A |
| chr1 | 69776102 | 69776122 | Tier5_OT26_1_Intron | 0,12 | - | EmpiricalOverlap | 1 | true | false | false | true | false | N/A | N/A | N/A | N/A | N/A | N/A | N/A | N/A | N/A | N/A | N/A |
| chr10 | 122573653 | 1,23E+08 | Tier5_OT29_1_Intron | 0,115 | + | EmpiricalOverlap | 1 | true | false | false | true | false | N/A | N/A | N/A | N/A | N/A | N/A | N/A | N/A | N/A | N/A | N/A |
| chr1 | 86205835 | 86205852 | Tier5_OT30_1_Intergenic | 0,111 | + | EmpiricalOverlap | 1 | true | false | false | true | false | N/A | N/A | N/A | N/A | N/A | N/A | N/A | N/A | N/A | N/A | N/A |
| chr11 | 41728999 | 41729019 | Tier5_OT32_1_Other | 0,095 | + | EmpiricalOverlap | 1 | true | true | false | true | false | N/A | N/A | N/A | N/A | N/A | N/A | N/A | N/A | N/A | N/A | N/A |
| chr1 | 1243610 | 1243630 | Tier5_OT36_1_Intron | 0,063 | - | EmpiricalOverlap | 1 | true | false | false | true | false | N/A | N/A | N/A | N/A | N/A | N/A | N/A | N/A | N/A | N/A | N/A |
| chr11 | 22652501 | 22652521 | Tier5_OT37_1_Other | 0,031 | + | EmpiricalOverlap | 1 | true | true | false | true | false | N/A | N/A | N/A | N/A | N/A | N/A | N/A | N/A | N/A | N/A | N/A |
| chr19 | 46260916 | 46260936 | Tier5_OT38_1_Intergenic | 0,024 | - | EmpiricalOverlap | 1 | true | false | false | true | false | N/A | N/A | N/A | N/A | N/A | N/A | N/A | N/A | N/A | N/A | N/A |
| chr1 | 62308232 | 62308252 | Tier5_OT39_1_Intron | 0,018 | + | EmpiricalOverlap | 1 | true | false | false | true | false | N/A | N/A | N/A | N/A | N/A | N/A | N/A | N/A | N/A | N/A | N/A |
| chr7 | 149422995 | 1,49E+08 | Tier6_OT35_1_Intergenic | 0,023 | - | EmpiricalOverlap | 1 | true | false | false | true | false | N/A | N/A | N/A | N/A | N/A | N/A | N/A | N/A | N/A | N/A | N/A |

| | | | | | | | | | | | | | | | | | | | | | | | |
|---|---|---|---|---|---|---|---|---|---|---|---|---|---|---|---|---|---|---|---|---|---|---|---|
| chr11 | 20364011 | 20364031 | Tier2_OT8_3_Exon | 1,561 | + | UNCOVERseqTier3 | 1 | true | false | false | false | false | N/A | N/A | N/A | N/A | N/A | N/A | N/A | N/A | N/A | N/A | N/A |
| chr21 | 8434220 | 8434239 | Tier2_OT11_2_Exon | 0,387 | - | UNCOVERseqTier3 | 3 | true | false | false | false | false | N/A | N/A | N/A | N/A | N/A | N/A | N/A | N/A | N/A | N/A | N/A |
| chr16 | 84331972 | 84331992 | Tier3_OT26_3_Intergenic | 4,695 | + | UNCOVERseqTier3 | 1 | true | false | false | false | false | N/A | N/A | N/A | N/A | N/A | N/A | N/A | N/A | N/A | N/A | N/A |
| chr9 | 131079068 | 1,31E+08 | Tier3_OT30_3_Intron | 3,52 | + | UNCOVERseqTier3 | 1 | true | false | false | false | false | N/A | N/A | N/A | N/A | N/A | N/A | N/A | N/A | N/A | N/A | N/A |
| chr5 | 236357 | 236377 | Tier3_OT32_3_Intron | 3,25 | - | UNCOVERseqTier3 | 1 | true | false | false | false | false | N/A | N/A | N/A | N/A | N/A | N/A | N/A | N/A | N/A | N/A | N/A |
| chr22 | 45198602 | 45198622 | Tier3_OT33_3_Intron | 3,165 | - | UNCOVERseqTier3 | 1 | true | false | false | false | false | N/A | N/A | N/A | N/A | N/A | N/A | N/A | N/A | N/A | N/A | N/A |
| chr8 | 65733265 | 65733285 | Tier3_OT35_3_Intron | 3,093 | - | UNCOVERseqTier3 | 1 | true | false | false | false | false | N/A | N/A | N/A | N/A | N/A | N/A | N/A | N/A | N/A | N/A | N/A |
| chr1 | 154497113 | 1,54E+08 | Tier3_OT41_3_Intron | 2,432 | + | UNCOVERseqTier3 | 1 | true | false | false | false | false | N/A | N/A | N/A | N/A | N/A | N/A | N/A | N/A | N/A | N/A | N/A |
| chr2 | 86459600 | 86459620 | Tier3_OT42_3_Intron | 2,385 | - | UNCOVERseqTier3 | 1 | true | false | false | false | false | N/A | N/A | N/A | N/A | N/A | N/A | N/A | N/A | N/A | N/A | N/A |
| chr4 | 185905181 | 1,86E+08 | Tier3_OT43_3_Intron | 2,254 | + | UNCOVERseqTier3 | 1 | true | false | false | false | false | N/A | N/A | N/A | N/A | N/A | N/A | N/A | N/A | N/A | N/A | N/A |
| chr6 | 127121244 | 1,27E+08 | Tier3_OT44_3_Intron | 1,956 | + | UNCOVERseqTier3 | 1 | true | false | false | false | false | N/A | N/A | N/A | N/A | N/A | N/A | N/A | N/A | N/A | N/A | N/A |
| chr1 | 115347314 | 1,15E+08 | Tier3_OT48_3_Other | 1,737 | + | UNCOVERseqTier3 | 1 | true | false | false | false | false | N/A | N/A | N/A | N/A | N/A | N/A | N/A | N/A | N/A | N/A | N/A |
| chr6 | 6724661 | 6724681 | Tier3_OT50_3_Other | 1,61 | + | UNCOVERseqTier3 | 1 | true | false | false | false | false | N/A | N/A | N/A | N/A | N/A | N/A | N/A | N/A | N/A | N/A | N/A |
| chr9 | 136425638 | 1,36E+08 | Tier3_OT52_3_Intergenic | 1,593 | + | UNCOVERseqTier3 | 1 | true | false | false | false | false | N/A | N/A | N/A | N/A | N/A | N/A | N/A | N/A | N/A | N/A | N/A |
| chr6 | 39830620 | 39830640 | Tier3_OT55_3_Intron | 1,428 | + | UNCOVERseqTier3 | 1 | true | false | false | false | false | N/A | N/A | N/A | N/A | N/A | N/A | N/A | N/A | N/A | N/A | N/A |
| chr9 | 124217502 | 1,24E+08 | Tier3_OT56_3_Intergenic | 1,41 | + | UNCOVERseqTier3 | 1 | true | false | false | false | false | N/A | N/A | N/A | N/A | N/A | N/A | N/A | N/A | N/A | N/A | N/A |
| chr1 | 99849937 | 99849957 | Tier3_OT57_3_Intergenic | 1,372 | + | UNCOVERseqTier3 | 1 | true | false | false | false | false | N/A | N/A | N/A | N/A | N/A | N/A | N/A | N/A | N/A | N/A | N/A |
| chr3 | 135257793 | 1,35E+08 | Tier3_OT58_3_Intron | 1,364 | + | UNCOVERseqTier3 | 2 | true | false | false | false | false | N/A | N/A | N/A | N/A | N/A | N/A | N/A | N/A | N/A | N/A | N/A |
| chr2 | 11483045 | 11483065 | Tier3_OT59_3_Other | 1,301 | - | UNCOVERseqTier3 | 1 | true | false | false | false | false | N/A | N/A | N/A | N/A | N/A | N/A | N/A | N/A | N/A | N/A | N/A |
| chr16 | 21048256 | 21048276 | Tier3_OT62_3_Intron | 1,131 | + | UNCOVERseqTier3 | 3 | true | false | false | false | false | N/A | N/A | N/A | N/A | N/A | N/A | N/A | N/A | N/A | N/A | N/A |
| chr4 | 6373174 | 6373194 | Tier3_OT64_3_Intron | 1,074 | + | UNCOVERseqTier3 | 1 | true | false | false | false | false | N/A | N/A | N/A | N/A | N/A | N/A | N/A | N/A | N/A | N/A | N/A |
| chr17 | 62357358 | 62357378 | Tier3_OT65_3_Intergenic | 1,04 | - | UNCOVERseqTier3 | 1 | true | false | false | false | false | N/A | N/A | N/A | N/A | N/A | N/A | N/A | N/A | N/A | N/A | N/A |
| chr19 | 1017580 | 1017600 | Tier3_OT66_3_Intron | 1,02 | + | UNCOVERseqTier3 | 1 | true | false | false | false | false | N/A | N/A | N/A | N/A | N/A | N/A | N/A | N/A | N/A | N/A | N/A |
| chr22 | 27373462 | 27373482 | Tier3_OT67_3_Intergenic | 1,009 | - | UNCOVERseqTier3 | 1 | true | false | false | false | false | N/A | N/A | N/A | N/A | N/A | N/A | N/A | N/A | N/A | N/A | N/A |
| chr19 | 33430791 | 33430811 | Tier3_OT72_3_Intron | 0,949 | + | UNCOVERseqTier3 | 1 | true | false | false | false | false | N/A | N/A | N/A | N/A | N/A | N/A | N/A | N/A | N/A | N/A | N/A |
| chr1 | 30459282 | 30459302 | Tier3_OT73_3_Intergenic | 0,907 | + | UNCOVERseqTier3 | 1 | true | false | false | false | false | N/A | N/A | N/A | N/A | N/A | N/A | N/A | N/A | N/A | N/A | N/A |
| chr17 | 2883429 | 2883449 | Tier3_OT74_3_Intron | 0,887 | + | UNCOVERseqTier3 | 1 | true | false | false | false | false | N/A | N/A | N/A | N/A | N/A | N/A | N/A | N/A | N/A | N/A | N/A |
| chr1 | 23692185 | 23692205 | Tier3_OT76_3_Intron | 0,736 | - | UNCOVERseqTier3 | 1 | true | false | false | false | false | N/A | N/A | N/A | N/A | N/A | N/A | N/A | N/A | N/A | N/A | N/A |
| chr2 | 77604202 | 77604221 | Tier3_OT81_2_Intergenic | 0,634 | + | UNCOVERseqTier3 | 2 | true | false | false | false | false | N/A | N/A | N/A | N/A | N/A | N/A | N/A | N/A | N/A | N/A | N/A |
| chr1 | 119021045 | 1,19E+08 | Tier3_OT83_2_Intergenic | 0,578 | - | UNCOVERseqTier3 | 2 | true | false | false | false | false | N/A | N/A | N/A | N/A | N/A | N/A | N/A | N/A | N/A | N/A | N/A |
| chr14 | 35459157 | 35459177 | Tier3_OT85_2_Intergenic | 0,541 | - | UNCOVERseqTier3 | 1 | true | false | false | false | false | N/A | N/A | N/A | N/A | N/A | N/A | N/A | N/A | N/A | N/A | N/A |
| chr11 | 42940758 | 42940778 | Tier3_OT86_2_Intergenic | 0,532 | - | UNCOVERseqTier3 | 1 | true | false | false | false | false | N/A | N/A | N/A | N/A | N/A | N/A | N/A | N/A | N/A | N/A | N/A |
| chr19 | 52170914 | 52170934 | Tier3_OT88_2_Intron | 0,52 | + | UNCOVERseqTier3 | 1 | true | false | false | false | false | N/A | N/A | N/A | N/A | N/A | N/A | N/A | N/A | N/A | N/A | N/A |
| chr2 | 149550777 | 1,5E+08 | Tier3_OT91_3_Intergenic | 0,511 | + | UNCOVERseqTier3 | 3 | true | false | false | false | false | N/A | N/A | N/A | N/A | N/A | N/A | N/A | N/A | N/A | N/A | N/A |
| chr14 | 47795052 | 47795072 | Tier3_OT92_3_Intergenic | 0,501 | - | UNCOVERseqTier3 | 3 | true | false | false | false | false | N/A | N/A | N/A | N/A | N/A | N/A | N/A | N/A | N/A | N/A | N/A |
| chr22 | 48726790 | 48726810 | Tier3_OT93_1_Intron | 0,5 | + | UNCOVERseqTier3 | 1 | true | false | false | false | false | N/A | N/A | N/A | N/A | N/A | N/A | N/A | N/A | N/A | N/A | N/A |
| chr1 | 43547835 | 43547855 | Tier3_OT94_2_Intron | 0,483 | + | UNCOVERseqTier3 | 1 | true | false | false | false | false | N/A | N/A | N/A | N/A | N/A | N/A | N/A | N/A | N/A | N/A | N/A |
| chr8 | 118510585 | 1,19E+08 | Tier3_OT95_1_Intron | 0,476 | + | UNCOVERseqTier3 | 1 | true | false | false | false | false | N/A | N/A | N/A | N/A | N/A | N/A | N/A | N/A | N/A | N/A | N/A |
| chr16 | 1183951 | 1183971 | Tier3_OT97_1_Intron | 0,413 | - | UNCOVERseqTier3 | 1 | true | false | false | false | false | N/A | N/A | N/A | N/A | N/A | N/A | N/A | N/A | N/A | N/A | N/A |
| chr18 | 27510435 | 27510455 | Tier3_OT98_2_Other | 0,39 | - | UNCOVERseqTier3 | 3 | true | false | false | false | false | N/A | N/A | N/A | N/A | N/A | N/A | N/A | N/A | N/A | N/A | N/A |
| chr6 | 14480399 | 14480419 | Tier3_OT99_2_Other | 0,386 | - | UNCOVERseqTier3 | 3 | true | false | false | false | false | N/A | N/A | N/A | N/A | N/A | N/A | N/A | N/A | N/A | N/A | N/A |
| chr9 | 98176417 | 98176437 | Tier3_OT101_1_Intron | 0,381 | - | UNCOVERseqTier3 | 1 | true | false | false | false | false | N/A | N/A | N/A | N/A | N/A | N/A | N/A | N/A | N/A | N/A | N/A |
| chr19 | 3288014 | 3288034 | Tier3_OT102_2_Intron | 0,375 | - | UNCOVERseqTier3 | 1 | true | false | false | false | false | N/A | N/A | N/A | N/A | N/A | N/A | N/A | N/A | N/A | N/A | N/A |
| chr5 | 64714620 | 64714640 | Tier3_OT103_3_Intron | 0,364 | + | UNCOVERseqTier3 | 3 | true | false | false | false | false | N/A | N/A | N/A | N/A | N/A | N/A | N/A | N/A | N/A | N/A | N/A |
| chrX | 98300767 | 98300787 | Tier3_OT107_2_Intergenic | 0,33 | - | UNCOVERseqTier3 | 1 | true | false | false | false | false | N/A | N/A | N/A | N/A | N/A | N/A | N/A | N/A | N/A | N/A | N/A |
| chr8 | 102317375 | 1,02E+08 | Tier3_OT111_2_Intron | 0,293 | - | UNCOVERseqTier3 | 3 | true | false | false | false | false | N/A | N/A | N/A | N/A | N/A | N/A | N/A | N/A | N/A | N/A | N/A |
| chr2 | 148154512 | 1,48E+08 | Tier3_OT112_2_Intron | 0,289 | + | UNCOVERseqTier3 | 1 | true | false | false | false | false | N/A | N/A | N/A | N/A | N/A | N/A | N/A | N/A | N/A | N/A | N/A |
| chr7 | 100867695 | 1,01E+08 | Tier3_OT113_3_Intron | 0,287 | + | UNCOVERseqTier3 | 1 | true | false | false | false | false | N/A | N/A | N/A | N/A | N/A | N/A | N/A | N/A | N/A | N/A | N/A |
| chr10 | 24094939 | 24094959 | Tier3_OT114_1_Intron | 0,282 | + | UNCOVERseqTier3 | 3 | true | false | false | false | false | N/A | N/A | N/A | N/A | N/A | N/A | N/A | N/A | N/A | N/A | N/A |
| chr6 | 36754783 | 36754803 | Tier3_OT115_1_Intron | 0,278 | - | UNCOVERseqTier3 | 1 | true | false | false | false | false | N/A | N/A | N/A | N/A | N/A | N/A | N/A | N/A | N/A | N/A | N/A |
| chr22 | 49433267 | 49433287 | Tier3_OT116_2_Other | 0,273 | - | UNCOVERseqTier3 | 1 | true | false | false | false | false | N/A | N/A | N/A | N/A | N/A | N/A | N/A | N/A | N/A | N/A | N/A |
| chrX | 107156192 | 1,07E+08 | Tier3_OT117_2_Intron | 0,272 | - | UNCOVERseqTier3 | 3 | true | false | false | false | false | N/A | N/A | N/A | N/A | N/A | N/A | N/A | N/A | N/A | N/A | N/A |
| chrX | 104950551 | 1,05E+08 | Tier3_OT118_1_Intron | 0,271 | + | UNCOVERseqTier3 | 1 | true | false | false | false | false | N/A | N/A | N/A | N/A | N/A | N/A | N/A | N/A | N/A | N/A | N/A |
| chr10 | 18627820 | 18627840 | Tier3_OT119_2_Intron | 0,261 | - | UNCOVERseqTier3 | 1 | true | false | false | false | false | N/A | N/A | N/A | N/A | N/A | N/A | N/A | N/A | N/A | N/A | N/A |
| chr17 | 17042051 | 17042071 | Tier3_OT121_2_Intergenic | 0,243 | - | UNCOVERseqTier3 | 1 | true | false | false | false | false | N/A | N/A | N/A | N/A | N/A | N/A | N/A | N/A | N/A | N/A | N/A |
| chr15 | 90221537 | 90221557 | Tier3_OT122_2_Intron | 0,227 | + | UNCOVERseqTier3 | 1 | true | false | false | false | false | N/A | N/A | N/A | N/A | N/A | N/A | N/A | N/A | N/A | N/A | N/A |
| chr4 | 97904012 | 97904032 | Tier3_OT123_2_Intron | 0,227 | - | UNCOVERseqTier3 | 2 | true | false | false | false | false | N/A | N/A | N/A | N/A | N/A | N/A | N/A | N/A | N/A | N/A | N/A |
| chr17 | 1159985 | 1160004 | Tier3_OT124_1_Intron | 0,225 | - | UNCOVERseqTier3 | 1 | true | false | false | false | false | N/A | N/A | N/A | N/A | N/A | N/A | N/A | N/A | N/A | N/A | N/A |
| chr17 | 81666856 | 81666876 | Tier3_OT125_1_Exon | 0,219 | - | UNCOVERseqTier3 | 1 | true | false | false | false | false | N/A | N/A | N/A | N/A | N/A | N/A | N/A | N/A | N/A | N/A | N/A |
| chr19 | 46227548 | 46227568 | Tier3_OT126_2_Intergenic | 0,209 | - | UNCOVERseqTier3 | 1 | true | false | false | false | false | N/A | N/A | N/A | N/A | N/A | N/A | N/A | N/A | N/A | N/A | N/A |
| chr9 | 136845404 | 1,37E+08 | Tier3_OT127_1_Exon | 0,2 | + | UNCOVERseqTier3 | 1 | true | false | false | false | false | N/A | N/A | N/A | N/A | N/A | N/A | N/A | N/A | N/A | N/A | N/A |
| chr8 | 97595282 | 97595302 | Tier3_OT128_2_Intergenic | 0,196 | + | UNCOVERseqTier3 | 3 | true | false | false | false | false | N/A | N/A | N/A | N/A | N/A | N/A | N/A | N/A | N/A | N/A | N/A |
| chr7 | 15940654 | 15940674 | Tier3_OT129_2_Intergenic | 0,191 | - | UNCOVERseqTier3 | 1 | true | false | false | false | false | N/A | N/A | N/A | N/A | N/A | N/A | N/A | N/A | N/A | N/A | N/A |
| chr11 | 28238311 | 28238331 | Tier3_OT131_2_Intron | 0,185 | + | UNCOVERseqTier3 | 2 | true | false | false | false | false | N/A | N/A | N/A | N/A | N/A | N/A | N/A | N/A | N/A | N/A | N/A |
| chr19 | 42310374 | 42310394 | Tier3_OT132_1_Exon | 0,181 | + | UNCOVERseqTier3 | 1 | true | false | false | false | false | N/A | N/A | N/A | N/A | N/A | N/A | N/A | N/A | N/A | N/A | N/A |
| chr5 | 181217934 | 1,81E+08 | Tier3_OT133_1_Exon | 0,174 | + | UNCOVERseqTier3 | 1 | true | false | false | false | false | N/A | N/A | N/A | N/A | N/A | N/A | N/A | N/A | N/A | N/A | N/A |
| chr1 | 44740222 | 44740242 | Tier3_OT134_2_Intron | 0,166 | + | UNCOVERseqTier3 | 1 | true | false | false | false | false | N/A | N/A | N/A | N/A | N/A | N/A | N/A | N/A | N/A | N/A | N/A |
| chr17 | 2082631 | 2082651 | Tier3_OT135_1_Intron | 0,16 | + | UNCOVERseqTier3 | 1 | true | false | false | false | false | N/A | N/A | N/A | N/A | N/A | N/A | N/A | N/A | N/A | N/A | N/A |
| chr1 | 2624197 | 2624217 | Tier3_OT136_1_Intron | 0,16 | - | UNCOVERseqTier3 | 1 | true | false | false | false | false | N/A | N/A | N/A | N/A | N/A | N/A | N/A | N/A | N/A | N/A | N/A |
| chr17 | 7226857 | 7226877 | Tier3_OT138_1_Intron | 0,136 | + | UNCOVERseqTier3 | 1 | true | false | false | false | false | N/A | N/A | N/A | N/A | N/A | N/A | N/A | N/A | N/A | N/A | N/A |
| chr22 | 24416909 | 24416929 | Tier3_OT139_1_Exon | 0,134 | - | UNCOVERseqTier3 | 1 | true | false | false | false | false | N/A | N/A | N/A | N/A | N/A | N/A | N/A | N/A | N/A | N/A | N/A |
| chr10 | 128071619 | 1,28E+08 | Tier3_OT140_1_Exon | 0,126 | - | UNCOVERseqTier3 | 1 | true | false | false | false | false | N/A | N/A | N/A | N/A | N/A | N/A | N/A | N/A | N/A | N/A | N/A |
| chr10 | 117504330 | 1,18E+08 | Tier3_OT141_1_Other | 0,122 | + | UNCOVERseqTier3 | 1 | true | false | false | false | false | N/A | N/A | N/A | N/A | N/A | N/A | N/A | N/A | N/A | N/A | N/A |
| chr10 | 55578869 | 55578889 | Tier3_OT142_1_Intron | 0,119 | - | UNCOVERseqTier3 | 1 | true | false | false | false | false | N/A | N/A | N/A | N/A | N/A | N/A | N/A | N/A | N/A | N/A | N/A |
| chrX | 48516949 | 48516969 | Tier3_OT143_1_Intron | 0,117 | + | UNCOVERseqTier3 | 1 | true | false | false | false | false | N/A | N/A | N/A | N/A | N/A | N/A | N/A | N/A | N/A | N/A | N/A |
| chr4 | 97633904 | 97633924 | Tier3_OT144_1_Intron | 0,117 | - | UNCOVERseqTier3 | 3 | true | false | false | false | false | N/A | N/A | N/A | N/A | N/A | N/A | N/A | N/A | N/A | N/A | N/A |
| chr15 | 42548847 | 42548867 | Tier3_OT145_1_Exon | 0,115 | - | UNCOVERseqTier3 | 1 | true | false | false | false | false | N/A | N/A | N/A | N/A | N/A | N/A | N/A | N/A | N/A | N/A | N/A |
| chr7 | 158518642 | 1,59E+08 | Tier3_OT147_1_Intron | 0,087 | + | UNCOVERseqTier3 | 1 | true | false | false | false | false | N/A | N/A | N/A | N/A | N/A | N/A | N/A | N/A | N/A | N/A | N/A |
| chr12 | 508980 | 509000 | Tier3_OT148_1_Intron | 0,085 | - | UNCOVERseqTier3 | 1 | true | false | false | false | false | N/A | N/A | N/A | N/A | N/A | N/A | N/A | N/A | N/A | N/A | N/A |
| chr3 | 64444755 | 64444776 | Tier3_OT149_1_Other | 0,079 | - | UNCOVERseqTier3 | 1 | true | false | false | false | false | N/A | N/A | N/A | N/A | N/A | N/A | N/A | N/A | N/A | N/A | N/A |
| chr19 | 13139600 | 13139620 | Tier3_OT151_1_Exon | 0,079 | + | UNCOVERseqTier3 | 1 | true | false | false | false | false | N/A | N/A | N/A | N/A | N/A | N/A | N/A | N/A | N/A | N/A | N/A |
| chr19 | 3668163 | 3668183 | Tier3_OT153_1_Intron | 0,077 | + | UNCOVERseqTier3 | 1 | true | false | false | true | false | A>G | 0,706819 | 0,912793 | Enhanced | 0,3 | 0,125 | 0,013 | 0,193 | 0,144 | N/A | N/A |
| chr15 | 100513286 | 1,01E+08 | Tier3_OT154_1_Intron | 0,077 | - | UNCOVERseqTier3 | 1 | true | false | false | false | false | N/A | N/A | N/A | N/A | N/A | N/A | N/A | N/A | N/A | N/A | N/A |
| chr4 | 338202 | 338222 | Tier3_OT155_1_Intron | 0,072 | + | UNCOVERseqTier3 | 1 | true | false | false | false | false | N/A | N/A | N/A | N/A | N/A | N/A | N/A | N/A | N/A | N/A | N/A |
| chr18 | 4095289 | 4095309 | Tier3_OT156_1_Intron | 0,071 | + | UNCOVERseqTier3 | 1 | true | false | false | false | false | N/A | N/A | N/A | N/A | N/A | N/A | N/A | N/A | N/A | N/A | N/A |
| chr13 | 99778629 | 99778648 | Tier3_OT157_1_Intron | 0,056 | + | UNCOVERseqTier3 | 1 | true | false | false | false | false | N/A | N/A | N/A | N/A | N/A | N/A | N/A | N/A | N/A | N/A | N/A |
| chr2 | 209690836 | 2,1E+08 | Tier3_OT158_1_Exon | 0,054 | - | UNCOVERseqTier3 | 1 | true | false | false | false | false | N/A | N/A | N/A | N/A | N/A | N/A | N/A | N/A | N/A | N/A | N/A |
| chr3 | 47008440 | 47008460 | Tier3_OT159_1_Exon | 0,047 | - | UNCOVERseqTier3 | 1 | true | false | false | false | false | N/A | N/A | N/A | N/A | N/A | N/A | N/A | N/A | N/A | N/A | N/A |
| chr1 | 21298140 | 21298160 | Tier3_OT161_1_Exon | 0,043 | + | UNCOVERseqTier3 | 1 | true | false | false | false | false | N/A | N/A | N/A | N/A | N/A | N/A | N/A | N/A | N/A | N/A | N/A |
| chr16 | 81826763 | 81826783 | Tier3_OT162_1_Intron | 0,042 | + | UNCOVERseqTier3 | 1 | true | false | false | false | false | N/A | N/A | N/A | N/A | N/A | N/A | N/A | N/A | N/A | N/A | N/A |
| chr8 | 50728188 | 50728208 | Tier3_OT163_1_Intron | 0,034 | - | UNCOVERseqTier3 | 1 | true | false | false | false | false | N/A | N/A | N/A | N/A | N/A | N/A | N/A | N/A | N/A | N/A | N/A |
| chr9 | 17290589 | 17290609 | Tier3_OT164_1_Intron | 0,032 | + | UNCOVERseqTier3 | 3 | true | false | false | false | false | N/A | N/A | N/A | N/A | N/A | N/A | N/A | N/A | N/A | N/A | N/A |
| chr10 | 32642350 | 32642370 | Tier3_OT165_1_Intron | 0,032 | + | UNCOVERseqTier3 | 1 | true | false | false | true | false | C>T | 0,351093 | 0,685925 | Enhanced | 0,045 | 0,18 | 0,252 | 0,167 | 0,169 | N/A | N/A |
| chrX | 138778729 | 1,39E+08 | Tier3_OT166_1_Intron | 0,029 | - | UNCOVERseqTier3 | 3 | true | false | false | false | false | N/A | N/A | N/A | N/A | N/A | N/A | N/A | N/A | N/A | N/A | N/A |
| chrX | 14809617 | 14809637 | Tier3_OT167_1_Other | 0,029 | - | UNCOVERseqTier3 | 1 | true | false | false | false | false | N/A | N/A | N/A | N/A | N/A | N/A | N/A | N/A | N/A | N/A | N/A |
| chr1 | 244994673 | 2,45E+08 | Tier3_OT168_1_Intron | 0,019 | + | UNCOVERseqTier3 | 1 | true | false | false | false | false | N/A | N/A | N/A | N/A | N/A | N/A | N/A | N/A | N/A | N/A | N/A |
| chr17 | 8236199 | 8236218 | plex329OT144 | 0 | + | NotUNCOVERseq_Charite | 1 | false | true | false | false | false | N/A | N/A | N/A | N/A | N/A | N/A | N/A | N/A | N/A | N/A | N/A |
| chr11 | 10674973 | 10674991 | plex329OT18 | 0 | - | NotUNCOVERseq_Charite | 1 | false | true | false | false | false | N/A | N/A | N/A | N/A | N/A | N/A | N/A | N/A | N/A | N/A | N/A |
| chr7 | 124863505 | 1,25E+08 | plex329OT269 | 0 | + | NotUNCOVERseq_Charite | 1 | false | true | false | false | false | N/A | N/A | N/A | N/A | N/A | N/A | N/A | N/A | N/A | N/A | N/A |
| chr7 | 44796771 | 44796787 | plex329OT280 | 0 | - | NotUNCOVERseq_Charite | 1 | false | true | false | false | false | N/A | N/A | N/A | N/A | N/A | N/A | N/A | N/A | N/A | N/A | N/A |

| | | | | | | | | | | | | | | | | | | | | | | | |
|---|---|---|---|---|---|---|---|---|---|---|---|---|---|---|---|---|---|---|---|---|---|---|---|
| chr8 | 144679271 | 1,45E+08 | plex329OT291 | 0 | + | NotUNCOVERseq_Charite | 1 | false | true | false | false | false | N/A | N/A | N/A | N/A | N/A | N/A | N/A | N/A | N/A | N/A | N/A |
| chrX | 55025838 | 55025848 | plex329OT328 | 0 | - | NotUNCOVERseq_Charite | 1 | false | true | false | false | false | N/A | N/A | N/A | N/A | N/A | N/A | N/A | N/A | N/A | N/A | N/A |
| chr14 | 22547566 | 22547582 | plex329OT82 | 0 | + | NotUNCOVERseq_Charite | 1 | false | true | false | false | false | N/A | N/A | N/A | N/A | N/A | N/A | N/A | N/A | N/A | N/A | N/A |
| chr14 | 22924021 | 22924035 | plex329OT83 | 0 | + | NotUNCOVERseq_Charite | 1 | false | true | false | false | false | N/A | N/A | N/A | N/A | N/A | N/A | N/A | N/A | N/A | N/A | N/A |
| chr2 | 133715387 | 1,34E+08 | plex329OT181 | 0 | - | EmpiricalOverlap | 1 | false | true | true | false | false | N/A | N/A | N/A | N/A | N/A | N/A | N/A | N/A | N/A | N/A | N/A |
| chr10 | 97283088 | 97283107 | 73E96DB9624844AZ0Z | 0 | + | EmpiricalOverlap | 3 | false | true | true | false | false | N/A | N/A | N/A | N/A | N/A | N/A | N/A | N/A | N/A | N/A | N/A |
| chr6 | 63211592 | 63211611 | plex329OT265 | 0 | + | NotUNCOVERseq_Charite | 1 | false | true | false | true | false | N/A | N/A | N/A | N/A | N/A | N/A | N/A | N/A | N/A | N/A | N/A |
| chr15 | 25798315 | 25798333 | plex329OT95 | 0 | + | NotUNCOVERseq_Charite | 1 | false | true | false | true | false | N/A | N/A | N/A | N/A | N/A | N/A | N/A | N/A | N/A | N/A | N/A |
| chr2 | 3054245 | 3054263 | plex329OT200 | 0 | + | NotUNCOVERseq_Charite | 1 | false | true | false | true | false | N/A | N/A | N/A | N/A | N/A | N/A | N/A | N/A | N/A | N/A | N/A |
| chr6 | 41771841 | 41771861 | plex329OT261 | 0 | + | NotUNCOVERseq_Charite | 1 | false | true | false | true | false | N/A | N/A | N/A | N/A | N/A | N/A | N/A | N/A | N/A | N/A | N/A |
| chr14 | 105529242 | 1,06E+08 | plex329OT80 | 0 | + | NotUNCOVERseq_Charite | 1 | false | true | false | true | false | N/A | N/A | N/A | N/A | N/A | N/A | N/A | N/A | N/A | N/A | N/A |
| chr3 | 41199180 | 41199196 | plex329OT218 | 0 | + | NotUNCOVERseq_Charite | 1 | false | true | false | true | false | N/A | N/A | N/A | N/A | N/A | N/A | N/A | N/A | N/A | N/A | N/A |
| chr9 | 87886273 | 87886292 | plex329OT314 | 0 | + | EmpiricalOverlap | 1 | false | true | true | true | false | N/A | N/A | N/A | N/A | N/A | N/A | N/A | N/A | N/A | N/A | N/A |
| chr9 | 91039968 | 91039986 | plex329OT315 | 0 | - | NotUNCOVERseq_Charite | 1 | false | true | false | true | false | N/A | N/A | N/A | N/A | N/A | N/A | N/A | N/A | N/A | N/A | N/A |
| chr5 | 62678505 | 62678522 | plex18OT14 | 0 | + | NotUNCOVERseq_Charite | 2 | false | true | false | true | false | N/A | N/A | N/A | N/A | N/A | N/A | N/A | N/A | N/A | N/A | N/A |
| chr6 | 48363518 | 48363536 | plex329OT263 | 0 | - | NotUNCOVERseq_Charite | 3 | false | true | false | true | false | N/A | N/A | N/A | N/A | N/A | N/A | N/A | N/A | N/A | N/A | N/A |
| chr2 | 42864363 | 42864381 | plex329OT201 | 0 | + | NotUNCOVERseq_Charite | 1 | false | true | false | true | false | C>T | 0,869679 | 0,731168 | Reduced | 0,012 | 0,00157 | 0 | 0,00001 | 0,00062 | N/A | N/A |
| chr9 | 135296676 | 1,35E+08 | plex329OT306 | 0 | + | NotUNCOVERseq_Charite | 1 | false | true | false | true | false | N/A | N/A | N/A | N/A | N/A | N/A | N/A | N/A | N/A | N/A | N/A |
| chr15 | 84200239 | 84200257 | plex18OT2 | 0 | - | NotUNCOVERseq_Charite | 3 | false | true | false | true | false | N/A | N/A | N/A | N/A | N/A | N/A | N/A | N/A | N/A | N/A | N/A |
| chr15 | 85204837 | 85204855 | plex329OT106 | 0 | - | NotUNCOVERseq_Charite | 3 | false | true | false | true | false | N/A | N/A | N/A | N/A | N/A | N/A | N/A | N/A | N/A | N/A | N/A |
| chr6 | 52062692 | 52062711 | plex329OT264 | 0 | + | NotUNCOVERseq_Charite | 1 | false | true | false | true | false | N/A | N/A | N/A | N/A | N/A | N/A | N/A | N/A | N/A | N/A | N/A |
| chr15 | 84473305 | 84473323 | plex329OT105 | 0 | + | NotUNCOVERseq_Charite | 3 | false | true | false | true | false | C>T | 0,808142 | 0,759549 | Reduced | 0,012 | 0,032 | 0 | 0,085 | 0,00909 | N/A | N/A |
| chr9 | 133310440 | 1,33E+08 | plex329OT304 | 0 | - | NotUNCOVERseq_Charite | 1 | false | true | false | true | false | N/A | N/A | N/A | N/A | N/A | N/A | N/A | N/A | N/A | N/A | N/A |
| chr7 | 150706666 | 1,51E+08 | plex329OT272 | 0 | - | NotUNCOVERseq_Charite | 2 | false | true | false | true | false | N/A | N/A | N/A | N/A | N/A | N/A | N/A | N/A | N/A | N/A | N/A |
| chr13 | 112680608 | 1,13E+08 | plex329OT63 | 0 | - | NotUNCOVERseq_Charite | 1 | false | true | false | true | false | N/A | N/A | N/A | N/A | N/A | N/A | N/A | N/A | N/A | N/A | N/A |
| chr5 | 181189224 | 1,81E+08 | plex329OT244 | 0 | + | NotUNCOVERseq_Charite | 1 | false | true | false | true | false | N/A | N/A | N/A | N/A | N/A | N/A | N/A | N/A | N/A | N/A | N/A |
| chr1 | 3690313 | 3690332 | plex329OT72 | 0 | - | NotUNCOVERseq_Charite | 1 | false | true | false | true | false | N/A | N/A | N/A | N/A | N/A | N/A | N/A | N/A | N/A | N/A | N/A |
| chr4 | 11235654 | 11235673 | Insilico1 | 0 | - | InSilicoCriteria | 3 | false | false | false | true | false | G>A | 0,351093 | 0,716391 | Enhanced | 0,127 | 0,201 | 0,257 | 0,047 | 0,036 | N/A | N/A |
| chr8 | 56122471 | 56122490 | Insilico2 | 0 | + | InSilicoCriteria | 1 | false | false | false | true | false | T>C | 0,61581 | 0,920352 | Enhanced | 0,043 | 0,176 | 0,00058 | 0,222 | 0,078 | N/A | N/A |
| chr8 | 141809999 | 1,42E+08 | Insilico3 | 0 | + | InSilicoCriteria | 1 | false | false | false | true | false | A>G | 0,549894 | 0,787042 | Enhanced | 0,119 | 0,018 | 0 | 0,00081 | 0,00041 | N/A | N/A |
| chr16 | 17348551 | 17348571 | Insilico4 | 0 | + | InSilicoCriteria | 1 | false | false | false | true | false | G>C | 0,157232 | 0,825874 | Enhanced | 0,075 | 0,00937 | 0 | 0,00047 | 0,00872 | N/A | N/A |
| chr8 | 98589678 | 98589697 | Insilico5 | 0 | + | InSilicoCriteria | 2 | false | false | false | true | false | C>A | 0,088301 | 0,645627 | Enhanced | 0,011 | 0,016 | 0,00019 | 0,046 | 0,06 | N/A | N/A |
| chr20 | 32233281 | 32233299 | Insilico6 | 0 | + | InSilicoCriteria | 1 | false | false | false | true | false | G>A | 0,107549 | 0,746832 | Enhanced | 0,053 | 0,00916 | 0 | 0,00028 | 0,00041 | N/A | N/A |
| chr16 | 89162639 | 89162658 | Insilico7 | 0 | - | InSilicoCriteria | 1 | false | false | false | true | false | A>G | 0,733526 | 0,733526 | Neutral | 0,032 | 0,00242 | 0 | 0,0001 | 0 | N/A | N/A |
| chr12 | 129801128 | 1,3E+08 | Insilico8 | 0 | - | InSilicoCriteria | 1 | false | false | false | true | false | G>C | N/A | 0,54094 | Enabled | 0,027 | 0,00183 | 0 | 0,00004 | 0,00062 | N/A | N/A |
| chr7 | 47724487 | 47724506 | Insilico9 | 0 | - | InSilicoCriteria | 1 | false | false | false | true | false | T>C | 0,38898 | 0,777321 | Enhanced | 0,027 | 0,00262 | 0 | 0,00006 | 0,00021 | N/A | N/A |
| chr11 | 89426665 | 89426684 | Insilico10 | 0 | - | InSilicoCriteria | 1 | false | false | false | true | false | G>A | 0,369169 | 0,700657 | Enhanced | 0,0013 | 0,025 | 0,013 | 0,00007 | 0,00166 | N/A | N/A |
| chr1 | 204685459 | 2,05E+08 | Insilico11 | 0 | + | InSilicoCriteria | 1 | false | false | false | true | false | N/A | 0,530963 | N/A | N/A | N/A | N/A | N/A | N/A | N/A | N/A | N/A |
| chr3 | 138348884 | 1,38E+08 | Insilico12 | 0 | - | InSilicoCriteria | 1 | false | false | false | true | false | N/A | 0,756767 | N/A | N/A | N/A | N/A | N/A | N/A | N/A | N/A | N/A |
| chr7 | 122306040 | 1,22E+08 | Insilico13 | 0 | + | InSilicoCriteria | 1 | false | false | false | true | false | N/A | 0,683932 | N/A | N/A | N/A | N/A | N/A | N/A | N/A | N/A | N/A |
| chr15 | 82472843 | 82472863 | Insilico14 | 0 | - | InSilicoCriteria | 1 | false | false | false | true | false | N/A | 0,814302 | N/A | N/A | N/A | N/A | N/A | N/A | N/A | N/A | N/A |
| chr17 | 74862650 | 74862667 | Insilico15 | 0 | + | InSilicoCriteria | 1 | false | false | false | true | false | N/A | 0,620545 | N/A | N/A | N/A | N/A | N/A | N/A | N/A | N/A | N/A |
| chr19 | 46405134 | 46405153 | Insilico16 | 0 | + | InSilicoCriteria | 1 | false | false | false | true | false | N/A | 0,743495 | N/A | N/A | N/A | N/A | N/A | N/A | N/A | N/A | N/A |
| chr20 | 23633810 | 23633828 | Insilico17 | 0 | + | InSilicoCriteria | 1 | false | false | false | true | false | N/A | 0,66162 | N/A | N/A | N/A | N/A | N/A | N/A | N/A | N/A | N/A |
| chr12 | 57097726 | 57097749 | CASTseq1 | 0 | - | CASTseqCriteria | 1 | false | false | false | false | true | N/A | N/A | N/A | N/A | N/A | N/A | N/A | N/A | N/A | 62 | 45 |
| chr5 | 138564795 | 1,39E+08 | CASTseq2 | 0 | - | CASTseqCriteria | 1 | false | false | false | false | true | N/A | N/A | N/A | N/A | N/A | N/A | N/A | N/A | N/A | 66 | 39 |
| chr19 | 19105438 | 19105461 | CASTseq3 | 0 | - | CASTseqCriteria | 1 | false | false | false | false | true | N/A | N/A | N/A | N/A | N/A | N/A | N/A | N/A | N/A | 26 | 21 |
| chr1 | 99355110 | 99355130 | CASTseq4 | 0 | + | CASTseqCriteria | 1 | false | false | false | false | true | N/A | N/A | N/A | N/A | N/A | N/A | N/A | N/A | N/A | 221 | 2 |
| chr9 | 131725484 | 1,32E+08 | CASTseq5 | 0 | + | CASTseqCriteria | 1 | false | false | false | false | true | N/A | N/A | N/A | N/A | N/A | N/A | N/A | N/A | N/A | 20 | 8 |
| chr1 | 74996950 | 74996975 | CASTseq6 | 0 | + | CASTseqCriteria | 1 | false | false | false | false | true | N/A | N/A | N/A | N/A | N/A | N/A | N/A | N/A | N/A | 22442 | 6 |
| chr4 | 74381251 | 74381273 | CASTseq7 | 0 | + | CASTseqCriteria | 1 | false | false | false | false | true | N/A | N/A | N/A | N/A | N/A | N/A | N/A | N/A | N/A | 7136 | 4 |
| chr17 | 69121408 | 69121429 | CASTseq8 | 0 | - | CASTseqCriteria | 1 | false | false | false | false | true | N/A | N/A | N/A | N/A | N/A | N/A | N/A | N/A | N/A | 14188 | 3 |
| chr6 | 100826377 | 1,01E+08 | CASTseq9 | 0 | + | CASTseqCriteria | 3 | false | false | false | false | true | N/A | N/A | N/A | N/A | N/A | N/A | N/A | N/A | N/A | 12546 | 2 |
| chr2 | 32575479 | 32575500 | CASTseq10 | 0 | - | CASTseqCriteria | 2 | false | false | false | false | true | N/A | N/A | N/A | N/A | N/A | N/A | N/A | N/A | N/A | 621 | 2 |
| chr5 | 37970288 | 37970310 | plex329OT248 | 0 | - | PreviousHit | 1 | false | false | false | true | false | N/A | N/A | N/A | N/A | N/A | N/A | N/A | N/A | N/A | N/A | N/A |
| chr17 | 37985648 | 37985672 | E0BDACADDEF44B0Z0Z | 0 | + | PreviousHit | 3 | false | false | false | true | false | N/A | N/A | N/A | N/A | N/A | N/A | N/A | N/A | N/A | N/A | N/A |
| chr11 | 70086588 | 70086611 | plex329OT35 | 0 | - | AnnotationDiscrepancy | 1 | false | false | false | true | false | N/A | N/A | N/A | N/A | N/A | N/A | N/A | N/A | N/A | N/A | N/A |

### Suppl. Table 6: Methods qualification of digital droplet PCR of FKBP12$^{KO}$-Treg

Results qualification for KO-efficacy read-outs
Assays 1-4
06.02. - 07.02.2025

| Operators | Donor 1 | Donor 1 | Donor 2 | Donor 1 | | | | | | | | | | | | |
|---|---|---|---|---|---|---|---|---|---|---|---|---|---|---|---|---|
| Date | 07.02.25 | 07.02.25 | 06.02.25 | 06.02.25 | A1 | A1 | A1 | A2 | A2 | A2 | A3 | A3 | A3 | A4 | A4 | A4 |
| Assay | A1 | A2 | A3 | A4 | 50 | 75 | 100 | 50 | 75 | 100 | 50 | 75 | 100 | 50 | 75 | 100 |
| 1 | 90,45 | 90,34 | 91,14 | 90,55 | 47,66 | 73,99 | 99,03 | 49,58 | 72,43 | 97,67 | 50,99 | 75,37 | 100,00 | 49,05 | 74,09 | 99,88 |
| 2 | 90,37 | 90,32 | 91,54 | 90,10 | 47,93 | 73,09 | 98,55 | 47,60 | 72,34 | 97,25 | 50,04 | 75,16 | 100,00 | 48,90 | 74,22 | 99,80 |
| 3 | 90,27 | 89,64 | 91,61 | 90,03 | | | | | | | | | | | | |
| 4 | 90,02 | 89,80 | 90,99 | 89,77 | | | | | | | | | | | | |
| 5 | 90,27 | 90,42 | 91,53 | 89,79 | | | | | | | | | | | | |
| 6 | 90,57 | 90,05 | 90,89 | 90,28 | | | | | | | | | | | | |
| 7 | 90,41 | 89,78 | 91,05 | 90,14 | | | | | | | | | | | | |
| 8 | 89,60 | 90,08 | 91,34 | 89,06 | | | | | | | | | | | | |
| Mean | 90,24 | 90,05 | 91,26 | 89,96 | | 73,54 | | | 72,39 | | | 75,27 | | | 74,15 | |
| standard deviation | 0,31 | 0,29 | 0,28 | 0,44 | | | | | | | | | | | | |
| coefficient of variation | 0,34 | 0,33 | 0,30 | 0,49 | | | | | | | | | | | | |
| Reference Sanger-seq. | 93,00 | 91,00 | 99,00 | 92,00 | | | | | | | | | | | | |

100

| | Intra-A | Inter-A I | Inter-O I | Ges. | Inter-O II | Inter-A II | A1 | A1 | A1 | A2 | A2 | A2 | A3 | A3 | A3 | A4 | A4 | A4 | A1-A4 | Sanger Seq. |
|---|---|---|---|---|---|---|---|---|---|---|---|---|---|---|---|---|---|---|---|---|
| | A1, A2 | A1, A4 | A3, A4 | A1, A2, A3, A4 | A1, A3 | A2, A3 | 50 | 75 | 100 | 50 | 75 | 100 | 50 | 75 | 100 | 50 | 75 | 100 | 75 | |
| Mean | 90,15 | 90,10 | 90,61 | 90,38 | 90,75 | 90,66 | 47,80 | 73,54 | 98,79 | 48,59 | 72,39 | 97,46 | 50,52 | 75,27 | 100,00 | 48,98 | 74,15 | 99,84 | 73,84 | 93,75 |
| standard deviation | 0,31 | 0,40 | 0,76 | 0,62 | 0,60 | 0,68 | | | | | | | | | | | | | | |
| coefficient of variation | 0,34 | 0,44 | 0,84 | 0,68 | 0,66 | 0,75 | | | | | | | | | | | | | | |
| % of deviation Assay vs. Reference | | | | | | | | | | | | | | | | | | | | -3,59 |

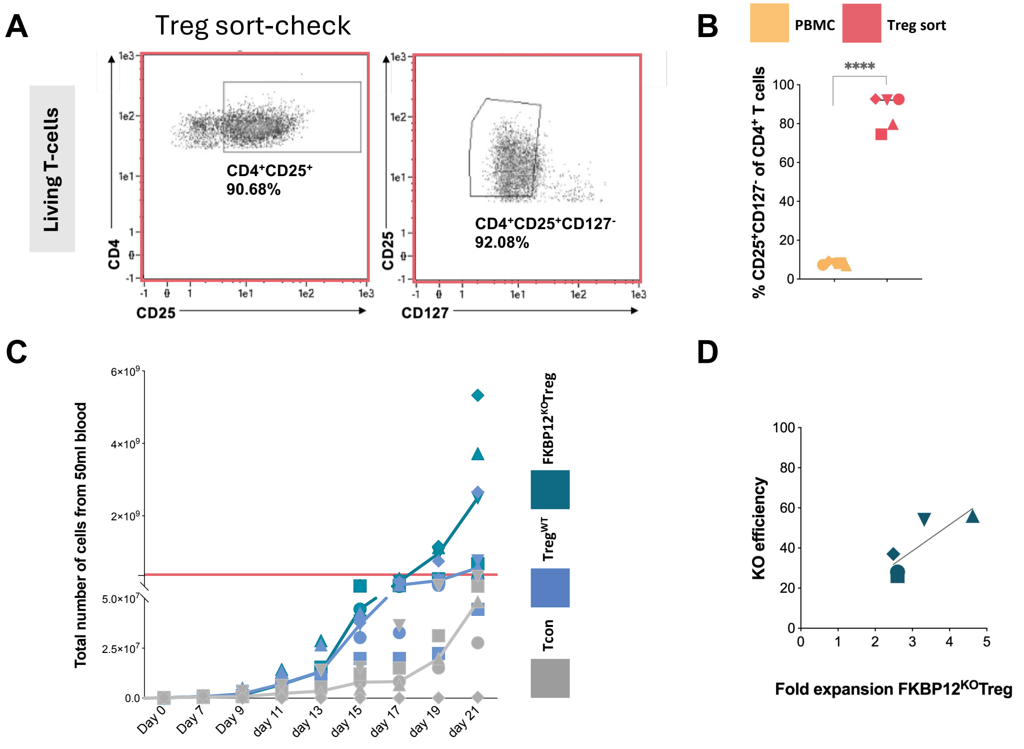


**Suppl. Figure 1: Tyto sort purity and Treg expansion.**

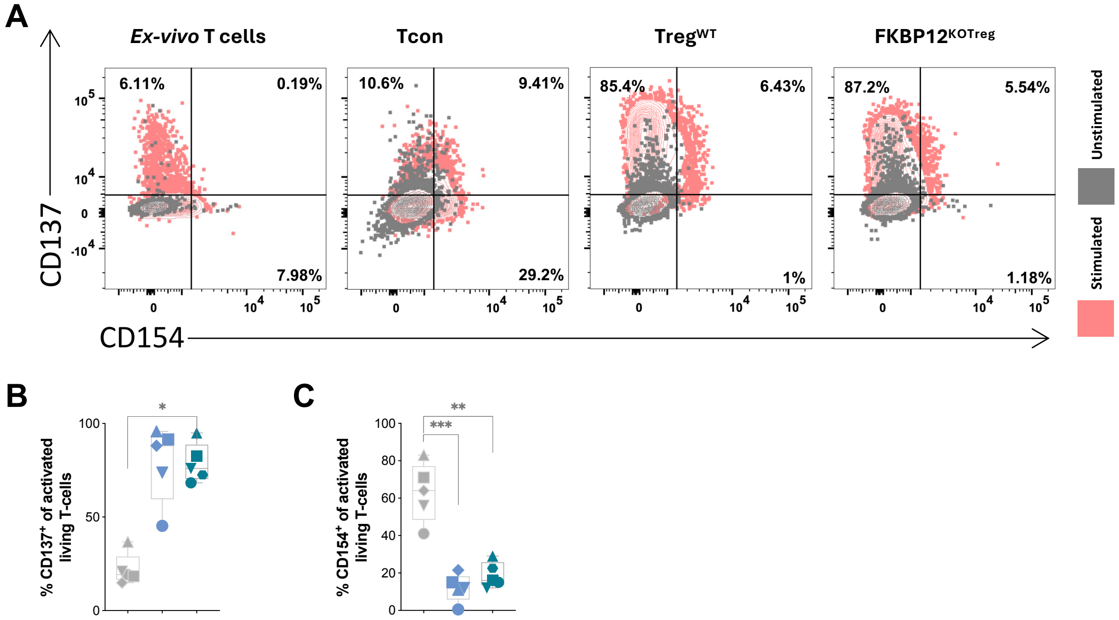


**Suppl. Figure 2: *FKBP12$^{KO}$-Treg activation is comparable with Treg$^{WT}$.***

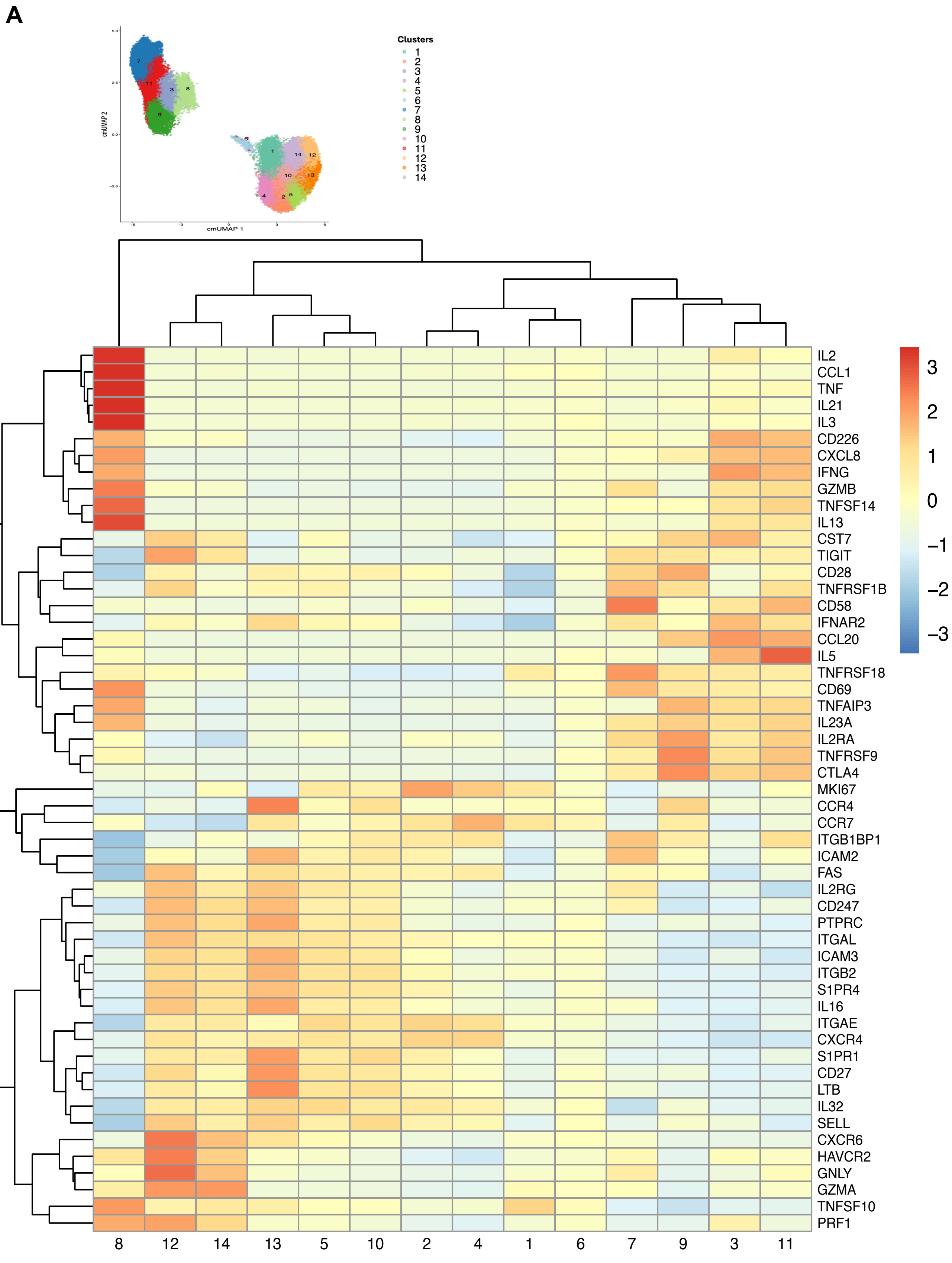


**Suppl. Figure 3:** ***Cellular Indexing of Transcriptomes and Epitope sequencing.***

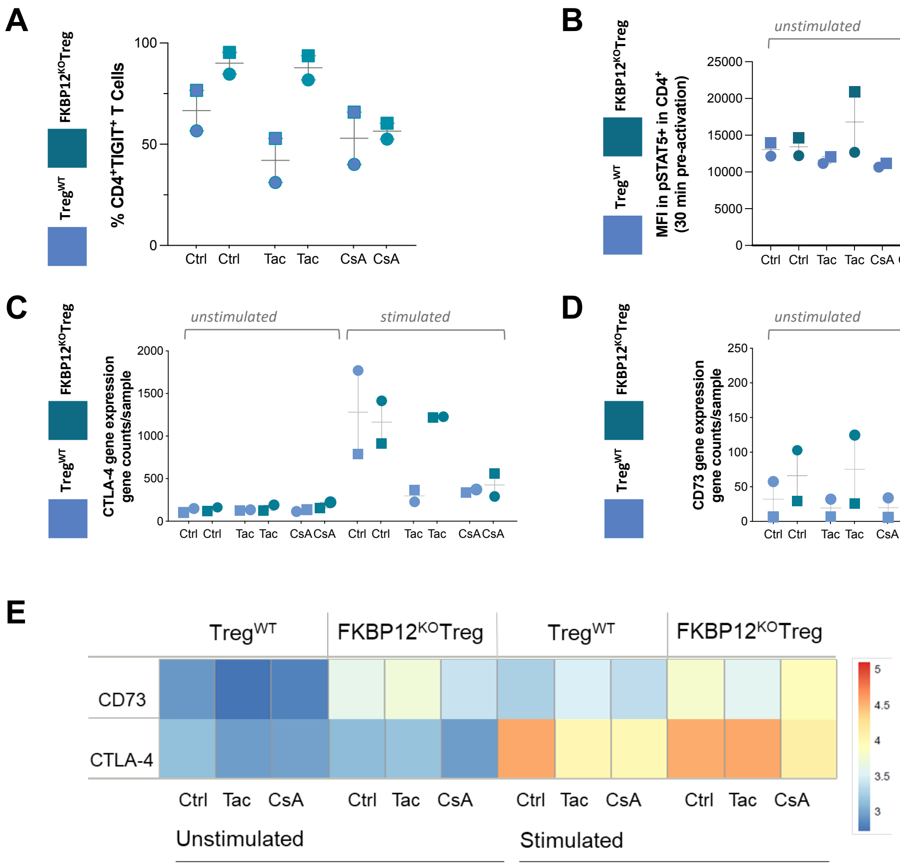


**Suppl. Figure 4: Beneficial functionality of FKBP12$^{KO}$-Treg in comparison to Treg$^{WT}$.**

**Supplementary table legends**

**Suppl. Table 1: List of general antibodies**

Summary of general Treg specific antibodies giving information on antibody, fluorescence label and clone.

**Suppl. Table 2: List of materials and instruments**

Summary of materials, supplements and instruments used giving information on material, concentration and company.

**Suppl. Table 3: List of CITEseq antibodies**

Summary of CITEseq antibodies giving information on reagent, clone, source and identifier.

**Suppl. Table 4: Database comparison of differentially expressed CITEseq genes in FKBP12$^{KO}$-Treg and Treg$^{WT}$**

Summary of significantly regulated genes pre-selected from volcano plots comparing expression levels in FKBP12$^{KO}$-Treg and Treg$^{WT}$. Those genes further underwent DAVID database analysis and resulting gene dependent pathways are displayed.

**Suppl. Table 5: Off-target nomination sites**

Summary of off-target nomination sites selected with the following criteria: 1) All sites that had overlap between one or more empirical method (abnoba-seq/guide-seq/UNCOVERseq); 2) All UNCOVERseq sites with Tiers 1 to 3 (our highest priorities); 3) All GUIDE-seq sites; 4) All Abnoba-seq sites; 5) All in silico sites with lev dist < 3 (reference or population >1%); 6) All in silico sites with lev dist <4 + an exonic region annotated (reference or population > 1%)

**Suppl. Table 6: Methods qualification of digital droplet PCR of FKBP12$^{KO}$-Treg**

Qualification results for KO-efficacy read-outs for in total 4 assays indicating operator, date, assay, mean and standard deviation and reference Sanger-sequencing.

**Supplementary figure legends**

**Suppl. Figure 1: Tyto sort purity and Treg expansion.** (A) Gating strategy for Tyto sort showing CD4, CD25 and CD127 gating. Remaining figures show data from individual donors by different symbols: (B) Sort purity shown as the percentage of $CD4^+CD25^+CD127^-$ T-cells pre- (yellow) and post sorting (red) p<0.0001(****). (C) Visualization of the total number of expanding cells from day 0 to day 21. Tcon (grey), $Treg^{WT}$ (blue), $FKBP12^{KO}$-Treg (teal), red line marks cut-off for Treg amount in clinical application (D) The correlation between knock-out efficiency and fold expansion of $FKBP12^{KO}$-Treg (teal).

**Suppl. Figure 2: *$FKBP12^{KO}$-Treg activation is comparable with $Treg^{WT}$.*** (A) Representative dot plots for CD137 and CD154 expression in *ex-vivo* T-cells, Tcon, $Treg^{WT}$ and $FKBP12^{KO}$-Treg on day 21. Unstimulated controls (grey) and stimulated conditions (red). Remaining figures show individual donors by different symbols. Tcon (grey), $Treg^{WT}$ (blue), $FKBP12^{KO}$-Treg (teal). (B) Percentage of $CD137^+$ among the living $CD4^+$ T-cells on day 21 in Tcon, $Treg^{WT}$ and $FKBP12^{KO}$-Treg, p<0.05(*). (C) Percentage of $CD154^+$ among the living T-cells on day 21 in Tcon, $Treg^{WT}$ and $FKBP12^{KO}$-Treg. For (B) and (C) one-way ANOVA (and nonparametric), p<0.05(*), p<0.01(**), p<0.001(***), mean +/- SD n=5.

**Suppl. Figure 3: *Cellular Indexing of Transcriptomes and Epitope sequencing.*** Detailed heatmap of RNA expression of T cell-associated genes within different clusters. Cluster distribution is shown for all 14 clusters and differential gene expression within each cluster (1-14) is highlighted within the heatmap.

**Suppl. Figure 4: Beneficial functionality of $FKBP12^{KO}$-Treg in comparison to $Treg^{WT}$.**

Each symbol represents data from an individual donor. Tcon (white), $Treg^{WT}$ (gray), $FKBP12^{KO}$-Treg (teal) n=2. For all figures conditions tacrolimus (Tac), cyclosporin A (CsA) and medium control (Ctrl) are shown. (A) Tigit expression among the living $CD4^+$

T-cells on day 21 in Treg$^{WT}$ and FKBP12$^{KO}$-Treg. (B) pSTAT5 MFI after 30 minutes stimulation by IL2 on day 21 in Treg$^{WT}$ and FKBP12$^{KO}$-Treg compared to unstimulated control. (C-D) CTLA-4 and CD73 gene expression levels are shown for Treg$^{WT}$ and FKBP12$^{KO}$-Treg in unstimulated and stimulated states (6h PMA/Ionomycin activation) on day 28. Mean values of 2 technical replicates are displayed from a total of 2 donors. (E) Heatmap of surface CD73 and CTLA-4 is shown for Treg$^{WT}$ vs. FKBP12$^{KO}$-Treg in unstimulated and stimulated states (6h PMA/Ionomycin activation). Surface expression levels are shown based on staining with CITE-seq antibody-derived tags included in the single cell data set.
